# Identification of Human Pathways Acting on Nuclear Non-Coding RNAs Using the Mirror Forward Genetic Approach

**DOI:** 10.1101/2024.09.26.615073

**Authors:** Rui Che, Monireh Panah, Bhoomi Mirani, Krista Knowles, Anastacia Ostapovich, Debarati Majumdar, Xiaotong Chen, Joseph DeSimone, William White, Megan Noonan, Hong Luo, Andrei Alexandrov

## Abstract

Despite critical roles in diseases, human pathways acting on strictly nuclear non-coding RNAs have been refractory to forward genetics. To enable their forward genetic discovery, we developed a single-cell approach that “Mirrors” activities of nuclear pathways with cytoplasmic fluorescence. Application of Mirror to two nuclear pathways targeting MALAT1’s 3′ end, the pathway of its maturation and the other, the degradation pathway blocked by the triple-helical Element for Nuclear Expression (ENE), identified nearly all components of three complexes: Ribonuclease P and the RNA Exosome, including nuclear DIS3, EXOSC10, and C1D, as well as the Nuclear Exosome Targeting (NEXT) complex. Additionally, Mirror identified DEAD-box helicase DDX59 associated with the genetic disorder Oral-Facial-Digital syndrome (OFD), yet lacking known substrates or roles in nuclear RNA degradation. Knockout of DDX59 exhibits stabilization of the full-length MALAT1 with a stability-compromised ENE and increases levels of 3′-extended forms of small nuclear RNAs. It also exhibits extensive retention of minor introns, including in OFD-associated genes, suggesting a mechanism for DDX59 association with OFD. Mirror efficiently identifies pathways acting on strictly nuclear non-coding RNAs, including essential and indirectly-acting components, and, as a result, uncovers unexpected links to human disease.

## INTRODUCTION

Nuclear-localized long non-coding RNAs (lncRNAs) carrying non-canonical 3′-ends with structured triple-helical Elements for Nuclear Expression (ENEs) (**Fig. 1a**)^1–4^ play critical roles in a variety of human diseases, including cancers^5,6^ as well as developmental^7^ and viral disorders^8,9^. Overexpression of the abundant, stable, vertebrate-specific, 5′-capped nuclear lncRNA Metastasis Associated Lung Adenocarcinoma Transcript 1 (MALAT1)^1,2,10,11^ promotes tumor growth by increasing cell proliferation, invasion, and metastasis. It correlates with poor survival in such prevalent cancers as lung, pancreatic, cervical, colorectal, and others^12–17^. The 3′ end of the nuclear lncRNA MALAT1 is not polyadenylated; instead, it is formed by RNase P cleavage (**Fig. 1b**) upstream of a tRNA-like RNA structure, mascRNA^18,19^. Since downregulation of MALAT1 RNA levels decreases tumor growth and metastasis and promotes cell differentiation^20,21^, this lncRNA represents a promising target for cancer treatment^5,11,21–25^. The ENE-containing human multiple endocrine neoplasia lncRNA^4^ (MEN-β, also called NEAT1_2), which is a component of nuclear paraspeckles, has also been implicated in a variety of cancers and diseases^6,26–29^.

**Figure 1.**
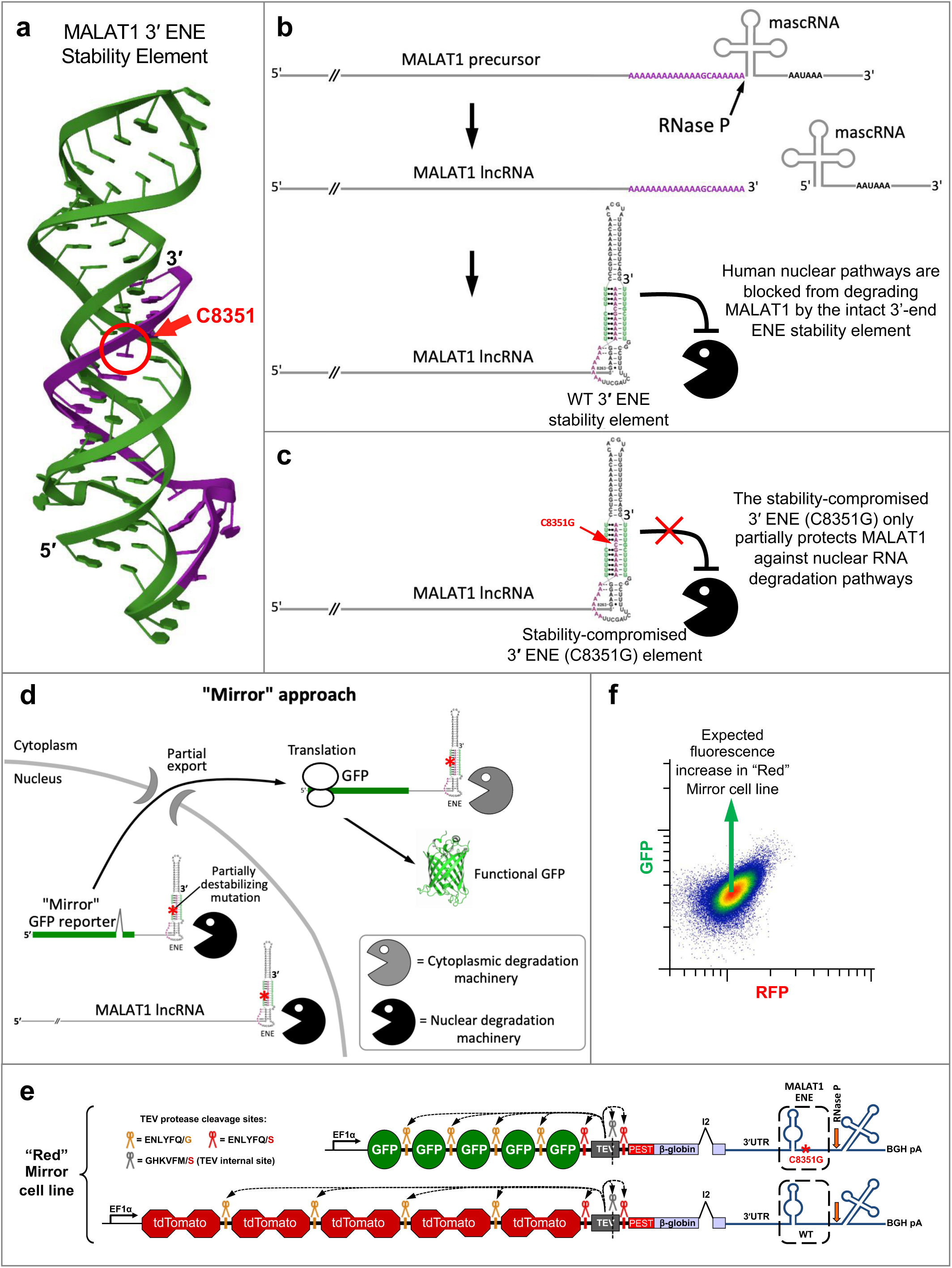
Schematics of the pathways acting on the 3′ end of MALAT1 and the Mirror approach to identify them. **a.** Crystal structure (PDB ID: 4PLX)^1,19^ of MALAT1 triple-helical ENE stability element that protects the 3′ end of MALAT1 against nuclear degradation pathways. Position of the ENE-destabilizing mutation C8351G^18^ used in this study is indicated by the red arrow. The image is made using Mol* Viewer^68^. **b.** Schematics of the MALAT1 3′-end formation by RNase P cleavage of the tRNA-like mascRNA structure and formation of the triple-helical ENE structure^1,18,19^ that protects the 3′ end of MALAT1 against nuclear degradation pathways. **c.** The stability-compromised ENE(C8351G) structure provides only partial protection against nuclear degradation pathways. **d.** Schematics of the Mirror concept for identifying nuclear pathways acting on nuclear long non-coding lncRNAs using reporters utilizing cytoplasmic fluorescence. **e.** The “Red” Mirror reporter cell line for simultaneous identification of two nuclear pathways acting on the MALAT1 3′ end: the first, a degradation pathway targeting the 3′ end of MALAT1 but prevented from degrading MALAT1 by its intact triple-helical ENE; and the second, the pathway required for MALAT1 3′-end processing. Major sensitivity-enhancing features of the Fireworks fluorescence amplification system employed by Mirror are described^35^. The 3′ end of the MALAT1 ENE Mirror reporters is formed by RNase P cleavage upstream of mascRNA, as indicated by the orange arrow. Steady-state levels of the MALAT1 ENE Mirror reporters depend on the (1) integrity of their 3′-end triple-helical ENE stability element, which is inserted after the β-globin 3′ UTR and can be disrupted by a single nucleotide substitution C8351G (shown in red). In addition to the “Red” Mirror cell line shown here, an orthogonal “Green” reporter cell line (shown schematically in **Figs. S1a** and **S2**) provides controls for cell line- and fluorescent protein-specific effects during genome-wide forward genetic screening and candidate gene validation. **f.** Anticipated effects of inhibiting pathways of nuclear degradation and processing on the fluorescence of “Red” Mirror cell line.

For each of these lncRNAs, a structured triple-helical ENE element (**Fig. 1a**) has been shown to protect the 3′ end from nuclear RNA degradation machinery (**Fig. 1b**) ^1–3,30^. This 3′ end-acting nuclear degradation machinery is efficient, as the lncRNAs’ half-life and steady-state levels critically depend on the protective function of the intact ENE structure^1,2^: a single-nucleotide destabilizing substitution C8351G within the MALAT1 ENE (**Fig. 1c**) results in a 20-fold reduction in MALAT1 levels, whereas the deletion of the entire 80-nucleotide ENE element leads to their dramatic 60-fold decrease^19^.

Despite the profound effect of nuclear pathways on steady-state levels of nuclear lncRNAs, and despite the major impact of these steady-state levels on human diseases, it is unknown what human nuclear degradation pathways are blocked from degrading nuclear lncRNAs by the 3′ triple-helical ENE elements. The reason such 3′-end targeting human pathways have not been interrogated using forward genetics is due to the strictly nuclear localization of the respective ENE-containing lncRNAs. Whereas forward genetic interrogation must use lncRNA abundance or processing efficiency as quantifiable screening readouts, their quantification can only be achieved in relatively low throughput. This stems from the fact that, unlike mRNAs, nuclear lncRNAs are not exported into cytoplasm and translated, precluding the use of translation-dependent reporter systems.

Such a technological barrier exists not only for the above-mentioned nuclear lncRNAs, but for nearly 30% of all human non-coding RNAs^31^ due to their nuclear localization. Due to the lack of robust genome-wide forward genetic approaches, most of our knowledge about human nuclear RNA pathways and their components, including the nuclear RNA exosome and its associated factors^32–34^ is derived from non-genetic approaches and studies in model organisms. This leads to a gap in knowledge about human-specific nuclear pathways and their regulation, contributing to incomplete understanding of human diseases.

We overcame these technological barriers by developing the forward genetic Mirror approach, which we applied to identify components of two human nuclear pathways targeting the 3′ end of MALAT1. One of them is the pathway of 3′ end maturation of MALAT1 and the other is the degradation pathway blocked by the triple-helical ENE stability element. Genome-wide Mirror screening identified numerous components of the three major nuclear RNA-processing complexes: Ribonuclease P (RNase P), the nuclear RNA Exosome, and the Nuclear Exosome Targeting (NEXT) complex, as well as three additional genes: DDX59, C1D, and BRF2. We found that the DEAD box helicase DDX59, which lacked a known substrate or role in nuclear RNA degradation, influences nuclear RNA degradation through an extensive retention of U12 introns within components of the RNA Exosome and the NEXT complex, suggesting a previously unknown function of DDX59 in minor intron splicing. Additional retained U12 introns affect genes associated with the rare genetic developmental disorder Oral-Facial-Digital (OFD) syndrome, suggesting a mechanism for DDX59 association with OFD. We further observe effects of DDX59 on endogenous snRNAs. Collectively, we demonstrate that the Mirror forward genetic approach efficiently identifies components of nuclear pathways acting on human nuclear lncRNAs, revealing surprising connections to human disease.

## RESULTS

### Design and implementation of the Mirror approach for forward genetic discovery of nuclear factors targeting 3′ end of the human MALAT1

Since translation-dependent reporter systems cannot be used directly for forward genetic interrogation of pathways acting on strictly *nuclear* human lncRNAs, we developed the Mirror approach (**Fig. 1d**), which relies on nuclear export of a fragment of non-polyadenylated nuclear RNA of interest to produce fluorescence signal. Here, we employed the previously reported finding that a GFP reporter carrying the downstream 3′ triple-helical ENE sequence of MALAT1 or MEN-β can undergo nuclear export and translation into a functional fluorescent protein^2^. We reasoned that inhibition of nuclear pathways acting on a similarly-designed reporter would affect (i.e., be ’mirrored’ by) its steady-state cytoplasmic fluorescence, enabling forward genetic screening for components of human nuclear machinery acting on lncRNAs that would otherwise never leave the nucleus. We further reasoned that whereas the genome-wide forward genetic screening using such a Mirror reporter would simultaneously identify components of both nuclear (black) and cytoplasmic (grey) pathways acting on it (schematically shown in **Fig. 1d**), subsequent post-screening analysis of the effects of knockout of individual identified genes on the endogenous nuclear lncRNA (**Fig. 1d**) will distinguish components of nuclear and cytoplasmic pathways, identifying components of nuclear pathways acting on the strictly nuclear lncRNA of interest.

We employed Mirror for the forward genetic identification of two human pathways acting on the 3′ end of the strictly nuclear lncRNA MALAT1: one of them is the pathway of the 3′ end maturation of MALAT1 and the other is the pathway of MALAT1 nuclear degradation that is blocked by the triple-helical ENE element (these two distinct pathways are identified by Mirror simultaneously in a single genetic screen). Each Mirror reporter employs multiple signal- and sensitivity-enhancing features of the previously described single-cell-based dual-color Fireworks system^35^. Similarly to Fireworks, each Mirror reporter expresses a single polyprotein consisting of (1) repeats of fluorescent proteins (GFP or RFP), (2) TEV protease, and (3) β-globin, which contains an intron to improve nuclear export. The proteolytic activity of TEV protease separates fused fluorescent proteins, the protease, and the β-globin to eliminate negative effects on localization, maturation, and the half-life of the fused fluorescent proteins. These features increase the fluorescence signal produced by a single transcription unit of the reporter to improve the sensitivity of the genetic screen^35^. Uniquely, each Mirror reporter (**Figs. 1e** and **S1a**) carries two elements of the human lncRNA MALAT1: the 3′ ENE stability element and the tRNA-like mascRNA element^18^ (**Fig. 1a-b**). The ENE of one of these reporters carries a single-nucleotide substitution, C8351G (**Figs. 1a, c-e** and **S1a**), introduced to make it more susceptible to nuclear degradation^18,19^. To enhance screening specificity, each individual Mirror cell expresses a second, wild-type ENE control reporter that differs from the stability-compromised ENE by just a single nucleotide (**Figs. 1e**). The ENE(C8351G)-mascRNA(WT)-containing Mirror reporters (**Fig. 2a**) yield lower steady-state RNA levels (**Fig. 2c**) and fluorescence (**Fig. 2e**) than those containing ENE(WT)-mascRNA(WT). To distinguish the effects of gene knockout on ENE-mascRNA from those on the fluorescent protein moiety of the reporters, and to enhance the rigor of screening and validation, we employed two orthogonal Mirror cell lines. These cell lines, named “Red” and “Green”, have WT ENE and ENE(C8351G) swapped (**Figs. 1e**, **S1a**, and **S2**).

**Figure 2.**
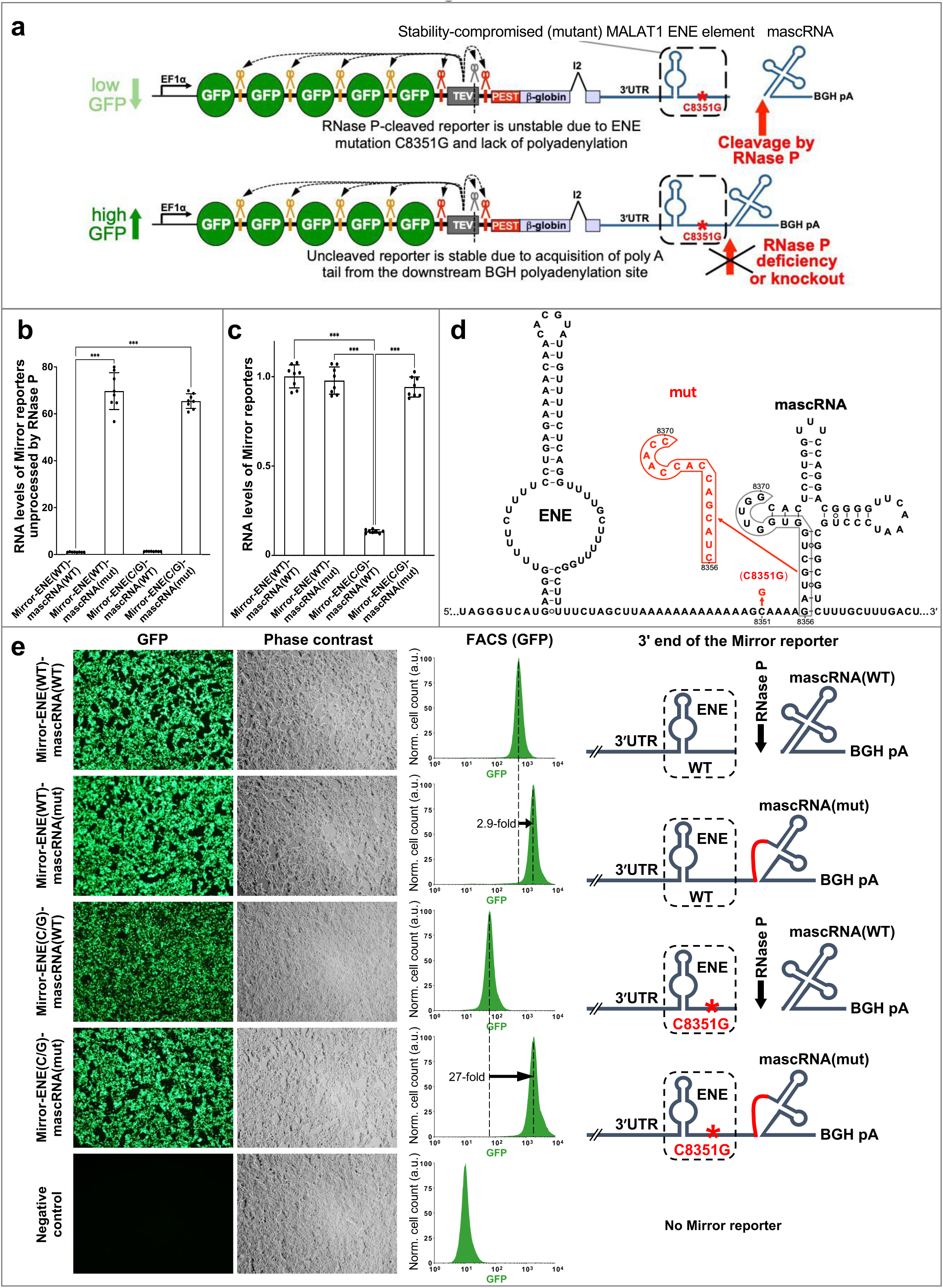
The Mirror reporters enable forward genetic identification of human factors required for MALAT1 processing at the ENE-mascRNA junction. **a.** Schematic of the effect of RNase P cleavage defect on fluorescence of the Mirror reporter. Low RNA levels of the RNase P-processed MALAT1(C8351) Mirror reporter result from a stability-compromising mutation (highlighted in red) within its 3′ ENE, leading to low GFP fluorescence. Inhibition of mascRNA cleavage causes a switch to the downstream Bovine Growth Hormone (BGH) cleavage and polyadenylation site, increasing stability and thus GFP fluorescence of the Mirror reporter, enabling identification of human factors required for MALAT1 3′-end processing. **b.** RT-qPCR quantification of RNase P processing of Mirror reporter variants shown in panels d and e. Values are expressed relative to the RNase P-unprocessed Mirror-ENE(WT)-mascRNA(WT) transcript levels. ΔΔCt was used to quantify the relative expression levels. Data are presented as means of two biological x four technical replicates; error bars represent standard deviation. Statistically significant differences between knockout and control samples were determined by one-way ANOVA. Posthoc comparisons using Tukey’s HSD test were conducted to determine the overall difference between groups, and labeled as “*”, P<0.05; “**”, P<0.01; “***”, P<0.001. Sequences of qPCR primers are listed in Supplemental Data S2. **c.** RT-qPCR quantification of steady-state RNA levels of Mirror reporter variants (shown in panels d and e) transiently transfected in HEK293T cells. RNA levels were normalized to blasticidin S resistance (bsr)^69^ mRNA expressed from a co-transfected control plasmid. All qPCR statistical tests for panels c were performed as described for panel b; qPCR primers are listed in Supplemental Data S2. **d.** Sequences and mutations in ENE-mascRNA regions of Mirror reporters shown in panels a, b, c, and e. Substitutions that destabilize ENE (C8351G)^19^ or disrupt RNase P cleavage (mut 8356-8370) at the ENE-mascRNA junction are highlighted in red. **e.** Fluorescence of transiently transfected Mirror reporters visualized using Olympus IX70 fluorescence microscope and FACS analysis of the fluorescence of FRT site-incorporated Mirror reporters carrying 3′-end variants schematically shown to the right (nucleotide substitutions are detailed in panel d).

To identify the nuclear degradation machinery that targets the 3′ end of MALAT1 but is prevented from degrading MALAT1 by its intact triple-helical ENE(WT), we hypothesized that knockout of a component of this degradation machinery (**Fig. 1c**) would result in a greater increase in the fluorescence of reporters with stability-compromised ENE(C8351G) compared to reporters with ENE(WT) (**Figs. 1e, f** and **S1a**, **b**).

To identify components of the second pathway required for 3′ end maturation of MALAT1, including separation of the tRNA-like mascRNA from the MALAT1 lncRNA precursor (**Fig. 1b**), Mirror leverages the enhanced stability – and, as a result, increased fluorescence – of the stability-compromised ENE(C8351G) Mirror reporter upon inhibition of mascRNA cleavage (**Fig. 2**), which stems from the resulting use of the downstream Bovine Growth Hormone (BGH) cleavage and polyadenylation site (**Fig. 2a**). Similar disruption of mascRNA cleavage and the resulting switch to BGH-directed polyadenylation produce considerably smaller changes in RNA levels (**Fig. 2b-d**) and fluorescence of the ENE(WT) reporter. These changes are 2.9-fold and 27-fold for the ENE(WT) and ENE(C8351G), respectively (**Fig. 2e**). As a result, defects in MALAT1 nuclear 3′-end processing are readily identifiable through their differential effects on fluorescence of the same-cell ENE(C8351G) and ENE(WT) Mirror reporters. We chose HeLa cells for Mirror forward genetic screening of MALAT1-acting human nuclear pathways due to the strong association of steady-state levels of MALAT1 with tumor aggressiveness in cervical cancers^14^.

In summary, the Mirror approach is designed to simultaneously identify two nuclear pathways acting on the MALAT1 3′ end. Components of the first—the degradation pathway targeting the 3′ end of MALAT1 but prevented from degrading MALAT1 by its intact triple-helical ENE—are expected to affect degradation of the full-length MALAT1 with the stability-compromised ENE(C8351G). Components of the second—the pathway required for MALAT1 3′-end processing—are expected to affect mascRNA cleavage and, potentially, the levels of the wild-type endogenous MALAT1.

### Genome-wide Mirror forward genetic screening for components of two pathways acting on the 3′ end of the human lncRNA MALAT1 enriched numerous components of *nuclear* RNA-processing complexes

Figures 3 and **S1c** show the outcomes of two iterative rounds of CRISPR sgRNA-based forward genetic Mirror screening to simultaneously identify components of two human nuclear pathways targeting MALAT1’s 3′ end: one being the pathway of its 3′-end maturation and the other being the degradation pathway blocked by the wild-type ENE element. The screening was conducted in the "Red" and “Green” Mirror cell lines (**Fig. 1e** and **S1a**) using FACS-based iterative rounds of guide RNA (sgRNA) library enrichment (**Fig. 3b**) intended to amplify pathway-inhibiting sgRNAs and eliminate stochastic false positives^35^. During the 1^st^ round of screening (**Fig. 3a** and **S1c**), the Mirror cell lines were transduced with the original lentiviral GeCKO-lentiCRISPR sgRNA library^36^ containing 64,751 sgRNAs targeting 18,080 human genes. During the 2^nd^ round of screening (**Fig. 3a** and **S1c**), the original “Red” and “Green” Mirror cell lines (**Fig. 1e** and **S1a**) were transduced with the enriched lentiviral sgRNA libraries, which were obtained at the end of the 1^st^ round of screening as shown in **Fig. 3b**. We expected that sgRNAs targeting each of the two pathways would increase (1) GFP fluorescence of the “Red” MALAT1 Mirror cells (*i.e.*, shift them up, **Figs. 1f** and **3a**) and (2) RFP fluorescence of the “Green” MALAT1 Mirror cells (*i.e.*, shift them right, **Fig. S1b**, **c**). Indeed, distinct markedly enriched populations of cells appeared in the blue gates, comprising 0.08% and 7.64% of cells (**Fig. 3a**) and the green gate, comprising 0.024% and 4.53% of cells (**Fig. S1c**), after the 1^st^ and 2^nd^ rounds of screening, respectively.

**Figure 3.**
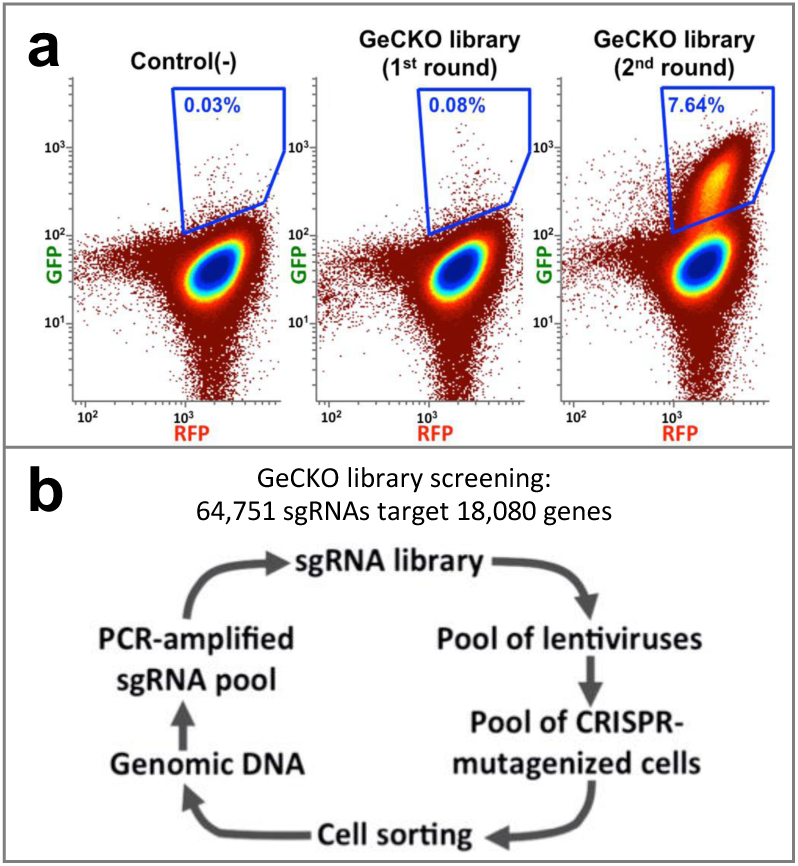
Iterative forward genetic Mirror screening to identify factors required for MALAT1 3′-end processing, as well as factors blocked from degrading MALAT1 by its triple-helical 3′-end ENE element. **a.** Genome-wide iterative FACS-based forward genetic screening in the “Red” (Fig. 1e) Mirror reporter cell line (analogous screening in the orthogonal “Green” Mirror cell line is shown in **Fig. S1c**). **b.** Schematics of iterative forward genetics screening rounds employed in panel a and **Fig. S1c**.

Deep sequencing of sgRNAs in the screening-enriched cell population (0.08% of cells in the blue FACS gate in **Fig. 3a**) revealed numerous sgRNAs targeting *nuclear* RNA-processing factors (**Fig. 4a** and **e1**). Indeed, 51 of the top 93 most abundant sgRNAs in the screening-enriched library (**Fig. 4a**) targeted: (i) 9 out of 10 protein components of the human nuclear RNase P^37^: RPP14, RPP21, RPP30, RPP38, RPP40, POP1, POP4 (RPP29), POP5, and POP7 (RPP20); (ii) 9 out of 11 components of the human RNA Exosome^38^: EXOSC2, EXOSC4, EXOSC5, EXOSC6, EXOSC7, EXOSC8, EXOSC9, EXOSC10, including its nuclear DIS3 (but not the cytoplasmic DIS3L) component^39^; (iii) 2 out of 3 components of the human nuclear NEXT (Nuclear Exosome Targeting) complex^40^: RBM7 and SKIV2L2 (hMTR4); and (iv) the nuclear exosome co-localized RNA-binding protein C1D known for its role in 3′- end processing of 5.8S rRNA^41^. For some of these factors, Mirror identified more than one sgRNA (**Fig. 4a**), and for others, such as RNase P components RPP14, RPP38, POP7, and POP5, nuclear RNA exosome components EXOSC2, EXOSC7, EXOSC8, EXOSC9, EXOSC10, as well as the exosome-associated C1D and the NEXT complex component SKIV2L2, it identified, among the top 93 sgRNAs, every sgRNA present in the original GeCKO library (**Fig. 4a** and **Supplemental Data S1**). Additional Mirror-identified and subsequently validated sgRNAs targeted TFIIB-like factor BRF2, known to control initiation of Pol lII^42^ and the predominantly nuclear DEAD-box helicase DDX59, which has no known substrate in any organism^43,44^. All Mirror-identified factors are summarized in **Figure 4e**1.

**Figure 4.**
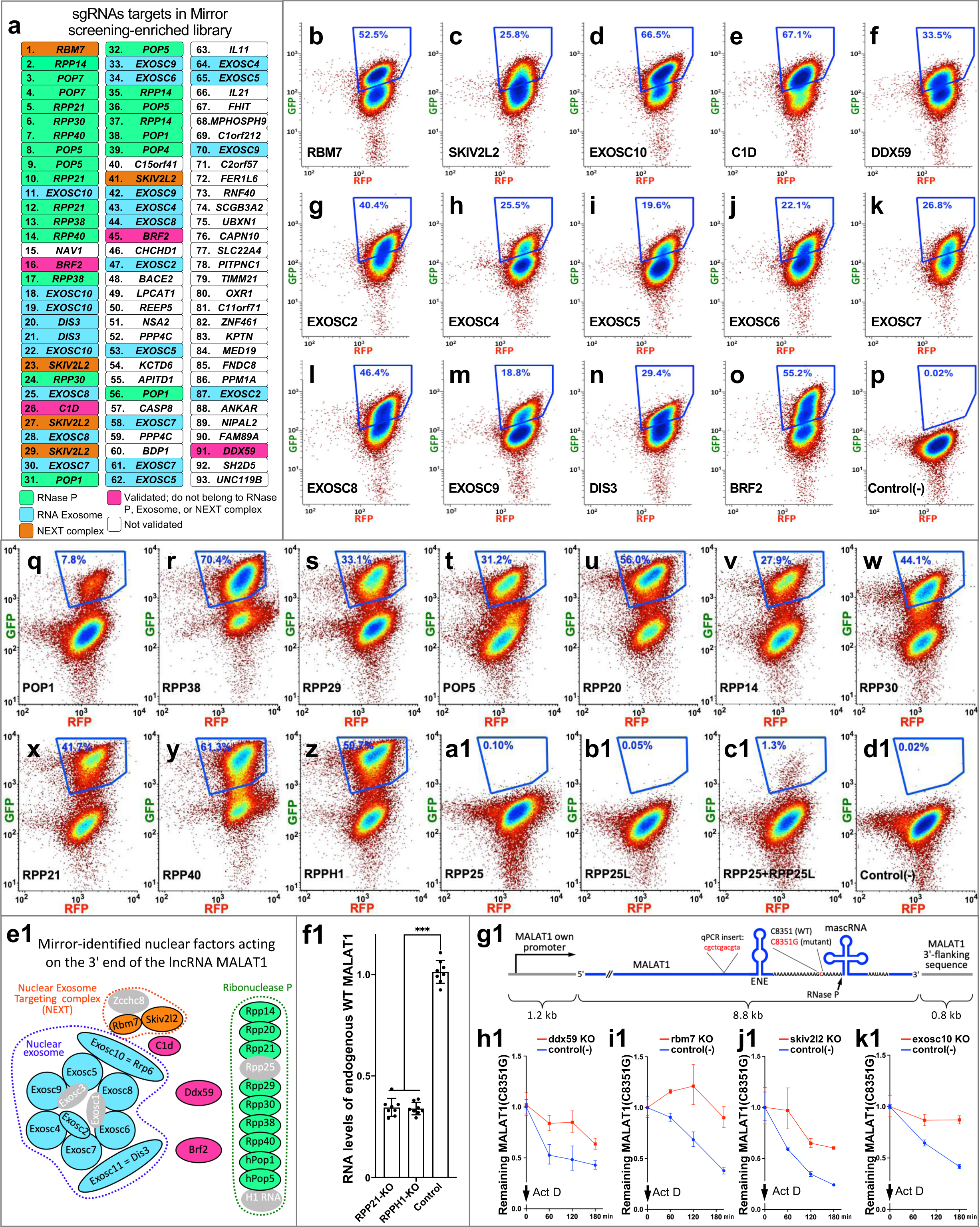
Mirror-identified components of human nuclear pathways acting on MALAT1’s 3′-end and their validation. **a.** sgRNAs and their gene targets ranked by their abundance in the Mirror screening-enriched library. Components of RNase P, RNA Exosome, and the NEXT complex are highlighted in green, blue, and orange, respectively. Three additional Mirror-identified and validated below genes – DDX59, C1D, and BRF2 – are shown in purple. **b-y** and **d1.** Validation of Mirror-identified sgRNAs in the “Red” MALAT1 Mirror cell line, which was transduced with individual sgRNAs and analyzed by FACS as described in Materials and Methods. **z-d1.** Validation of additional components of Mirror-identified nuclear complexes. **e1.** Schematics of Mirror-identified complexes acting on the 3’ end of MALAT1. Mirror-identified complex components are highlighted consistent with their coloring in panel a. **f1.** RT-qPCR analysis of the effects of knockouts of the RNase P-specific components (i.e., not shared with RNase MRP), RPP21 and RPPH1, on the levels of the endogenous wild-type lncRNA MALAT1. For RT-qPCRs, RNA levels were normalized to 18S RNA; ΔΔCt was used to quantify the relative expression levels. Data are presented as means of two biological x four technical replicates; error bars represent standard deviation. Statistically significant differences between knockout and control samples were determined by one-way ANOVA. Posthoc comparisons using Tukey’s HSD test were conducted to determine the overall difference between groups, and labeled as “*”, P<0.05; “**”, P<0.01; “***”, P<0.001. Sequences of qPCR primers are listed in Supplemental Data S2. **g1.** Schematics of the full-length lncRNA MALAT1(C8351G) driven by its own promoter, followed by its own 3′-flanking sequence, and carrying an insertion of 11 nucleotides in a non-conserved region to distinguish it from endogenously-expressed MALAT1 using RT-qPCR. **h1-k1**. Knockouts of Mirror-identified DDX59, RBM7, SKIV2L2, and EXOSC10 inhibit degradation of the full-length lncRNA MALAT1 with a stability-compromised ENE(C8351G). RT-qPCR-derived MALAT1(C8351G) levels were normalized to 18S RNA and are shown relative to the time of Actinomycin D addition. ΔΔCt was used to quantify the relative expression levels. Data are presented as means of two biological x four technical replicates; error bars represent standard deviation.

Underscoring Mirror’s efficiency in identifying *nuclear* factors acting on *nuclear* RNAs using reporter’s *cytoplasmic* fluorescence, transduction of sgRNAs targeting the individual Mirror-enriched factors (**Fig. 4a** and **4e1**) produced specific cell populations with a 2- to 15-fold increase in GFP fluorescence (**Fig. 4b-y** and **d1**). For all 23 genes shown, only minimal effects are observed on the fluorescence of the same-cell control RFP reporter (**Figs. 1e** as well as **4b-y** and **d1**). Notably, 21 out of 23 (91%) of these genes are classified as "Common Essential" in the DepMap database^45^, demonstrating the Mirror’s effectiveness in identifying genes critical for viability.

### Corroborating the design, one set of the Mirror-identified factors impacts the 3′-end processing and the levels of the endogenous wild-type MALAT1

Collectively, Mirror correctly identified 9 out of 10 known protein components of RNase P required for MALAT1 3′-end processing (**Fig. 4a, 4q-e1**). We hypothesized that the absence of RPP25 (hPOP6) among the RNase P protein components in Mirror results reflects a limitation of forward genetics in detecting redundant factors. Indeed, whereas individual knockouts of RPP25 and RPP25L^46^ produced no populations with increased green fluorescence (**Fig. 4a1** and **b1**), a simultaneous double knockout of these genes produced such a population (**Fig. 4c1**), confirming functional redundancy.

Since the GeCKO library lacked sgRNAs targeting the catalytic RNA component of RNase P, RPPH1, identifying it using this library would not have been possible. We demonstrate, however, that not only do sgRNAs targeting protein-encoding genes increase fluorescence in the Mirror system, but also those targeting RNA-encoding genes, as shown by the CRISPR-sgRNA knockout of RPPH1 (**Fig. 4z**).

We show that knockouts of the RNase P-specific (i.e., not shared with RNase MRP) components impact the levels of the endogenous wild-type lncRNA MALAT1 (**Fig. 4f1**), underscoring the unique ability of the Mirror approach to genetically identify nuclear factors that impact the 3′-end processing and levels of this cancer-associated non-coding RNA.

### Corroborating the design, the second set of Mirror-identified complexes and factors impacts the nuclear degradation of the full-length lncRNA MALAT1 with stability-compromised ENE

As shown in **Fig. 4e1**, the second set of Mirror-identified factors is part of the established nuclear RNA-degradation complexes, such as the RNA Exosome and the NEXT complex. However, the predominantly nuclear DEAD-box helicase DDX59 could not be readily assigned to them. Furthermore, nothing is known about the substrate of DDX59 in any organism, nor has it been determined whether this substrate is RNA or DNA^14^. To ascertain how DDX59 affects 3′ end of MALAT1, we first confirmed that DDX59 knockout specifically affects the fluorescence of Mirror reporters with a stability-compromised 3′ ENE(C8351G) and that additional sgRNAs toward DDX59 can replicate this effect. In addition to reproducing an increase in fluorescence (**Fig. 4f**) in the "Red" Mirror cell line (**Fig. 1e, f**), in which the first step of validation was performed (**Fig. 4b-d1**), the individual transductions of four distinct DDX59-targeting sgRNAs into the orthogonal "Green" Mirror cell line (**Fig. S1a**), in which the ENE(C8351G) and ENE(WT) are swapped, produced an increase in RFP fluorescence without significantly affecting GFP fluorescence (**Fig. S1b** and **d-h**). This result confirms that DDX59 knockout specifically influences the fluorescence of Mirror reporters with a stability-compromised ENE, and not the wild-type ENE, and ruled out off-target effects of sgRNAs on the fluorescent protein moiety of the reporters.

To confirm that the effects of DDX59 knockout on fluorescence of stability-compromised Mirror reporters represent an accurate reflection of its effects on nuclear degradation of the RNA of MALAT1 with stability-compromised ENE, we analyzed the degradation of the full-length MALAT1(C8351G) RNA expressed from its own native promoter (**Fig. 4g1**) following the CRISPR sgRNA-based knockout of DDX59 and the cessation of transcription with actinomycin D. Indeed, knockout of DDX59 (quantified in **Fig. 6g**) results in stabilization of the full-length MALAT1(C8351G) RNA (“red” curve in **Fig. 4h1**) as compared with non-targeting sgRNA control (“blue” curve in **Fig. 4h1**). As expected for the same-cell stable control provided by the WT ENE during screening, and in contrast to MALAT1 with the stability-compromised ENE(C8351G), we do not expect, nor we do observe, a major effect of DDX59 knockout on endogenous WT MALAT1 during the same actinomycin D time course or on its steady-state levels (**Fig. S1i, j**).

**Figure 5.**
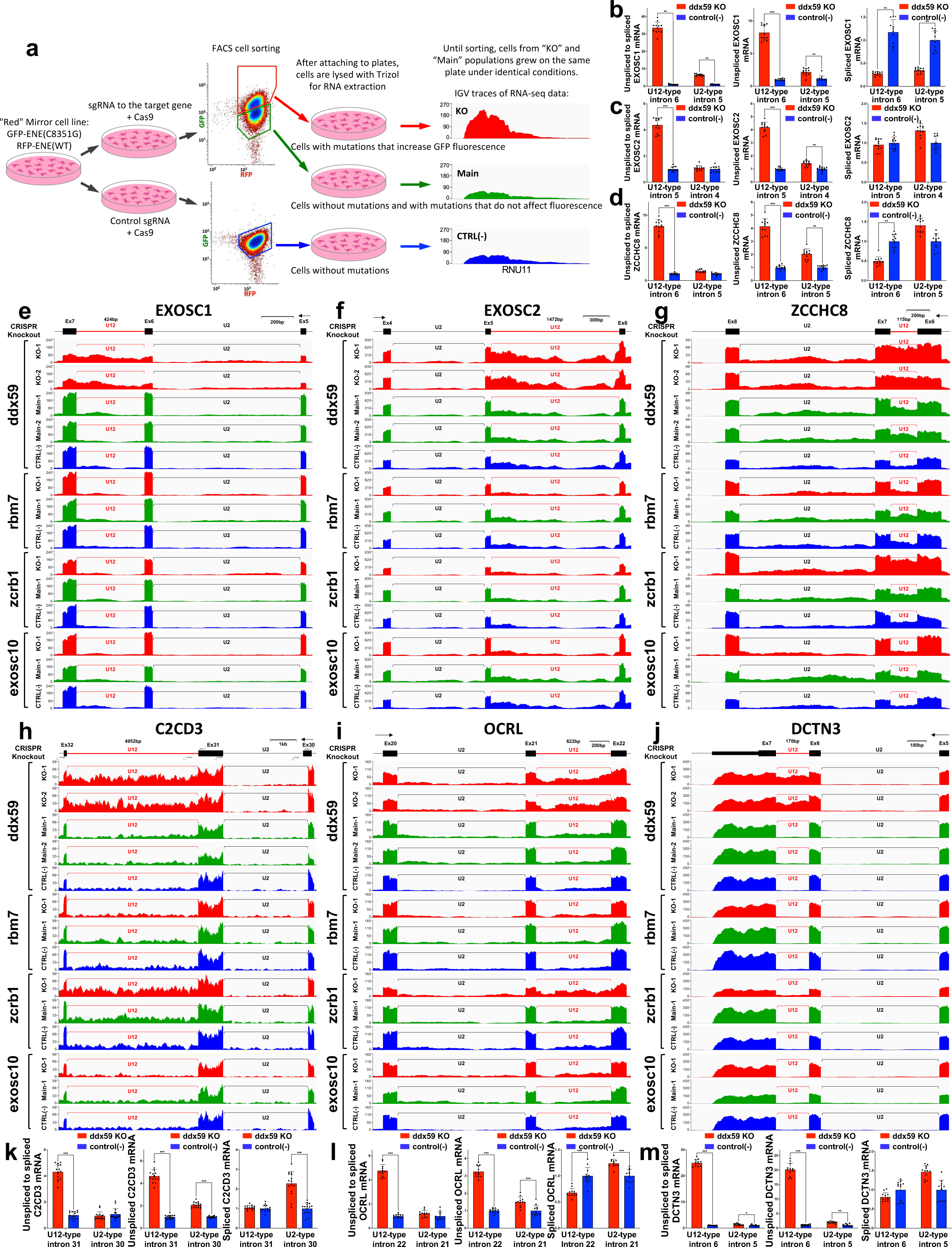
DDX59 is required for the proper splicing of U12 introns; its deficiency disrupts the RNA exosome, the NEXT complex, and genes crucial for Oral-Facial-Digital syndrome (OFD) and ciliary function. **a.** Schematics of generating knockout (KO), Main, and CTRL(-) populations for RNA-seq analysis, in which respective traces are colored in red, green, and blue. These knockouts are rigorously controlled for growth conditions, transduction efficiency, and antibiotic selection, as, throughout the entire experiment, the cell populations grow together on the same plate until sorted by FACS. The IGV tracks are shown only as an illustration of color-coding that is further used in **Figs. 5b-m, 6a-e, S3, S4**, and **S6. b-d.** RT-qPCR quantification of intron retention in mRNAs of the exosome components EXOSC1 and EXOSC2 as well as the scaffold component of the Nuclear Exosome Targeting (NEXT) complex, ZCCHC8 after DDX59 knockout as described in panel a. **e-j.** Integrated Genome Viewer (IGV) traces of RNA-seq reads for minor intron-containing genes, the exosome components EXOSC1 and EXOSC2, as well as the scaffold component of the Nuclear Exosome Targeting (NEXT) complex, ZCCHC8, and genes associated with ciliary function, DCTN3 and OCRL, as well as C2CD3, which is also associated with Oral-Facial-Digital (OFD) syndrome. Cell populations with DDX59, RBM7, ZCRB1, and EXOSC10 knockouts were obtained as shown in panel a and their respective IGV traces are colored accordingly. Major and minor introns are labeled as U2 and U12, respectively. **k-m.** RT-qPCR quantification of intron retention in mRNAs of genes associated with ciliary function, DCTN3 and OCRL, as well as C2CD3, which is also associated with Oral-Facial-Digital (OFD) syndrome. For RT-qPCRs, RNA levels were normalized to 18S RNA; ΔΔCt was used to quantify the relative expression levels. Data are presented as means of two biological x four technical replicates; error bars represent standard deviation. Statistically significant differences between knockout and control samples were determined by one-way ANOVA. Posthoc comparisons using Tukey’s HSD test were conducted to determine the overall difference between groups, and labeled as “*”, P<0.05; “**”, P<0.01; “***”, P<0.001. Positions of qPCR primers are indicated and their sequences are listed in Supplemental Data S2.

**Figure 6.**
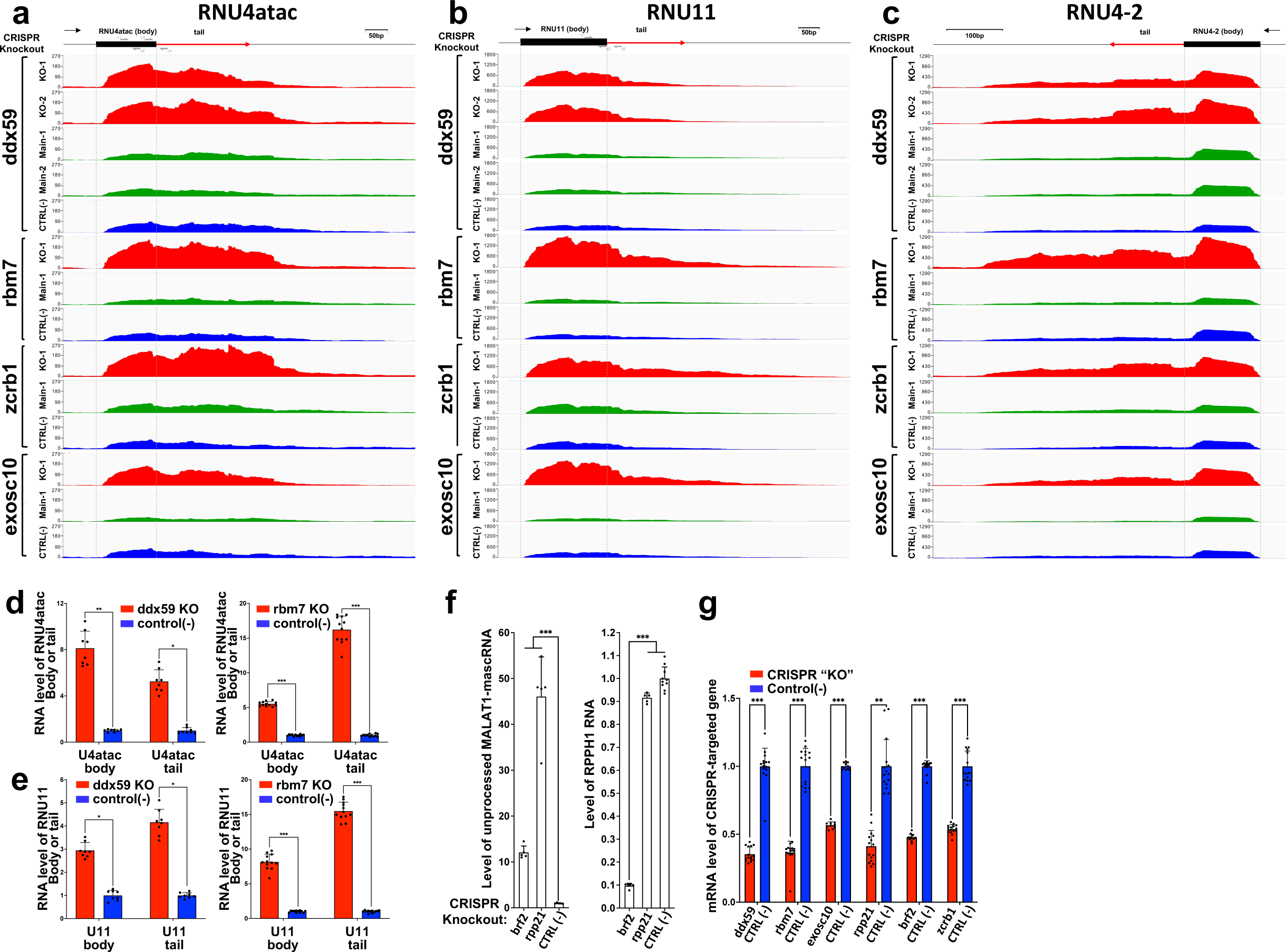
Knockout of DDX59 results in the accumulation of 3′-extended precursors of endogenous snRNAs. **a-c.** Integrated Genome Viewer (IGV) traces of RNA-seq (performed without poly(A) selection) are shown for minor spliceosomal snRNAs, U4atac and U11 (panels a and b), and for a major spliceosomal snRNA, U4 (panel c). Cell populations with knockouts of DDX59, RBM7, ZCRB1, and EXOSC10 were obtained as shown in Fig. 5a; their respective IGV traces are colored accordingly.**d-e.** RT-qPCR quantification of the levels of 3′-extended forms of minor spliceosomal snRNAs U4atac (panel d) and U11 (panel e). **f.** RT-qPCR analysis of the effects of BRF2 knockout on (i) processing of the endogenous full-length lncRNA MALAT1 by RNase P and (ii) RNA levels of its catalytic component RNA H1. The left graph shows analysis of cleavage across the RNase P site of the endogenous full-length lncRNA MALAT1, and the right graph shows RNA levels of RNA H1 in knockout cell populations. Knockout of RPP21, which is unique for RNase P (and absent in the closely-related RNase MRP) and essential for its function^37^, is shown as a positive and negative control in the left and right graphs, respectively. **g.** RT-qPCR quantifications of mRNA levels in cell populations with CRISPR-induced knockouts of indicated genes, obtained as shown in Fig. 5a. All qPCR quantifications and statistical tests were performed as described in the legend of Fig. 5; qPCR primers are listed in Supplemental Data S2.

The stabilizing effect of the DDX59 knockout on full-length MALAT1(C8351G) RNA (**Fig. 4h1**) parallels the effects observed with knockouts of EXOSC10, SKIV2L2, and RBM7 (quantified in **Fig. 6g**) for this lncRNA (**Fig. 4i1-k1**). Since EXOSC10, a catalytic subunit of the nuclear RNA Exosome, as well as RBM7 and SKIV2L2, subunits of the Nuclear Exosome Targeting (NEXT) complex^40^, are key constituents of two major established complexes involved in nuclear RNA degradation^47,48^, their stabilizing effects corroborate correct identification of these complexes by Mirror. The parallel effects of DDX59 suggest it is also required for the nuclear degradation of stability-compromised MALAT1 through a yet-unidentified mechanism.

### DDX59 knockout results in retention of minor (U12) introns, impacting components of the RNA Exosome and NEXT complex

To address the role of DDX59 in the nuclear degradation of stability-compromised lncRNA MALAT1, we analyzed transcriptome-wide effects of DDX59 knockout.

Remarkably, knockout of DDX59, rigorously obtained as shown in **Figure 5a**, results in profound retention of minor (U12) spliceosomal introns across numerous genes (**Figs. 5**, **S3d-i**, **S4**, and **S6**). To illustrate the rigor of the FACS-isolation strategy, we compare the knockout cell population (“KO”, red traces) not only to cells transduced with a control guide RNA (“CTRL(-)”, blue traces), but also to cells propagated on the same plate as knockout cells but lacking a gene knockout (“Main”, green traces). The second control mitigates unintended differences in growth conditions, transduction efficiency, and antibiotic selection, as throughout the entire experiment, the cells of the "KO" and "Main" populations grow together on the same plate until sorted by FACS (**Fig. 5a**). This retention is more pronounced than that resulting from the knockout of the minor spliceosomal factor ZCRB1^49,50^, yet exhibits little to no effect on major (U2) introns (**Figs. 5**, **S3d-i**, **S4**, and **S6**; also shown for U2 introns of housekeeping genes in **Fig. S3a-c**). Importantly, neither the knockout of the Mirror-identified NEXT complex component RBM7, nor the knockout of the exosome component EXOSC10 (quantified in **Fig. 6g**) shows comparable retention of U12 or U2 introns (**Figs. 5**, **S3**, **S4**, and **S6**), suggesting that the increase in retained minor introns observed with DDX59 knockout is not due to the stabilization of unspliced transcripts resulting from a deficiency in these complexes. The U12 intron retention is evident not only for DDX59 knockout cells compared to negative control cells transduced with a non-targeting sgRNA (compare the red "KO" IGV traces with the blue "CTRL-" traces in **Figs. 5**, **S3d-i**, **S4**, and **S6**) but also for DDX59 knockout cells compared to cells in which the same sgRNA toward DDX59 yielded no FACS-detectable increase in fluorescence (compare the red "KO" IGV traces with the green "Main" traces in **Figs. 5**, **S3d-i**, **S4**, and **S6**). Quantification of U12 intron retention (**Figs. 5b-d** and **k-m**, **S4b**, and **S6b**) demonstrates that DDX59 is required for the splicing of a set of human minor introns.

As shown in **Figures 5b-g** and **S4g**, the genes with pronounced DDX59-dependent retention of minor introns include RNA Exosome components EXOSC1, EXOSC2, and EXOSC5, as well as the scaffold component of the Nuclear Exosome Targeting (NEXT) complex, ZCCHC8. Since Mirror screening and its validation have shown that both the RNA Exosome and the NEXT complex are required for the degradation of stability-compromised lncRNA MALAT1(C8351G), inactivation of integral components of these complexes through U12 intron retention (**Figs. 5b-g** and **S4g**) and subsequent reduction of their protein levels (**Fig. S5**) explains the Mirror-identified role of DDX59 in the stabilization of lncRNA MALAT1 with a compromised 3′-end triple-helical ENE structure.

### Retention of numerous minor (U12) introns in cilia-related genes provides an explanation for the role of DDX59 in Oral-Facial-Digital syndrome (OFD)

Whereas mutations in the DEAD-box helicase DDX59 are known to associate with the rare genetic developmental disorder Oral-Facial-Digital (OFD) syndrome^43,51^, the mechanism linking defects in DDX59 to OFD remains unknown. It has been found that nearly 20 genes encoding proteins that either represent components of cilia or influence ciliogenesis are associated with at least 14 classes of OFD^52–54^. However, DDX59 is a notable exception among OFD-associated genes because it lacks known cilia-associated roles^52^.

We found that knockout of DDX59 induces retention of U12 introns in mRNAs of genes with established roles in OFD and OFD-associated ciliopathies such as C2CD3, TCTN3, and TMEM107 (**Figs. 5h** and **k**, **S4b** and **d**), mutations in which are associated with, and have been proposed to be causative for, OFD subtypes OFDXIV, OFDIV, and OFDVI, respectively^52–54^. Additionally, knockout of DDX59 results in retention of U12 introns in DCTN3, OCRL, C2CD3, ARMC9, KATNIP, SFI1, CUL1, ACTR10, CCDC28B, PPP5C, TCTN3, ACTL6A, TMEM107, KIFAP3, RABL2A (**Figs. 5h-m**, **S3d-i**, and **S4a-f**), all of which are associated with ciliogenesis or cilia assembly^55^.

Whereas the knockout of DDX59 produces a major effect on retention of minor (U12) spliceosomal introns, retention of major (U2) spliceosomal introns is virtually unaffected (**Figs. 5**, **S3**, **S4**, and **S6**). As negative controls, knockouts of the NEXT complex component RBM7 and the exosome component EXOSC10 (quantified in **Fig. 6g**) do not increase retention of either U12 or U2 introns (**Figs. 5**, **S3**, **S4**, and **S6**). As a positive control, knockout of ZCRB1 increases retention of U12 introns (**Figs. 5**, **S3**, **S4**, and **S6**), as expected for a knockout of a known component of the human minor spliceosome^49,50^. Collectively, the DDX59-induced retention of U12 introns in OFD-associated and cilia-related genes provides an explanation for its role in Oral-Facial-Digital syndrome.

### Knockout of DDX59 results in the accumulation of 3′-extended precursors of endogenous small nuclear RNAs (snRNAs)

Consistent with the Mirror-identified stabilizing effect of DDX59 knockout on the nuclear MALAT1(C8351G), which undergoes non-canonical 3′-end processing by RNase P, lacks a poly(A) tail, and is partially devoid of the protection of the intact 3′-end triple-helical ENE structure, we observe that CRISPR-Cas9 knockout of DDX59 increases the steady-state levels of 3′-extended precursors of several endogenous Sm-class snRNAs (**Fig. 6a-c**) known to undergo complex non-canonical 3′-end processing and lack poly(A) tails^56,57^. RT-qPCR quantification and IGV traces of RNA-seq performed without poly(A) selection show a 5.3- and 4.1-fold increase in the levels of 3’-extended forms of snRNAs U4atac and U11, respectively (**Fig. 6d, e**). Consistent with (i) our finding that knockout of DDX59 impacts the function of the RNA Exosome and the NEXT complex via retention of minor (U12) introns in mRNAs of key components of these two complexes (**Figs. 5e-g** and **S4g**), and (ii) known roles of both complexes in the 3′ end processing of snRNA precursor transcripts^48,58^, knockouts of a known minor spliceosome component ZCRB1, RNA Exosome component EXOSC10, and the NEXT complex component RBM7 similarly increase the levels of 3′-extended forms of U4atac, U11 and U4-2 (**Fig. 6a-c**). Two lines of evidence suggest that the increase in 3′-extended forms of minor spliceosomal snRNAs is not the primary cause of the minor intron retention observed in DDX59 knockout: First, whereas the effects of DDX59, RBM7, EXOSC10, and ZCRB1 knockouts on the levels of 3′-extended forms of U4atac, U11, and U4-2 snRNAs show significant similarities (**Fig. 6a-c**), only the knockouts of DDX59 and ZCRB1, not those of EXOSC10 or RBM7, result in U12 intron retention (**Figs. 5** and **S3**, **S4**, and **S6**). Second, whereas DDX59 knockout increases the levels of 3′-extended forms of both minor (U12) and major (U2) spliceosomal snRNAs (**Fig. 6a-c**), retention of only U12 introns is observed (**Figs. 5** and **S3**, **S4**, and **S6**). This suggests that inactivation of DDX59 impacts a subset of endogenous human nuclear non-canonical RNAs lacking poly(A) tails in a manner similar to that resulting from the minor spliceosome inactivation and the ensuring impairment of the RNA Exosome and NEXT complex.

### The Mirror-identified BRF2 is required for MALAT1-mascRNA processing through expression of the RNA component of RNase P, H1 (RPPH1)

To elucidate the effect of BRF2 (**Fig. 4o** and **e1**) on the 3′ end of MALAT1, we analyzed cleavage of the endogenous wild-type lncRNA MALAT1 (**Fig. 1b**) using RT-qPCR across the RNase P cleavage site. The quantification revealed a 12-fold increase in unprocessed MALAT1-mascRNA transcript following BRF2 knockout (**Fig. 6f** and **g**). This defect parallels a nearly 50-fold increase in RNase P-unprocessed MALAT1 following knockout of RPP21, which is an essential RNase P subunit (**Fig. 6f** and **g**). Additionally, knockout of BRF2 results in a dramatic 10-fold decrease in the steady-state levels of H1 (RPPH1) RNA (**Fig. 6f**), the catalytic RNA subunit of human RNase P^59^, whereas knockout of RPP21 (**Fig. 6g**) causes less than a 10% reduction (**Fig. 6f**). Together, these results demonstrate that BRF2, which controls initiation of human Pol III RNA transcripts^42^, is required for expression of Pol III-transcribed H1 RNA, and the loss of the H1 RNA leads to the MALAT1 3′-end processing defect identified by Mirror.

## DISCUSSION

We developed the Mirror approach, which enabled forward genetic identification of the human post-transcriptional machinery acting on nuclear non-coding RNAs by employing specialized non-polyadenylated reporters that undergo nuclear export and translation (**Figs. 1d** and **e**, **S1a**, and **S2**). We demonstrate that the fluorescence signal of the Mirror reporters provides a reliable proxy measurement for processing efficiency and/or abundance of a fused fragment of nuclear non-coding RNA of interest (**Figs. 2** and **4**), thereby enabling forward genetic screening of strictly nuclear pathways acting on it. Employed to discover two human nuclear pathways acting on the 3′ end of nuclear lncRNA MALAT1 with stability-compromised ENE, Mirror identified nearly all (**Fig. 4**) components of three major nuclear processing and degradation complexes: RNase P, RNA Exosome, and the NEXT complex. It additionally identified a nuclear exosome-associated factor C1D, TFIIB-like factor BRF2, and the DEAD box helicase DDX59 which previously had no known substrate in any organism or role in nuclear RNA processing and degradation (**Fig. 4e1**). Since Mirror does not rely on growth metrics and instead detects early changes in cell fluorescence that precedes the onset of the knockout-induced lethality, it efficiently identifies essential genes. Underscoring this feature, 21 of the 23 top genes (i.e., 91%) identified by Mirror (**Fig. 4e1**) are essential^45^. This comprehensive list, alongside with validation of effects on the endogenous lncRNA by at least one component of each of the three identified complexes (**Fig. 4i1-k1**) and additional factors (**Figs. 4h1** and **6f**), demonstrates the effectiveness of Mirror in identifying pathways acting on nuclear non-coding RNAs.

Of the three Mirror-identified genes outside the three major complexes, C1D plays an established role in nuclear RNA degradation as a SKIV2L2- and EXOSC10-interacting partner^41^, while BRF2 is involved in the initiation of Pol III transcription^42^, a process we confirmed as required for the expression of the RNase P catalytic subunit, RNA H1. Whereas Mirror accurately identified the DEAD box helicase DDX59 as judged by its impact on the degradation of the full-length lncRNA MALAT1(C8351G), the function of DDX59 remained obscure. Examination of the effects of DDX59 on nuclear RNA degradation revealed that they arise from a previously unknown role of DDX59 in minor intron splicing, as knockout of DDX59 produces extensive retention of U12 introns in components of the RNA Exosome EXOSC1, EXOSC2, and EXOSC5 as well as the NEXT complex component ZCCHC8, with the degree of retention being comparable to or greater than that observed with the knockout of the known minor spliceosome component ZCRB1^49,50^. Whereas retention of U12 introns in key components of the RNA Exosome and NEXT complex could potentially fully account for the observed (i) Mirror-identified stabilization of the full-length lncRNA MALAT1(C8351G) and (ii) increased levels of 3′-extended forms of endogenous Sm-class snRNAs, resulting from DDX59 knockout, an additional to minor intron splicing role of DDX59 in nuclear RNA degradation cannot be ruled out. Although no obvious physical associations of DDX59 with spliceosomes have been reported, such an interaction could be transient, which has been observed for DEAD-box helicases. Alternatively, DDX59 may be required for minor intron splicing through an indirect mechanism that remains to be elucidated; emerging studies continue to reveal unexpected twists in minor intron recognition^49,60^. Mirror’s identification of the roles of DDX59 in nuclear RNA degradation and minor intron splicing underscores the advantages of forward genetics in identifying components that impact pathways of interest, independently on whether such components act directly, indirectly, transiently, or are essential.

A new role of DDX59 in minor intron splicing provides a missing link between defects in DDX59 and the rare developmental genetic disorder Oral-Facial-Digital syndrome, which is often associated with intellectual disability^52^. Our finding that DDX59 knockout induces retention of minor introns in TCTN3, TMEM107, and C2CD3, all associated with distinct subtypes of OFD and linked to cilia and ciliogenesis, has revealed a mechanism by which mutations in DDX59 may affect these and potentially other OFD and cilia genes, contributing to this genetic disorder^43^. This mechanism of DDX59 in OFD is supported by several facts: First, deficiencies in cilia genes can cause OFD. Of the nearly 20 genes linked to at least 14 subtypes of OFD, only two, including DDX59, have not been definitively associated with cilia structure or function^52–54^. Second, a disproportionately high number (6%-7.6%) of cilia-related genes harbor U12 introns, compared to an average of 2% of U12-containing genes in the human genome^55^, suggesting a greater impact of minor splicing defects. Third, deficiencies in U12 intron splicing, including those caused by mutations in minor spliceosome snRNAs, are known to result in ciliary defects in humans^55^. Fourth, minor spliceosome deficiencies are known to associate with OFD syndrome via mis-splicing of genes affecting primary cilia^61^. Additionally, consistent with the impaired Sonic Hedgehog (SHH) signaling observed in DDX59-associated OFD^43^, DDX59 knockout leads to extensive retention of U12 introns in the mRNA of Derlin 2 (**Fig. S6**), a factor that mediates the retro-translocation of the SHH protein at the endoplasmic reticulum^43,61,62^. Altogether, our finding of DDX59’s impact on U12 intron splicing suggests that a deficiency in DDX59 may affect as many as 86 known cilia- and SHH-related genes spliced by the minor spliceosome^55,61^, contributing to OFD.

As we demonstrate with MALAT1, the Mirror strategy can be applied to more than one nuclear pathway, and may be extended to pathways of degradation, biogenesis, and surveillance of other nuclear non-coding RNAs. It also can be applied to identify human factors required to maintain steady-state levels of disease-associated non-coding nuclear RNAs, including the metastasis-promoting lncRNA MALAT1 itself, for which a search for destabilizing small molecules is underway^25,63^. Inhibition or inactivation of such factors would result in a decrease in levels of MALAT1, providing additional potential targets for cancers with poor survival linked to high MALAT1 levels^12–17^. Mirror’s unambiguous identification of the components of the nuclear ribonucleoprotein RNase P (**Fig. 4**), whose inactivation reduces the steady-state levels of the endogenous wild-type lncRNA MALAT1 (**Fig. 4f1**), represents the first example of the use of forward genetics to address this challenge.

In summary, the Mirror approach enables forward genetic discovery of components of human nuclear pathways acting on strictly nuclear non-coding RNAs. It efficiently identifies pathways components essential for cell viability, those that act indirectly, and, as a result, uncovers unexpected connections to human disease.

## EXPERIMENTAL PROCEDURES

### Construction of plasmids

*Mirror reporter plasmids*: The 171-nucleotide sequences (shown in Supplemental Data S2) of the human wild-type and C8351G mutant MALAT1’s ENE-mascRNA were amplified using PCR primers MALAT-Fg and MALAT-Rg, listed in Supplemental Data S2, from constructs C-G(WT) βΔ1,2-MALAT1 ENE+A+mascRNA and G-G(C8351G) βΔ1,2-MALAT1 ENE+A+mascRNA^19^, kindly provided by Dr. Jessica Brown. Mirror reporter plasmids were constructed by cloning these 171-mers into the unique PspXI restriction site of the Fireworks reporter plasmids pAVA2598[GFP(PTC-)] and pAVA2515[RFP(PTC-)]^35^ using Gibson cloning, producing Mirror reporter plasmids pAVA2965[GFP-ENE(WT)-mascRNA], pAVA2987[RFP-ENE(WT)-mascRNA], pAVA3000[GFP-ENE(C8351G)-mascRNA], pAVA2995[RFP-ENE(C8351G)-mascRNA]. *Guide RNA-expressing plasmids*: Plasmids expressing two sgRNAs were constructed by PCR-amplifying a 332-nucleotide fragment comprising the Cas9 scaffold and 7SK promoter from pAVA3129 (Supplemental Data S4), using a forward primer for sgRNA1 and a reverse primer for sgRNA2 (sequences in Supplemental Data S3), and cloned into BsmBI-linearized blasticidin-resistant lentiCRISPR vector^36^ to express first sgRNA from the U6 promoter and the second sgRNA from 7SK promoter. Plasmids expressing single guide RNAs (sequences in Supplemental Data S3) were constructed by cloning sgRNA sequences into BsmBI-linearized blasticidin-resistant lentiCRISPR vector^36^ using Gibson Assembly (New England Biolabs). The resulting lentiviral sgRNA-expressing constructs for gene knockout were sequences and named as follows: pAVA3250 for RBM7, pAVA3251 for SKIV2L2, pAVA3300 for ZCRB1, pAVA3258 for BRF2, pAVA3259 for RPP21, pAVA3586 for RPPH1, pAVA3318 for EXOSC10, pP18(T)_B9 for C1D, pAVA3791 as sgRNA1 for DDX59, pAVA3796 as sgRNA2 for DDX59, pAVA3317 for as sgRNA3 DDX59, pAVA3794 as sgRNA4 for DDX59, pP18(T)_C3 for EXOSC2, pP18(T)_C11 for EXOSC4, pP18(T)_B12 for EXOSC5, pP18(T)_B11 for EXOSC6, pP18(T)_C4 for EXOSC7, pP18(T)_B1 for EXOSC8, pP18(T)_A7 for EXOSC9, pP18(T)_A5 for DIS3, pAVA3773 for Control(-), pAVA3497 for POP1, pAVA3523 for RPP38, pAVA3545 for RPP29, pAVA3498 for POP5, pAVA3546 for RPP20, pAVA3507 for RPP14, pAVA3573 for RPP30, pAVA3540 for RPP40, pAVA3522 for RPP25, pAVA3548 for RPP25L. *Plasmids expressing full-length MALAT1*: Genomic DNA fragment consisting of (i) 1.2 kb of MALAT1 5′ flanking sequence, (ii) 8.8 kb sequence corresponding to MALAT1 lncRNA, and (iii) 0.7 kb of MALAT1 3′ flanking sequence was PCR-amplified from genomic DNA of the HeLa Fireworks cell line^35^, cloned into the BglII and XhoI restriction sites of the pcDNA5/FRT plasmid (Invitrogen) and verified by sequencing. An 11-nucleotide sequence (5′-cgctcgacgta-3′) was inserted 43 nucleotides upstream of the MALAT1 ENE sequence using the NheI restriction site to distinguish exogenously expressed MALAT1 from its endogenously expressed copies. C8351G mutation was introduced into the resulting 10.8 kb construct using PCR mutagenesis with primers listed in Supplemental Data S2. The resulting plasmids pAVA3169(MALAT1(C8351G)) and pAVA3171(MALAT1(WT)) are schematically shown in **Fig. 4g1**. Full sequences of all plasmids are shown in Supplemental Data S4; they will be deposited to the Addgene repository by the time of publication.

### Construction of stable cell lines

Stable orthogonal “Red” and “Green” Mirror cell lines shown in **Fig. S2** were obtained by exchanging FRT-integrated Fireworks NMD reporters^35^ in the “Green” Fireworks HeLa cell line AVAM526^35^ for Mirror reporters using transient co-transfection of pAVA2987 [RFP-ENE(WT)-mascRNA], pAVA3000 [GFP-ENE(C8351G)-mascRNA], pAVA2995 [RFP-ENE(C8351G)-mascRNA], and/or pAVA2965 [GFP-ENE(WT)-mascRNA] with the Flp recombinase-expressing plasmid pOG44 (Invitrogen). Single colonies of cells with stably integrated Mirror reporters were selected using hygromycin (150-300 μg/mL) and puromycin (0.08-0.16 μg/mL). Expanded colonies were FACS-sorted, propagated, and saved as Mirror “Red” AVAM712 (GFP-ENE(C8351G)-mascRNA, RFP-ENE(WT)-mascRNA) and “Green” AVAM742 (GFP-ENE(WT)-mascRNA, RFP-ENE(C8351G)-mascRNA) stable cell lines.

### Cell culture and maintenance

The “Red” Mirror cell line was maintained in DMEM (Gibco) media supplemented with 10% fetal bovine serum (FBS, Gibco), 100 U/mL penicillin-streptomycin (Corning), 150 μg/mL hygromycin (InvivoGen), and 0.08 μg/mL puromycin (InvivoGen). The “Green” Mirror cell line was maintained in DMEM media supplemented with 10% FBS, 100 U/mL penicillin-streptomycin, 300 μg/mL hygromycin, and 0.16 μg/mL puromycin. All fluorescent cell lines were routinely FACS-analyzed and sorted for stable GFP and RFP expression. HEK293T cells were maintained in DMEM supplemented with 10% FBS and 100 U/mL Penicillin-Streptomycin. All cell lines are authenticated using Short Tandem Repeat (STR) analysis and confirmed to be mycoplasma-free using PCR-based tests.

### Production of lentiviruses and transduction of human cell lines

To produce lentiviruses, sgRNA-expressing LentiCRISPR plasmids, listed in Supplemental Data S3, were co-transfected with the pCMV-dR8.91 packaging and pMD2.G envelope plasmids into the packaging HEK293T cells (AVAM761) using the TransIT-293 Transfection Reagent (Mirus Bio). Following daily media changes (DMEM, 30% FBS, 100 U/mL penicillin-streptomycin), lentivirus-containing media was collected 48 and 72 hours post-transfection, centrifuged at 900 g for 6 minutes to remove cellular debris, passed through a 0.45 μm filter, supplemented with 15 μg/mL polybrene (Millipore Sigma), and added to the Mirror cell lines growing at about 40% confluency. Selection for blasticidin-expressing infected cells was performed for 3 to 4 days using 3.0 μg/mL blasticidin (InvivoGen), which was added on the 3^rd^ day post-infection.

### Forward genetic screening using the Mirror approach

A single biological screen was performed in which total of 5x10^7^ cells (2 × 15 cm plates) of the “Red” “Mirror” cell line were transduced with blasticidin-resistant GeCKO-lentiCRISPR^36^ viral library and propagated in DMEM media supplemented with 10% FBS, 150 μg/mL hygromycin, 0.08 μg/mL puromycin and 3.0μg/mL blasticidin, with the blasticidin removed 4 days post-infection. Cell populations with increased fluorescence were FACS-isolated 10-11 days post-infection by sorting 360 million cells using the Bio-Rad S3e cell sorter, resulting in the screening each of the 64,751 sgRNAs in the GeCKO library^36^ on average 5,800 times. In FACS sorting and analysis, the gates were applied to exclude cell debris [SSC(area) vs FSC(area)], and cell doublets [FSC(width) vs FSC(hight)] (Supplemental Data S6). FACS data were collected using 488 nm and 561 nm excitation and 525/30 and 586/25 emission filters, for GFP and RFP fluorescence channels, respectively. The FACS-isolated cells were pelleted by centrifugation for 10 minutes at 1000g and frozen. Genomic DNA was isolated from FACS-isolated cells using phenol extraction. Sequences of guide RNAs were amplified from genomic DNA in two steps. First, a linear^64^ sgRNA amplification was performed using Herculase II DNA polymerase using 13 thermal cycles with a single sgRNA promoter-specific primer, RandomF (sequences of all primers are listed in Supplemental Data S2), and the following cycling parameters: 96°C 20s, 63°C 1min, 72°C 90s. Then, the second PCR primer, RandomR, was added to the reaction and a regular PCR was performed for 35 cycles as follows: 96°C 3min; 35 cycles of 96°C 20s, 63°C 1min, 72°C 45s; 72°C 10min; 4°C. The PCR-amplified pools of sgRNAs were Illumina-sequenced and/or cloned into a BsmBI-linearized, blasticidin-resistant lentiCRISPR vector^36^ to create an enriched lentiCRISPR sgRNA library for subsequent forward genetic screening rounds, as illustrated in **Fig. 3b**. The FACS-screening-enriched pool of sgRNAs and the pool of sgRNAs in the original GeCKO-lentiCRISPR library used for viral transduction were Illumina-sequenced. For each sgRNA in these pools, the enrichment coefficient was calculated as the ratio of sgRNA abundances after and before the screening. sgRNAs with low abundance in the original GeCKO-lentiCRISPR library (read count below 100) were excluded from ranking (Supplemental Data S1). Processing of deep sequencing data was performed as previously described^36^.

### In vivo analysis of RNA degradation using Actinomycin D

FACS-isolated cell populations of lentivirally-transduced cells expressing Cas9 and gene-specific sgRNAs (pAVA3317 targeting DDX59, pAVA3250 targeting RBM7, pAVA3251 targeting SKIV2L2, pAVA3318 targeting EXOSC10, and pAVA3773 representing negative control; sequences are provided in Supplemental Data S3) were seeded on 10 cm plates at 30% confluency. Eighteen hours later, each plate was transfected using TransIT-293 Transfection Reagent with 18 µg of the plasmid (pAVA3169, full sequence provided in Supplemental Data S4) expressing full-length lncRNA MALAT1 driven by its own promoter and carrying the ENE-destabilizing C8351G mutation. Twenty-four hours post-transfection, the cells were transferred to 6-well plates, recovered for 3 hours, and treated with 5 µg/mL of Actinomycin D for 0, 1, 2, and 3 hours, after which they were lysed using the TRIzol reagent and stored at -80°C for subsequent RNA purification. RNA was purified using TRI reagent (Zymo Research) according to manufacturer’s protocol and subjected to two 15-minute rounds of DNase I (Promega) treatment, phenol extraction, and ethanol precipitation. RT-qPCR quantification of RNA levels was performed as described below.

### RT-qPCR quantification of RNA levels

For each sample, 1 µg of total RNA was reverse-transcribed using iScript™ Reverse Transcription Supermix according to the manufacturer’s protocols. The resulting cDNA was diluted 20 times and qPCR-analysed using Applied Biosystems’ SYBR Green PCR Master Mix according to the manufacturer’s instructions for the CFX Opus 384 system and the following cycling conditions: 95°C for 3 min; 40 cycles of 95°C for 15 sec, 60°C for 30 sec. Relative quantification of transcript levels was performed using the ΔΔCT method and 18S RNA levels as the reference. Sequences of all primers are listed in Supplemental Data S2.

### Western blotting

Western blot of whole-cell lysates from CRISPR-knockout and control cells (obtained as shown in **Fig. 5a**; DDX59 mRNA knockout levels are shown in **Fig. 6g**) was performed by boiling samples in 2xLaemmli Sample Buffer (Bio-Rad), resolving them using 4-15% Criterion Precast Gels (Bio-Rad), transferring to Odyssey nitrocellulose membrane (Li-Cor) using TRIS-Glycine buffer (Bio-Rad) supplemented with 20% methanol, blocking the membrane with 5% non-fat dry milk in Tris Buffered Saline-Tween (Boston Bioproducts) for 2 hours at room temperature, incubating with an antibody (overnight at 4C for primary and two hours at room temperature for secondary) in blocking solution, developing using WesternBright ECL kit (Advansta), and imaging using G:Box iChemi XR5 (Syngene). Antibodies are described in Supplemental Data S5.

### Analysis of RNase P processing of the endogenous full-length lncRNA MALAT1

The levels of RNase P-unprocessed endogenous full-length lncRNA MALAT1 were measured by RT-qPCR using nested primers across the RNase P cleavage site of MALAT1-mascRNA. RNA extraction and reverse transcription were performed as described above. To quantify the unprocessed MALAT1 transcript, 20 pre-amplification cycles with outside primers (JD08-F and JD08-R; sequences in Supplemental Data S2) were conducted using Applied Biosystems’ SYBR Green PCR Master Mix with the PCR cycling parameters: 95°C for 3 min, followed by 20 cycles of 95°C for 15 s, and 60°C for 30 s. Each resulting pre-amplification reaction was diluted 40-fold with water and then used as a template for the subsequent qPCR reaction, which employed inside primers (JD07-F and JD07-R; sequences in Supplemental Data S2) and utilized Applied Biosystems’ SYBR Green PCR Master Mix with the PCR cycling parameters: 95°C for 3 min, followed by 40 cycles of 95°C for 15 s, and 60°C for 30 s. The levels of unprocessed MALAT1-mascRNA RNA were normalized to 18S RNA; ΔΔCt was used to quantify the relative expression levels. Data are presented as means of two biological x four technical replicates; error bars represent standard deviation. Statistically significant differences between knockout and control samples were determined by one-way ANOVA. Posthoc comparisons using Tukey’s HSD test were conducted to determine the overall difference between groups, and labeled as “*”, P<0.05; “**”, P<0.01; “***”, P<0.001. Positions of qPCR primers are indicated and their sequences are listed in Supplemental Data S2.

### RNA-Seq and genome mapping

Extracted RNA was diluted to 50ng/μl and sent for cDNA library construction (without the use of poly(A) selection) and paired-end sequencing (100-bp paired-end reads) following the manufacturer’s protocols (BGI-America, San Jose, CA). 60 millions clean reads received for each sample were mapped to human genome assembly GRCh38.p14 (GCA_000001405.29) using STAR^65^ (version 2.7.10b) with BAM output sorted by coordinate and standard output attributes. The resulting alignments were indexed using SAMtools^66^ (Version 1.18) and examined using IGV^67^ (version 2.16.1).

## DATA AVAILABILITY

The high-throughput sequencing data generated in this study, including CRISPR sgRNA library screening datasets, and RNA-Seq datasets have been deposited in NCBI Sequence Read Archive (SRA), accession number: PRJNA1186702. Source data are provided with this paper.

## Supporting information

Supplemental Data Descriptions

## ACKNOWLEDGMENTS

We thank Suzanne DeGregorio and Elizabeth Greif for help with cloning as well as Drs. Joan A. Steiz, David F. Clayton, Kazimierz Tycowski, and Raul Jobava for critical comments on the manuscript. This work was partially supported by funding to Joan A. Steitz from the Howard Hughes Medical Institute, and by National Institutes of Health grants HG009362 and GM139769 (project 1) to Andrei Alexandrov.

