## Supplemental Figures and Legends for "Identification of Human Pathways Acting on Nuclear Non-Coding RNAs Using the Mirror Forward Genetic Approach"

**Figure S1. The orthogonal "Green" Mirror reporter cell line for screening and rapid candidate validation. a.** Schematics of the "Green" Mirror reporter cell line, which is orthogonal to the "Red" Mirror reporter cell line. Reporters in the "Green" Mirror cell line differ from the reporters in the "Red" Mirror cell line (shown in **Fig**. **1e**) by just a single nucleotide (C8351 versus G8351), resulting in the orthogonal fluorescence shift upon pathway inhibition (as shown in **Figs**. **1f** and **S1b**). Together, the "Green" and "Red" Mirror cell lines provide robust controls against cell line- and fluorescent protein-specific effects. **b.** Anticipated effects of inhibiting pathways of nuclear degradation and processing on the fluorescence of "Green" Mirror cell line. **c.** Genome-wide iterative FACS-based forward genetic screening in the orthogonal "Green" Mirror reporter cell line. Respective screening in the "Red" Mirror cell line (**Fig**. **1e**) is shown in **Fig**. **3**. Unlike fluorescence of the "Red" reporter cell line, which shifts up, fluorescence of the "Green" reporter cell line shifts right upon stabilization of the ENE(C8351G) reporter or inhibition of the ENE-mascRNA cleavage. **d-h.** Individual transductions of distinct DDX59-targeting sgRNAs produce cell populations with increased RFP fluorescence in the orthogonal "Green" Mirror cell line. DDX59 knockouts with four different sgRNAs lead to increased RFP fluorescence, albeit to varying degrees, in the "Green" Mirror reporter cell line. This cell line is orthogonal to the "Red" reporter cell line (**Fig**. **1e**), in which it causes an increase in GFP fluorescence (**Figs**. **1f** and **4f**). Sequences of sgRNA are listed in Supplemental Data S3. **i.** The effects of knockout of Mirror-identified DDX59 on degradation of the full-length lncRNA MALAT1 with the wild-type ENE. RT-qPCR-derived MALAT1 levels were normalized to 18S RNA and are shown relative to the time of Actinomycin D addition. **j.** RT-qPCR analysis of the effects of knockouts of DDX59 on the levels of the endogenous wild-type lncRNA MALAT1. For RT-qPCRs, RNA levels were normalized to 18S RNA; $\Delta\Delta$Ct was used to quantify the relative expression levels. $\Delta\Delta$Ct was used to quantify the relative expression levels. Data are presented as means of two biological x four technical replicates; error bars represent standard deviation. Statistically significant differences between knockout and control samples were determined by one-way ANOVA. Posthoc comparisons using Tukey's HSD test were conducted to determine the overall difference between groups, and labeled as "*", $P<0.05$; "**", $P<0.01$; "***", $P<0.001$. Sequences of qPCR primers are listed in Supplemental Data S2.

**Figure S2. Schematics of the genome-integrated Mirror reporters in the "Red" and "Green" Mirror stable cell lines.** To generate each Mirror cell line, two Mirror reporters were integrated at two separate FRT sites in the precursor "Green" Fireworks cell line[35] using FRT cassette exchange mediated by Flp recombinase. Note that puromycin N-acetyl-transferase (PAC) expressed by the FRT-integrated GFP-expressing Mirror cassettes confers puromycin resistance in combination with reporters expressing TEV protease. This is due to a PAC-inhibiting N-terminal peptide produced as a result of transcription and translation of the nucleotide sequence of the FTR site, required for cassette insertion. In the Mirror cell lines, this inhibitory N-terminal peptide is cleaved by the reporter-expressed TEV protease, yielding an active PAC enzyme that provides puromycin resistance.

**Figure S3. DDX59 knockout results in little, if any, retention of U2 introns, but pronounced retention of U12 introns in genes associated with cilia, ciliogenesis, and Oral-Facial-Digital syndrome. a-c.** Integrated Genome Viewer (IGV) traces of RNA-seq reads for mRNAs of general housekeeping genes β-actin (ACTB, shown in panel a), β-tubulin (shown in panel b), and β-2-microglobulin (B2M, shown in panel c), illustrating little to no observable retention of major introns. **d-i.** IGV traces of RNA-seq reads for mRNAs of minor intron-containing genes associated with cilia or ciliary function, ARMC9, KATNIP, SIF1, CUL1, ACTR10, and CCDC28B, illustrating significant retention of minor introns. Cell populations with knockouts of DDX59, RBM7, ZCRB1, and EXOSC10 were obtained as shown in **Fig**. **5a** and their respective IGV traces are colored here accordingly. Major and minor introns are labeled as U2 and U12, respectively.

**Figure S4. DDX59 knockout results in pronounced retention of U12 introns in genes associated with cilia, ciliogenesis, Oral-Facial-Digital syndrome (OFD), and RNA Exosome. a-f.** IGV traces of RNA-seq reads for mRNAs of minor intron-containing genes associated with cilia or ciliary function: PPP5C, TCTN3, ACTL6A, TMEM107, KIFAP3, and RABL2A. Two of these genes, TCTN3 and TMEM107, are additionally associated with Oral-Facial-Digital syndrome types OFDIV and OFDVI, respectively. Cell populations with knockouts of DDX59, RBM7, ZCRB1, and EXOSC10 were obtained as shown in **Fig**. **5a** and their respective IGV traces are colored here accordingly. Major and minor introns are labeled as U2 and U12, respectively. Panel b additionally shows RT-qPCR quantification of intron retention in mRNA of the OFD-associated TCTN3 after DDX59 knockout. All qPCR quantifications and statistical tests were performed as described in the legend of **Fig**. **5**; qPCR primers are listed in Supplemental Data S2. **g.** Integrated Genome Viewer (IGV) traces of RNA-seq reads for minor intron-containing gene, the exosome components EXOSC5. Cell populations with DDX59, RBM7, ZCRB1, and EXOSC10 knockouts were obtained as shown in **Fig**. **5a** and their respective IGV traces are colored accordingly. Major and minor introns are labeled as U2 and U12, respectively.

**Figure S5. DDX59 deficiency reduces protein levels of the endogenous RNA exosome component EXOSC1, the NEXT complex component ZCCHC8, and the minor intron-containing TCTN3 minigene. a.** Western blot with 2-fold serial dilutions of whole-cell lysates from CRISPR-knockout and control cells (obtained as shown in **Fig**. **5a**; DDX59 mRNA knockout levels are quantified in **Fig**. **6g**). Knockouts of EXOSC1 and ZCCHC8 are included to show specificity of the respective antibodies (detailed in Supplemental Data S5). **b.** Western blot with 2-fold serial dilutions of whole-cell lysates from DDX59 CRISPR-knockout and control cells (obtained as shown in **Fig**. **5a**; DDX59 mRNA knockout levels are quantified in **Fig**. **6g**) probed with anti-c-Myc antibody (detailed in Supplemental Data S5). A plasmid expressing the minor intron-containing TCTN3 minigene (under control of the EF1α promoter), consisting of (N- to C-terminus): 6xMyc tag, exon 10 of TCTN3, minor intron 10 of TCTN3, exon 11 of TCTN3, GFP, and 3xFLAG tag, was co-transfected with a control plasmid expressing 6xMyc-tagged control protein CWC22 into DDX59 knockout and control cells. A no 6xMyc-CWC22 control, no TCTN3 minigene control, and untransfected control are included to confirm the identity of the respective Western blot bands.

**Figure S6. Knockout of DDX59 leads to extensive retention of U12 introns in the mRNA of Derlin 2, a protein that mediates the retro-translocation of the Sonic Hedgehog (SHH) at the endoplasmic reticulum, providing a potential mechanism for SHH pathway deficiency observed in DDX59-associated OFD. a.** IGV traces of RNA-seq reads for DERL2 mRNA. Cell populations with knockouts of DDX59, RBM7, ZCRB1, and EXOSC10 were obtained as shown in **Fig**. **5a** and their respective IGV traces are colored here accordingly. **b.** RT-qPCR quantification of intron retention in mRNA of DERL2. RNA levels were normalized to 18S RNA; $\Delta\Delta$Ct was used to quantify the relative expression levels. Data are presented as means of two biological x four technical replicates; error bars represent standard deviation. Statistically significant differences between knockout and control samples were determined by one-way ANOVA. Posthoc comparisons using Tukey's HSD test were conducted to determine the overall difference between groups, and labeled as “*”, P<0.05; “**”, P<0.01; “***”, P<0.001. Positions of qPCR primers are indicated and their sequences are listed in Supplemental Data S2.

### Figure S1

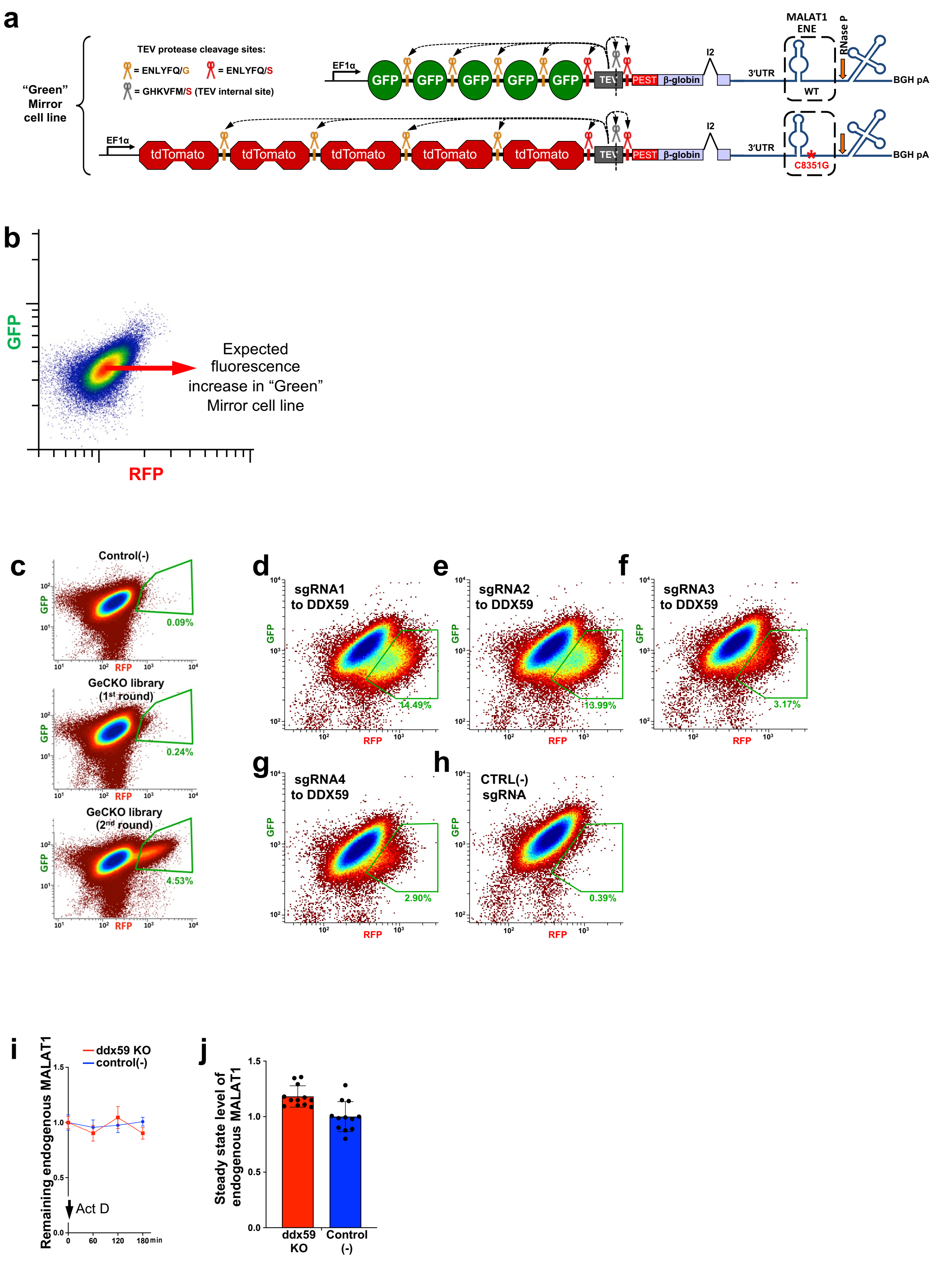
a
"Green" Mirror cell line
TEV protease cleavage sites:
= ENLYFQ/G
= ENLYFQ/S
= GHKVFM/S (TEV internal site)
EF1α
GFP
TEV
PEST
β-globin
I2
3'UTR
MALAT1 ENE
WT
RNase P
BGH pA
tdTomato
C8351G
b
GFP
RFP
Expected fluorescence increase in "Green" Mirror cell line
c
Control(-)
0.09%
GeCKO library (1st round)
0.24%
GeCKO library (2nd round)
4.53%
d
sgRNA1 to DDX59
14.49%
e
sgRNA2 to DDX59
13.99%
f
sgRNA3 to DDX59
3.17%
g
sgRNA4 to DDX59
2.90%
h
CTRL(-) sgRNA
0.39%
i
ddx59 KO
control(-)
Remaining endogenous MALAT1
Act D
min
j
Steady state level of endogenous MALAT1
ddx59 KO
Control (-)

### Figure S2

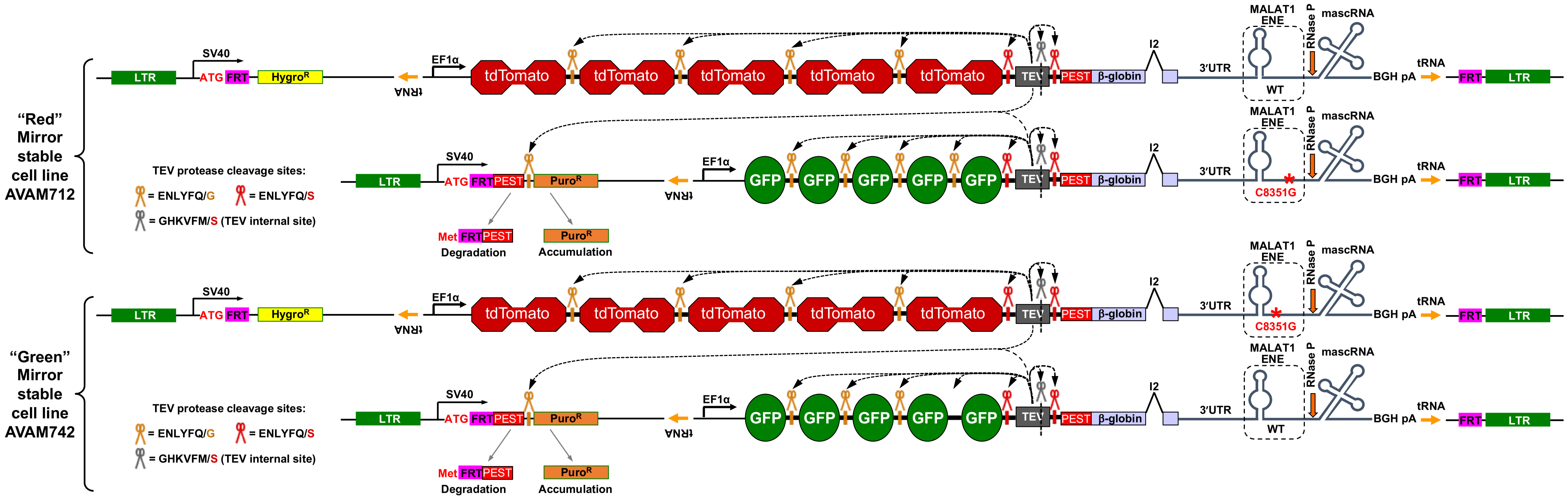

"Red" Mirror stable cell line AVAM712
"Green" Mirror stable cell line AVAM742
LTR
SV40
ATG
FRT
HygroR
tRNA
EF1α
tdTomato
TEV
PEST
β-globin
I2
3'UTR
MALAT1 ENE
WT
C8351G
RNase P
mascRNA
BGH pA
GFP
PuroR
Met FRT PEST
Degradation
Accumulation
TEV protease cleavage sites:
= ENLYFQ/G
= ENLYFQ/S
= GHKVFM/S (TEV internal site)

### Figure S3

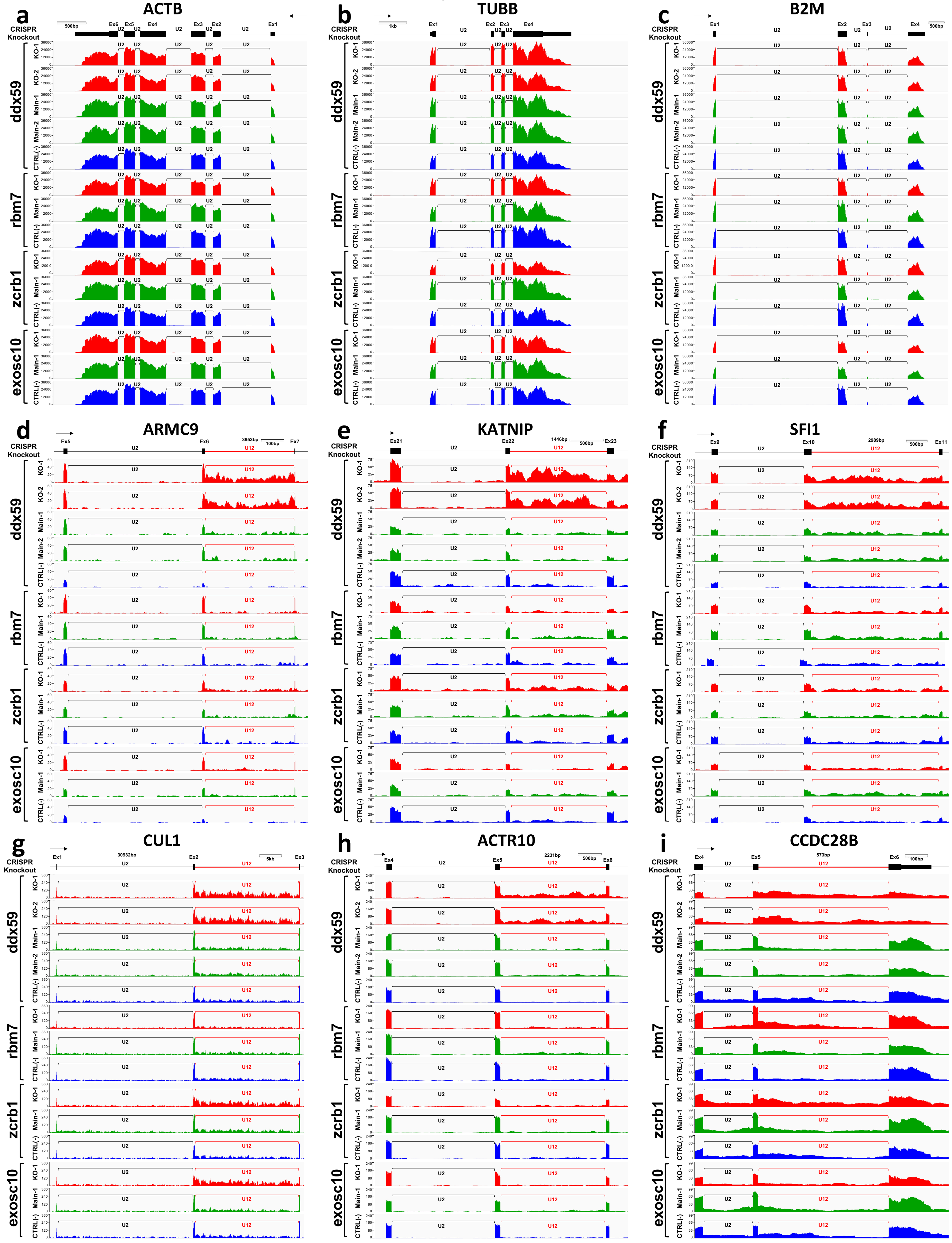
a
ACTB
b
TUBB
c
B2M
d
ARMC9
e
KATNIP
f
SFI1
g
CUL1
h
ACTR10
i
CCDC28B
CRISPR
Knockout
ddx59
rbm7
zcrb1
exosc10
KO-1
KO-2
Main-1
Main-2
CTRL(-)
U2
U12

### Figure S4

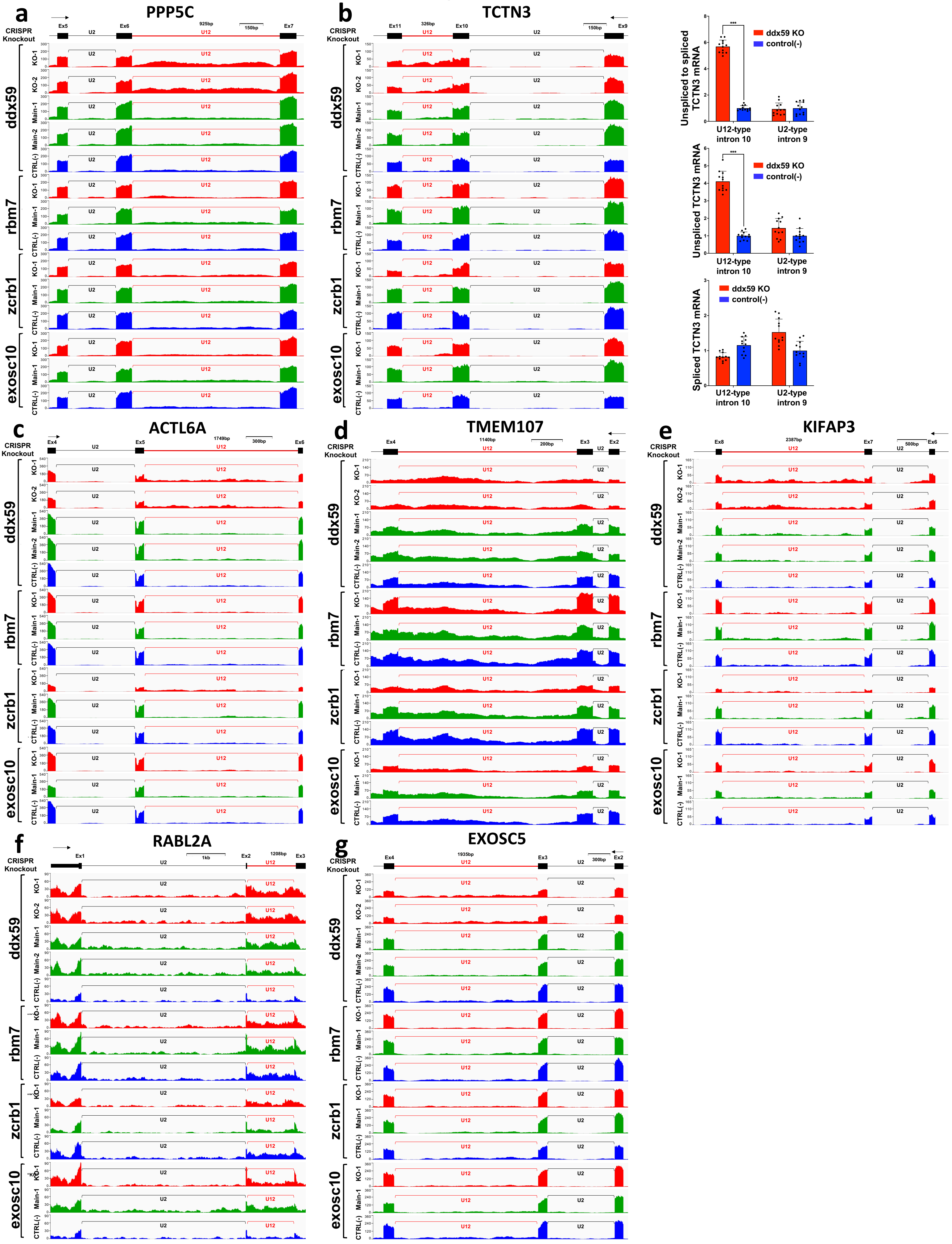

a
PPP5C
b
TCTN3
c
ACTL6A
d
TMEM107
e
KIFAP3
f
RABL2A
g
EXOSC5
CRISPR Knockout
ddx59
rbm7
zcrb1
exosc10
KO-1
KO-2
Main-1
Main-2
CTRL(-)
U2
U12
Unspliced to spliced TCTN3 mRNA
Unspliced TCTN3 mRNA
Spliced TCTN3 mRNA
ddx59 KO
control(-)
U12-type intron 10
U2-type intron 9
***

### Figure S5

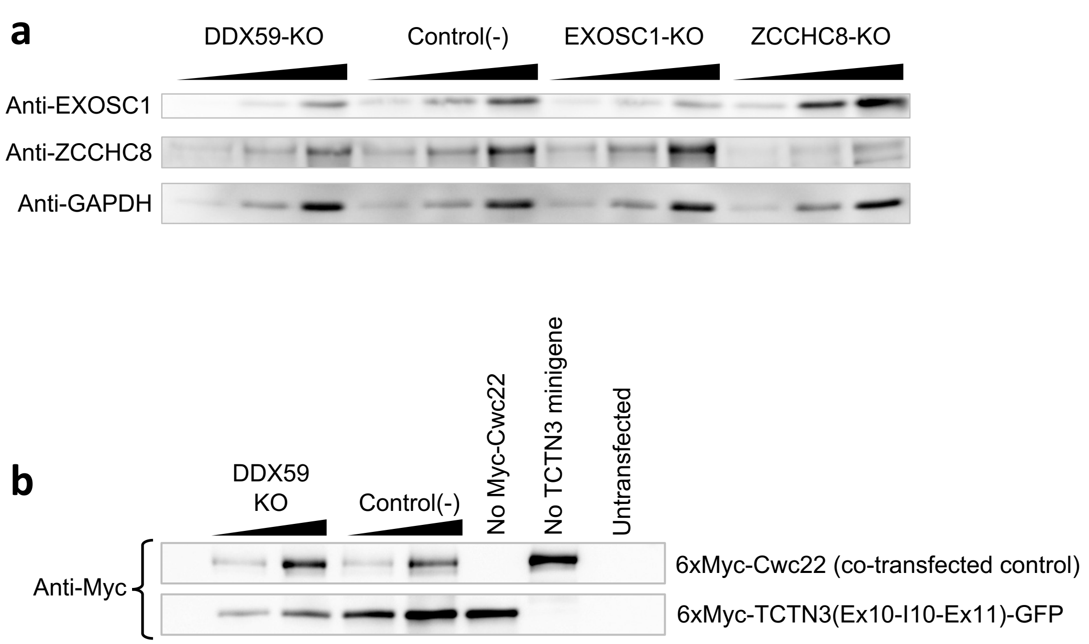
a
DDX59-KO
Control(-)
EXOSC1-KO
ZCCHC8-KO
Anti-EXOSC1
Anti-ZCCHC8
Anti-GAPDH
b
DDX59
KO
Control(-)
No Myc-Cwc22
No TCTN3 minigene
Untransfected
Anti-Myc
6xMyc-Cwc22 (co-transfected control)
6xMyc-TCTN3(Ex10-I10-Ex11)-GFP

### Figure S6

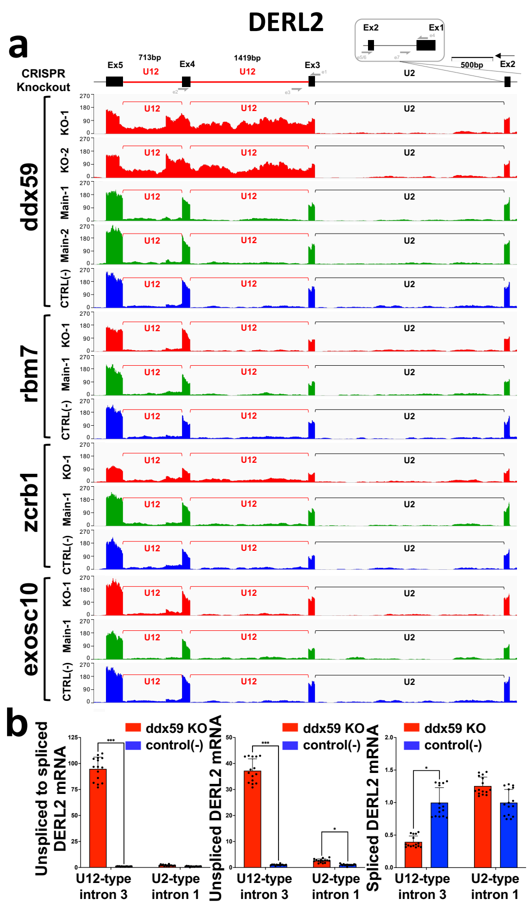
a
DERL2
CRISPR Knockout
Ex5
713bp
U12
Ex4
1419bp
U12
Ex3
U2
Ex2
Ex1
500bp
ddx59
KO-1
KO-2
Main-1
Main-2
CTRL(-)
rbm7
zcrb1
exosc10
b
ddx59 KO
control(-)
Unspliced to spliced DERL2 mRNA
Unspliced DERL2 mRNA
Spliced DERL2 mRNA
U12-type intron 3
U2-type intron 1
