## Supplemental Data Descriptions for "Identification of Human Pathways Acting on Nuclear Non-Coding RNAs Using the Mirror Forward Genetic Approach"

### DESCRIPTION OF SUPPLEMENTAL DATA

**Supplemental Data S1. Sequences of guide RNAs in the Mirror screening-enriched library**

**Supplemental Data S2. Sequences of Cloning Oligonucleotides, DNA Oligonucleotides used in qPCR, and Sequencing Primers**

**Supplemental Data S3. Sequences of guide RNAs for CRISPR knockouts**

**Supplemental Data S4. Plasmid Sequences**

Sequences of Mirror reporters:

pAVA2987 [RFP-ENE(WT)-mascRNA]

pAVA3000 [GFP-ENE(C8351G)-mascRNA]

pAVA2995 [RFP-ENE(C8351G)-mascRNA]

pAVA2965 [GFP-ENE(WT)-mascRNA]

pAVA3871 [GFP-ENE(C8351G)-mascRNA(mut 8356-8370)]

pAVA3874 [GFP-ENE(WT)-mascRNA(mut 8356-8370)]

Sequences of plasmids expressing full-length MALAT1:

pAVA3169 MALAT1(C8351G)

pAVA3171 MALAT1(WT)

Sequences of plasmids expressing two sgRNAs:

pAVA3129 [U6-sgRNA1, 7SK-sgRNA2]

Sequence of plasmid expressing minor intron-containing TCTN3 minigene:

pAVA3939 [minor intron-containing TCTN3 minigene (EFS-6xMyc-TCTN3-Ex10-I10-Ex11-copGFP-3xFLAG)]

**Supplemental Data S5. Antibodies**

**Supplemental Data S6. Schematics of the employed FACS gates.**
