## Supplemental Data S4 for "Identification of Human Pathways Acting on Nuclear Non-Coding RNAs Using the Mirror Forward Genetic Approach"

### Supplemental Data File S4: Supplemental vector sequences, related to Experimental Procedures

#### A) pAVA2987 [RFP-ENE(WT)-mascRNA, Hygromycin$^R$]

**tDNA1: 1045-1141**
**EF1alfa promoter: 1176-2679**
**5xtdTomato: 2702-9994**
**TEV protease: 9995-10720**
**PEST (degradation sequence): 10748-10882**
**Human beta-globin(ΔI1): 10889-12271**
**MALAT1-ENE(WT)-mascRNA: 12278-12437**
**BGH polyA: 12470-12694**
**tDNA2: 12980-13076**
**FRT: 13087-13113**
**Hygromycin$^R$ (lacks promoter and the 1$^{st}$ Methionine): 13154-14430**

acttcattttaatttaaaaggatctaggtgaagatcctttttgataatctcatgaccaaaatcccttaacgtgagt
tttcgttccactgagcgtcagaccccgtagaaaagatcaaaggatcttcttgagatcctttttttctgcgcgtaatc
tgctgcttgcaaacaaaaaaaccaccgctaccagcggtggtttgtttgccggatcaagagctaccaactctttttcc
gaaggtaactggcttcagcagagcgcagataccaaatactgtccttctagtgtagccgtagttaggccaccacttca
agaactctgtagcaccgcctacatacctcgctctgctaatcctgttaccagtggctgctgccagtggcgataagtcg
tgtcttaccgggttggactcaagacgatagttaccggataaggcgcagcggtcgggctgaacggggggttcgtgcac
acagcccagcttggagcgaacgacctacaccgaactgagatacctacagcgtgagctatgagaaagcgccacgcttc
ccgaagggagaaaggcggacaggtatccggtaagcggcagggtcggaacaggagagcgcacgagggagcttccaggg
ggaaacgcctggtatctttatagtcctgtcgggtttcgccacctctgacttgagcgtcgatttttgtgatgctcgtc
aggggggcggagcctatggaaaaacgccagcaacgcggcctttttacggttcctggccttttgctggccttttgctc
acatgttctttcctgcgttatcccctgattctgtggataaccgtattaccgcctttgagtgagctgataccgctcgc
cgcagccgaacgaccgagcgcagcgagtcagtgagcgaggaagcggaagagcgcccaatacgcaaaccgcctctccc
cgcgcgttggccgattcattaatgcagctggcacgacaggtttcccgactggaaagcgggcagtgagcgcaacgcaa
ttaatgtgagttagctcactcattaggcaccccaggcACGCGAagcgttgttggtttgtagtgcccggtttcgaacc
ggggacctttcgcgtgttaggcgaacgtgataaccactacactacggaaaccaacggtgctagACGCGTattgtcta
ttctgactcggatcCGTACGaattctcatgtttgacagcttatcatcgattagctttggagctaagccagcaatggt
agagggaagattctgcacgtcccttccaggcggcctccccgtcaccaccccccccaacccgccccgaccggagctga
gagtaattcatacaaaggactcgcccctgccttggggaatcccagggaccgtcgttaaactcccactaacgtagaa
cccagagatcgctgcgttcccgcccctcacccgcccgctctcgtcatcactgaggtggagaagagcatgcgtgagg
ctccggtgcccgtcagtgggcagagcgcacatcgcccacagtccccgagaagttggggggaggggtcggcaattgaa
ccggtgcctagagaaagtggcgcggggtaaactgggaaagtgatgtcgtgtactggctccgcctttttcccgagggt
gggggagaaccgtatataagtgcagtagtcgccgtgaacgttctttttcgcaacgggtttgccgccagaacacaggt
aagtgccgtgtgtggttcccgcgggcctggcctctttacgggttatggcccttgcgtgccttgaattacttccacgc
ccctggctgcagtacgtgattcttgatcccgagcttcgggttggaagtgggtgggagagttcgaggccttgcgctta
aggagccccttcgcctcgtgcttgagttgaggcctggcctgggcgctggggccgccgcgtgcgaatctggtggcacc
ttcgcgcctgtctcgctgctttcgataagtctctagccatttaaaatttttgatgacctgctgcgacgctttttttc
tggcaagatagtcttgtaaatgcgggccaagatctgcacactggtatttcggtttttggggccgcgggcggcgacgg
ggcccgtgcgtcccagcgcacatgttcggcgaggcggggcctgcgagcgcggccaccgagaatcggacgggggtagt
ctcaagctggccggcctgctctggtgcctggcctcgcgccgccgtgtatcgccccgccctgggcggcaaggctgg
cccggtcggcaccagttgcgtgagcggaaagatggccgcttcccggccctgctgcagggagctcaaaatggaggacg
cggcgctcgggagagcgggcgggtgagtcacccacacaaaggaaaagggcctttccgtcctcagccgtcgcttcatg
tgactccacggagtaccgggcgccgtccaggcacctcgattagttctcgagcttttggagtacgtcgtctttaggtt
ggggggaggggttttatgcgatggagtttccccacactgagtgggtggagactgaagttaggccagcttggcacttg
atgtaattctccttggaatttgccctttttgagtttggatcttggttcattctcaagcctcagacagtggttcaaag
tttttttcttccatttcaggtgtcgtgaaaactctagcgtttccgctctagaactagtccaGCTAGCGCTACCGGTC
GCCACCatggtgagcaagggcgaggaggtcatcaaagagttcatgcgcttcaaggtgcgcatggagggctccatgaa

cggccacgagttcgagatcgagggcgagggcgagggccgcccctacgagggcacccagaccgccaagctgaaggtga
ccaagggcggcccccctgcccttcgcctgggacatcctgtcccccagttcatgtacggctccaaggcgtacgtgaag
caccccgccgacatccccgattacaagaagctgtccttccccgagggcttcaagtgggagcgcgtgatgaacttcga
ggacggcggtctggtgaccgtgacccaggactcctccctgcaggacggcacgctgatctacaaggtgaagatgcgcg
gcaccaacttccccccgacggccccgtaatgcagaagaagaccatgggctgggaggcctccaccgagcgcctgtac
cccgcgacggcgtgctgaagggcgagatccaccaggccctgaagctgaaggacggcggccactacctggtggagtt
caagaccatctacatggccaagaagcccgtgcaactgcccggctactactacgtggacaccaagctggacatcacct
cccacaacgaggactacaccatcgtggaacagtacgagcgctccgagggccgccaccacctgttcctggggcatggc
accggcagcaccggcagcggcagctccggcaccgcctcctccgaggacaacaacatggccgtcatcaaagagttcat
gcgcttcaaggtgcgcatggagggctccatgaacggccacgagttcgagatcgagggcgagggcgagggccgcccct
acgagggcacccagaccgccaagctgaaggtgaccaagggcggcccccctgcccttcgcctgggacatcctgtccccc
cagttcatgtacggctccaaggcgtacgtgaagcaccccgccgacatccccgattacaagaagctgtccttccccga
gggcttcaagtgggagcgcgtgatgaacttcgaggacggcggtctggtgaccgtgacccaggactcctccctgcagg
acggcacgctgatctacaaggtgaagatgcgcggcaccaacttccccccgacggccccgtaatgcagaagaagacc
atgggctgggaggcctccaccgagcgcctgtaccccgcgacggcgtgctgaagggcgagatccaccaggccctgaa
gctgaaggacggcggccactacctggtggagttcaagaccatctacatggccaagaagcccgtgcaactgcccggct
actactacgtggacaccaagctggacatcacctcccacaacgaggactacaccatcgtggaacagtacgagcgctcc
gagggccgccaccacctgttcctgtacggcatggacgagctgtacaag**GTcactCcgtgT**gaaaacctgtacttcca
aggt**atg**gtgagcaagggcgaggaggtcatcaaagagttcatgcgcttcaaggtgcgcatggagggctccatgaacg
gccacgagttcgagatcgagggcgagggcgagggccgcccctacgagggcacccagaccgccaagctgaaggtgacc
aagggcggcccccctgcccttcgcctgggacatcctgtcccccagttcatgtacggctccaaggcgtacgtgaagca
ccccgccgacatccccgattacaagaagctgtccttccccgagggcttcaagtgggagcgcgtgatgaacttcgagg
acggcggtctggtgaccgtgacccaggactcctccctgcaggacggcacgctgatctacaaggtgaagatgcgcggc
accaacttccccccgacggccccgtaatgcagaagaagaccatgggctgggaggcctccaccgagcgcctgtaccc
ccgcgacggcgtgctgaagggcgagatccaccaggccctgaagctgaaggacggcggccactacctggtggagttca
agaccatctacatggccaagaagcccgtgcaactgcccggctactactacgtggacaccaagctggacatcacctcc
cacaacgaggactacaccatcgtggaacagtacgagcgctccgagggccgccaccacctgttcctggggcatggcac
cggcagcaccggcagcggcagctccggcaccgcctcctccgaggacaacaacatggccgtcatcaaagagttcatgc
gcttcaaggtgcgcatggagggctccatgaacggccacgagttcgagatcgagggcgagggcgagggccgcccctac
gagggcacccagaccgccaagctgaaggtgaccaagggcggcccccctgcccttcgcctgggacatcctgtcccccca
gttcatgtacggctccaaggcgtacgtgaagcaccccgccgacatccccgattacaagaagctgtccttccccgagg
gcttcaagtgggagcgcgtgatgaacttcgaggacggcggtctggtgaccgtgacccaggactcctccctgcaggac
ggcacgctgatctacaaggtgaagatgcgcggcaccaacttccccccgacggccccgtaatgcagaagaagaccat
gggctgggaggcctccaccgagcgcctgtaccccgcgacggcgtgctgaagggcgagatccaccaggccctgaagc
tgaaggacggcggccactacctggtggagttcaagaccatctacatggccaagaagcccgtgcaactgcccggctac
tactacgtggacaccaagctggacatcacctcccacaacgaggactacaccatcgtggaacagtacgagcgctccga
gggccgccaccacctgttcctgtacggcatggacgagctgtacaag**ACcactGcgtgC**gaaaacctgtacttccaag
gt**atg**gtgagcaagggcgaggaggtcatcaaagagttcatgcgcttcaaggtgcgcatggagggctccatgaacggc
cacgagttcgagatcgagggcgagggcgagggccgcccctacgagggcacccagaccgccaagctgaaggtgaccaa
gggcggcccccctgcccttcgcctgggacatcctgtcccccagttcatgtacggctccaaggcgtacgtgaagcacc
ccgccgacatccccgattacaagaagctgtccttccccgagggcttcaagtgggagcgcgtgatgaacttcgaggac
ggcggtctggtgaccgtgacccaggactcctccctgcaggacggcacgctgatctacaaggtgaagatgcgcggcac
caacttccccccgacggccccgtaatgcagaagaagaccatgggctgggaggcctccaccgagcgcctgtaccccc
gcgacggcgtgctgaagggcgagatccaccaggccctgaagctgaaggacggcggccactacctggtggagttcaag
accatctacatggccaagaagcccgtgcaactgcccggctactactacgtggacaccaagctggacatcacctccca
caacgaggactacaccatcgtggaacagtacgagcgctccgagggccgccaccacctgttcctggggcatggcaccg
gcagcaccggcagcggcagctccggcaccgcctcctccgaggacaacaacatggccgtcatcaaagagttcatgcgc
ttcaaggtgcgcatggagggctccatgaacggccacgagttcgagatcgagggcgagggcgagggccgcccctacga
gggcacccagaccgccaagctgaaggtgaccaagggcggcccccctgcccttcgcctgggacatcctgtcccccagt
tcatgtacggctccaaggcgtacgtgaagcaccccgccgacatccccgattacaagaagctgtccttccccgagggc
ttcaagtgggagcgcgtgatgaacttcgaggacggcggtctggtgaccgtgacccaggactcctccctgcaggacgg
cacgctgatctacaaggtgaagatgcgcggcaccaacttccccccgacggccccgtaatgcagaagaagaccatgg
gctgggaggcctccaccgagcgcctgtaccccgcgacggcgtgctgaagggcgagatccaccaggccctgaagctg
aaggacggcggccactacctggtggagttcaagaccatctacatggccaagaagcccgtgcaactgcccggctacta
ctacgtggacaccaagctggacatcacctcccacaacgaggactacaccatcgtggaacagtacgagcgctccgagg
gccgccaccacctgttcctgtacggcatggacgagctgtacaag**GTcactAcgtgT**gaaaacctgtacttccaaggt
**atg**gtgagcaagggcgaggaggtcatcaaagagttcatgcgcttcaaggtgcgcatggagggctccatgaacggcca

cgagttcgagatcgagggcgagggcgagggccgcccctacgagggcacccagaccgccaagctgaaggtgaccaagg
gcggcccccctgcccttcgcctgggacatcctgtcccccagttcatgtacggctccaaggcgtacgtgaagcacccc
gccgacatccccgattacaagaagctgtccttccccgagggcttcaagtgggagcgcgtgatgaacttcgaggacgg
cggtctggtgaccgtgacccaggactcctccctgcaggacggcacgctgatctacaaggtgaagatgcgcggcacca
acttcccccccgacggccccgtaatgcagaagaagaccatgggctgggaggcctccaccgagcgcctgtacccccgc
gacggcgtgctgaagggcgagatccaccaggccctgaagctgaaggacggcggccactacctggtggagttcaagac
catctacatggccaagaagcccgtgcaactgcccggctactactacgtggacaccaagctggacatcacctcccaca
acgaggactacaccatcgtggaacagtacgagcgctccgagggccgccaccacctgttcctggggcatggcaccggc
agcaccggcagcggcagctccggcaccgcctcctccgaggacaacaacatggccgtcatcaaagagttcatgcgctt
caaggtgcgcatggagggctccatgaacggccacgagttcgagatcgagggcgagggcgagggccgcccctacgagg
gcacccagaccgccaagctgaaggtgaccaagggcggcccccctgcccttcgcctgggacatcctgtcccccagttc
atgtacggctccaaggcgtacgtgaagcaccccgccgacatccccgattacaagaagctgtccttccccgagggctt
caagtgggagcgcgtgatgaacttcgaggacggcggtctggtgaccgtgacccaggactcctccctgcaggacggca
cgctgatctacaaggtgaagatgcgcggcaccaacttcccccccgacggccccgtaatgcagaagaagaccatgggc
tgggaggcctccaccgagcgcctgtacccccgcgacggcgtgctgaagggcgagatccaccaggccctgaagctgaa
ggacggcggccactacctggtggagttcaagaccatctacatggccaagaagcccgtgcaactgcccggctactact
acgtggacaccaagctggacatcacctcccacaacgaggactacaccatcgtggaacagtacgagcgctccgagggc
cgccaccacctgttcctgtacggcatggacgagctgtacaagACcactTcgtgCgaaaacctgtacttccaaggtat
ggtgagcaagggcgaggaggtcatcaaagagttcatgcgcttcaaggtgcgcatggagggctccatgaacggccacg
agttcgagatcgagggcgagggcgagggccgcccctacgagggcacccagaccgccaagctgaaggtgaccaagggc
ggcccccctgcccttcgcctgggacatcctgtcccccagttcatgtacggctccaaggcgtacgtgaagcaccccgc
cgacatccccgattacaagaagctgtccttccccgagggcttcaagtgggagcgcgtgatgaacttcgaggacggcg
gtctggtgaccgtgacccaggactcctccctgcaggacggcacgctgatctacaaggtgaagatgcgcggcaccaac
ttcccccccgacggccccgtaatgcagaagaagaccatgggctgggaggcctccaccgagcgcctgtacccccgcga
cggcgtgctgaagggcgagatccaccaggccctgaagctgaaggacggcggccactacctggtggagttcaagacca
tctacatggccaagaagcccgtgcaactgcccggctactactacgtggacaccaagctggacatcacctcccacaac
gaggactacaccatcgtggaacagtacgagcgctccgagggccgccaccacctgttcctggggcatggcaccggcag
caccggcagcggcagctccggcaccgcctcctccgaggacaacaacatggccgtcatcaaagagttcatgcgcttca
aggtgcgcatggagggctccatgaacggccacgagttcgagatcgagggcgagggcgagggccgcccctacgagggc
acccagaccgccaagctgaaggtgaccaagggcggcccccctgcccttcgcctgggacatcctgtcccccagttcat
gtacggctccaaggcgtacgtgaagcaccccgccgacatccccgattacaagaagctgtccttccccgagggcttca
agtgggagcgcgtgatgaacttcgaggacggcggtctggtgaccgtgacccaggactcctccctgcaggacggcacg
ctgatctacaaggtgaagatgcgcggcaccaacttcccccccgacggccccgtaatgcagaagaagaccatgggctg
ggaggcctccaccgagcgcctgtacccccgcgacggcgtgctgaagggcgagatccaccaggccctgaagctgaagg
acggcggccactacctggtggagttcaagaccatctacatggccaagaagcccgtgcaactgcccggctactactac
gtggacaccaagctggacatcacctcccacaacgaggactacaccatcgtggaacagtacgagcgctccgagggccg
ccaccacctgttcctgtacggcatggacgagctgtacaaggaaaacctgtacttccaaAgtggagaAAGCTTgttta
agggaccacgtgattacaacccgatatcgagcaccatttgtcatttgacgaatgaatctgatgggcacacaacatcg
ttgtatggtattggatttggtcccttcatcattacaaacaagcacttgtttagaagaaataatggaacactgttggt
ccaatcactacatggtgtattcaaggtcaagaacaccacgactttgcaacaacacctcattgatgggagggacatga
taattattcgcatgcctaaggatttcccaccatttcctcaaaagctgaaatttagagagccacaaagggaagagcgc
atatgtcttgtgacaaccaacttccaaactaagagcatgtctagcatggtgtcagacactagttgcacattcccttc
atctgatggcatattctggaagcattggattcaaaccaaggatgggcagtgtggcagtccattagtatcaactagag
atgggttcattgttggtatacactcagcatcgaatttcaccaacacaaacaattatttcacaagcgtgccgaaaaac
ttcatggaattgttgacaaatcaggaggcgcagcagtgggttagtggttggcgattaaatgctgactcagtattgtg
gggggccataaagttttcatgagcaaacctgaagagccttttcagccagttaaggaagcgactcaactcatgaatg
aattggtgtactcgcaagaaaacctgtacttccaaagtATGCATAGATCACGAGATATCAGCCATGGCTTCCCGCCG
GCGGTGGCGGCGCAGGATGATGGCACGCTGCCCATGTCTTGTGCCCAGGAGAGCGGGATGGACCGTCACCCTGCAGC
CTGTGCTTCTGCTAGGATCAATGTGGAGCTCGTGCACCTGACTCCTGAGGAGAAGTCTGCCGTTACTGCCCTGTGGG
GCAAGGTGAACGTGGAT**GAAGTTGGTGGTGAGGCCCTG**GGCAGGCTGCTGGTGGTCTACCCTTGGACCCAGAGGTTC
TTTGAGTCCTTTGGGGATCTGTCCACTCCTGATGCTGTTATGGGCAACCCTAAGGTGAAGGCTCATGGCAAGAAAGT
GCTCGGTGCCTTTAGTGATGGCCTGGCTCACCTGGACAACCTCAAGGGCACCTTTGCCACACTGAGTGAGCTGCACT
GTGACAAGCTGCACGTGGATCCTGAGAACTTCAGGGTGAGTCTATGGGACCCTTGATGTTTTCTTTCCCCTTCTTTT
CTATGGTTAAGTTCATGTCATAGGAAGGGGATAAGTAACAGGGTACAGTTTAGAATGGGAAACAGACGAATGATTGC
ATCAGTGTGGAAGTCTCAGGATCGTTTTAGTTTCTTTTATTTGCTGTTCATAACAATTGTTTTCTTTTGTTTAATTC
TTGCTTTCTTTTTTTTTCTTCTCCGCAATTTTTACTATTATACTTAATGCCTTAACATTGTGTATAACAAAAGGAAA
TATCTCTGAGATACATTAAGTAACTTAAAAAAAAAACTTTACACAGTCTGCCTAGTACATTACTATTTGGAATATAT

GTGTGCTTATTTGCATATTCATAATCTCCCTACTTTATTTTCTTTTATTTTTAATTGATACATAATCATTATACATA
TTTATGGGTTAAAGTGTAATGTTTTAATATGTGTACACATATTGACCAAATCAGGGTAATTTTGCATTTGTAATTTT
AAAAAATGCTTTCTTCTTTTAATATACTTTTTTGTTTATCTTATTTCTAATACTTTCCCTAATCTCTTTCTTTCAGG
GCAATAATGATACAATGTATCATGCCTCTTTGCACCATTCTAAAGAATAACAGTGATAATTTCTGGGTTAAGGCAAT
AGCAATATTTCTGCATATAAATATTTCTGCATATAAATTGTAACTGATGTAAGAGGTTTCATATTGCTAATAGCAGC
TACAATCCAGCTACCATTCTGCTTTTATTTTATGGTTGGGATAAGGCTGGATTATTCTGAGT**CCAAGCTAGGCCCTT**
**TTGCTAATCATG**TTCATACCTCTTATCTTCCTCCCACAGCTCCTGGGCAACGTGCTGGTCTGTGTGCTGGCCCATCA
CTTTGGCAAAGAATTCACCCCACCAGTGCAGGCTGCCTATCAGAAAGTGGTGGCTGGTGTGGCTAATGCCCTGGCCC
ACAAGTATCAC**TAA**GC**GGCCGC**TCGCTTTCTTGCTGTCCAATTTCTATTAAAGGTTCCTTTGTTCCCTAAGTCCAAC
TACTAAACTGGGGGATATTATGAAGGGCCTCGAGtagggtcatgaaggtttttcttttcctgagaaaacaacacgta
ttgttttctcaggttttgctttttggcctttttctagcttaaaaaaaaaaaaagcaaaagatgctggtggttggcac
tcctggtttccaggacggggttcaaatccctgcggcgtctttgctttgactCTCGAGGCT**GATC**A***GCCTCGACTGTG***
***CCTTCTAG***TTGCCAGCCATCTGTTGTTTGCCCCTCCCCCGTGCCTTCCTTGACCCTGGAAGGTGCCACTCCCACTGT
CCTTTCCTAATAAAATGAGGAAATTGCATCGCATTGTCTGAGTAGGTGTCATTCTATTCTGGGGGGTGGGGTGGGGC
AGGACAGCAAGGGGGAGGATTGGGAAGACAATAGCAGGCATGCTGGGGATGCGGTGGGCTCTATGGCTTCTGAGGCG
GAAAGAACCAGCTGGGGCTCTAGGGGGTATCCCCACGCGCCCTGTAGCGGCGCATTAAGCGCGGCGGGTGTGGTGGT
TACGCGCAGCGTGACCGCTACACTTGCCAGCGCCCTAGCGCCCGCTCCTTTCGCTTTCTTCCCTTCCTTTCTCGCCA
CGTTCGCCGGCTTTCCCCGTCAAGCTCTAAATCGGGGGCTCCCTTTAGGGTTCCGATTTAGTGCTTTACGGCACCTC
GACCCCAAAAAACTTGATTAGGGTGATGGTTCACGTAGGTAC**G**ctagcaccgttggtttccgtagtgtagtggttat
cacgttcgcctaacacgcgaaaggtccccggttcgaaaccgggcactacaaaccaacaacgctGGTACCCCTA*GAAG*
*TTCCTATTCCGAAGTTCCTATTCTCTAGAAAGTATAGGAACTTC*CTTGGCCAAAAAGCCTGAACTCACCGCGACGTC
TGTCGAGAAGTTTCTGATCGAAAAGTTCGACAGCGTCTCCGACCTGATGCAGCTCTCGGAGGGCGAAGAATCTCGTG
CTTTCAGCTTCGATGTAGGAGGGCGTGGATATGTCCTGCGGGTAAATAGCTGCGCCGATGGTTTCTACAAAGATCGT
TATGTTTATCGGCACTTTGCATCGGCCGCGCTCCCGATTCCGGAAGTGCTTGACATTGGGGAATTCAGCGAGAGCCT
GACCTATTGCATCTCCCGCCGTGCACAGGGTGTCACGTTGCAAGACCTGCCTGAAACCGAACTGCCCGCTGTTCTGC
AGCCGGTCGCGGAGGCCATGGATGCGATCGCTGCGGCCGATCTTAGCCAGACGAGCGGGTTCGGCCCATTCGGACCG
CAAGGAATCGGTCAATACACTACATGGCGTGATTTCATATGCGCGATTGCTGATCCCCATGTGTATCACTGGCAAAC
TGTGATGGACGACACCGTCAGTGCGTCCGTCGCGCAGGCTCTCGATGAGCTGATGCTTTGGGCCGAGGACTGCCCCG
AAGTCCGGCACCTCGTGCACGCGGATTTCGGCTCCAACAATGTCCTGACGGACAATGGCCGCATAACAGCGGTCATT
GACTGGAGCGAGGCGATGTTCGGGGATTCCCAATACGAGGTCGCCAACATCTTCTTCTGGAGGCCGTGGTTGGCTTG
TATGGAGCAGCAGACGCGCTACTTCGAGCGGAGGCATCCGGAGCTTGCAGGATCGCCGCGGCTCCGGGCGTATATGC
TCCGCATTGGTCTTGACCAACTCTATCAGAGCTTGGTTGACGGCAATTTCGATGATGCAGCTTGGGCGCAGGGTCGA
TGCGACGCAATCGTCCGATCCGGAGCCGGGACTGTCGGGCGTACACAAATCGCCCGCAGAAGCGCGGCCGTCTGGAC
CGATGGCTGTGTAGAAGTACTCGCCGATAGTGGAAACCGACGCCCCAGCACTCGTCCGAGGGCAAAGGAA**TAG**CACG
TACTACGAGATTTCGATTCCACCGCCGCCTTCTATGAAAGGTTGGGCTTCGGAATCGTTTTCCGGGACGCCGGCTGG
ATGATCCTCCAGCGCGGGGATCTCATGCTGGAGTTCTTCGCCCACCCCAACTTGTTTATTGCAGCTTATAATGGTTA
CAAATAAAGCAATAGCATCACAAATTTCACAAATAAAGCATTTTTTTCACTGCATTCTAGTTGTGGTTTGTCCAAAC
TCATCAATGTATCTTATCATGTCTGTATACCGTCGACaggtggcacttttcggggaaatgtgcgcggaacccctatt
tgtttatttttctaaatacattcaaatatgtatccgctcatgagacaataaccctgataaatgcttcaataatattg
aaaaaggaagagtatgagtattcaacatttccgtgtcgcccttattcccttttttgcggcattttgccttcctgttt
ttgctcacccagaaacgctggtgaaagtaaaagatgctgaagatcagttgggtgcacgagtgggttacatcgaactg
gatctcaacagcggtaagatccttgagagttttcgccccgaagaacgttttccaatgatgagcacttttaaagttct
gctatgtggcgcggtattatcccgtattgacgccgggcaagagcaactcggtcgccgcatacactattctcagaatg
acttggttgagtactcaccagtcacagaaaagcatcttacggatggcatgacagtaagagaattatgcagtgctgcc
ataaccatgagtgataacactgcggccaacttacttctgacaacgatcggaggaccgaaggagctaaccgcttttt
gcacaacatgggggatcatgtaactcgccttgatcgttgggaaccggagctgaatgaagccataccaaacgacgagc
gtgacaccacgatgcctgtagcaatggcaacaacgttgcgcaaactattaactggcgaactacttactctagcttcc
cggcaacaattaatagactggatggaggcggataaagttgcaggaccacttctgcgctcg**gcccttccggc**tggctg
gtttattgctgataaatctggagccggtgagcgtgggtctcgcggtatcattgcagcactggggccagatggtaagc
cctcccgtatcgtagttatctacacgacggggagtcaggcaactatggatgaacgaaatagacagatcgctgagata
ggtgcctcactgattaagcattggtaactgtcagaccaagtttactcatatatactttagattgatttaaa

**B) pAVA3000 [GFP-ENE(C8351G)-mascRNA, Puromycin$^{R}$]**

**tDNA1: 1045-1141**
**EF1alfa promoter: 1176-2679**
**5xEGFP: 2702-6439**
**TEV protease: 6440-7165**
**PEST (degradation sequence): 7193-7327**
**Human beta-globin (HBB): 7334-8718**
**MALAT1-ENE(C8351G)-mascRNA: 8723-8882**
**BGH polyA: 8963-9141**
**tDNA2: 9425-9521**
**FRT: 9532-9558**
**Puromycin$^{R}$ (lacks promoter and the 1$^{st}$ Methionine): 9755-10584**

acttcattttaatttaaaaggatctaggtgaagatcctttttgataatctcatgaccaaaatcccttaacgtgagt
tttcgttccactgagcgtcagaccccgtagaaaagatcaaaggatcttcttgagatcctttttttctgcgcgtaatc
tgctgcttgcaaacaaaaaaaccaccgctaccagcggtggtttgtttgccggatcaagagctaccaactctttttcc
gaaggtaactggcttcagcagagcgcagataccaaatactgtccttctagtgtagccgtagttaggccaccacttca
agaactctgtagcaccgcctacatacctcgctctgctaatcctgttaccagtggctgctgccagtggcgataagtcg
tgtcttaccgggttggactcaagacgatagttaccggataaggcgcagcggtcgggctgaacggggggttcgtgcac
acagcccagcttggagcgaacgacctacaccgaactgagatacctacagcgtgagctatgagaaagcgccacgcttc
ccgaagggagaaaggcggacaggtatccggtaagcggcagggtcggaacaggagagcgcacgagggagcttccaggg
ggaaacgcctggtatctttatagtcctgtcgggtttcgccacctctgacttgagcgtcgatttttgtgatgctcgtc
aggggggcggagcctatggaaaaacgccagcaacgcggcctttttacggttcctggccttttgctggccttttgctc
acatgttctttcctgcgttatcccctgattctgtggataaccgtattaccgcctttgagtgagctgataccgctcgc
cgcagccgaacgaccgagcgcagcgagtcagtgagcgaggaagcggaagagcgcccaatacgcaaaccgcctctccc
cgcgcgttggccgattcattaatgcagctggcacgacaggtttcccgactggaaagcgggcagtgagcgcaacgcaa
ttaatgtgagttagctcactcattaggcaccccaggcACGCGAagcgttgttggtttgtagtgcccggtttcgaacc
ggggacctttcgcgtgttaggcgaacgtgataaccactacactacggaaaccaacggt**gctag**ACGCGTattgtcta
ttctgactcggatcCGTACG**aattctcatgtttgacagcttatcatcgat*tagctt*tggagctaagccagcaatggt**
**agagggaagattctgcacgtcccttccaggcggcctccccgtcaccaccccccccaacccgccccgaccggagctga**
**gagtaattcatacaaaaggactcgcccctgccttggggaatcccagggaccgtcgttaaactcccactaacgtagaa**
**cccagagatcgctgcgttcccgcccctcacccgcccgctctcgtcatcactgaggtggagaagagcatgcgtgagg**
**ctccggtgcccgtcagtgggcagagcgcacatcgcccacagtccccgagaagttggggggaggggtcggcaattgaa**
**ccggtgcctagagaaagtggcgcggggtaaactgggaaagtgatgtcgtgtactggctccgcctttttcccgagggt**
**gggggagaaccgtatataagtgcagtagtcgccgtgaacgttctttttcgcaacgggtttgccgccagaacacaggt**
**aagtgccgtgtgtggttcccgcgggcctggcctctttacgggttatggcccttgcgtgccttgaattacttccacgc**
**ccctggctgcagtacgtgattcttgatcccgagcttcgggttggaagtgggtgggagagttcgaggccttgcgctta**
**aggagccccttcgcctcgtgcttgagttgaggcctggcctgggcgctggggccgccgcgtgcgaatctggtggcacc**
**ttcgcgcctgtctcgctgctttcgataagtctctagccatttaaaatttttgatgacctgctgcgacgctttttttc**
**tggcaagatagtcttgtaaatgcgggccaagatctgcacactggtatttcggtttttggggccgcgggcggcgacgg**
**ggcccgtgcgtcccagcgcacatgttcggcgaggcggggcctgcgagcgcggccaccgagaatcggacgggggtagt**
**ctcaagctggccggcctgctctggtgcctggcctcgcgccgccgtgtatcgccccgccctgggcggcaaggctgg**
**cccggtcggcaccagttgcgtgagcggaaagatggccgcttcccggccctgctgcagggagctcaaaatggaggacg**
**cggcgctcgggagagcgggcgggtgagtcacccacacaaaggaaaagggcctttccgtcctcagccgtcgcttcatg**
**tgactccacggagtaccgggcgccgtccaggcacctcgattagttctcgagcttttggagtacgtcgtctttaggtt**
**ggggggaggggttttatgcgatggagtttccccacactgagtgggtggagactgaagttaggccagcttggcacttg**
**atgtaattctccttggaatttgcccttttttgagtttggatcttggttcattctcaagcctcagacagtggttcaaag**
**tttttttcttccatttcaggtgtcgtgaaaactctagcgtttccgctctagaactagtcca**GCTAGCGCTACCGGTC
GCCACC**atggtgagcaagggcgaggagctgttcaccggggtggtgcccatcctggtcgagctggacggcgacgtaaa**
**cggccacaagttcagcgtgtccggcgagggcgagggcgatgccacctacggcaagctgaccctgaagttcatctgca**
**ccaccggcaagctgcccgtgccctggcccaccctcgtgaccaccctgacctacggcgtgcagtgcttcagccgctac**
**cccgaccacatgaagcagcacgacttcttcaagtccgccatgcccgaaggctacgtccaggagcgcaccatcttctt**
**caaggacgacggcaactacaagacccgcgccgaggtgaagttcgagggcgacaccctggtgaaccgcatcgagctga**
**agggcatcgacttcaaggaggacggcaacatcctggggcacaagctggagtacaactacaacagccacaacgtctat**

atcatggccgacaagcagaagaacggcatcaaggtgaacttcaagatccgccacaacatcgaggacggcagcgtgca
gctcgccgaccactaccagcagaacacccccatcggcgacggccccgtgctgctgcccgacaaccactacctgagca
cccagtccgccctgagcaaagaccccaacgagaagcgcgatcacatggtcctgctggagttcgtgaccgccgccggg
atcactctcggcatggacgagctgtacaagGTcactCcgtgTgaaaacctgtacttccaaggtatggtgagcaaggg
cgaggagctgttcaccggggtggtgcccatcctggtcgagctggacggcgacgtaaacggccacaagttcagcgtgt
ccggcgagggcgagggcgatgccacctacggcaagctgaccctgaagttcatctgcaccaccggcaagctgcccgtg
ccctggcccaccctcgtgaccaccctgacctacggcgtgcagtgcttcagccgctaccccgaccacatgaagcagca
cgacttcttcaagtccgccatgcccgaaggctacgtccaggagcgcaccatcttcttcaaggacgacggcaactaca
agacccgcgccgaggtgaagttcgagggcgacaccctggtgaaccgcatcgagctgaagggcatcgacttcaaggag
gacggcaacatcctggggcacaagctggagtacaactacaacagccacaacgtctatatcatggccgacaagcagaa
gaacggcatcaaggtgaacttcaagatccgccacaacatcgaggacggcagcgtgcagctcgccgaccactaccagc
agaacacccccatcggcgacggccccgtgctgctgcccgacaaccactacctgagcacccagtccgccctgagcaaa
gaccccaacgagaagcgcgatcacatggtcctgctggagttcgtgaccgccgccgggatcactctcggcatggacga
gctgtacaagACcactGcgtgCgaaaacctgtacttccaaggtatggtgagcaagggcgaggagctgttcaccgggg
tggtgcccatcctggtcgagctggacggcgacgtaaacggccacaagttcagcgtgtccggcgagggcgagggcgat
gccacctacggcaagctgaccctgaagttcatctgcaccaccggcaagctgcccgtgccctggcccaccctcgtgac
caccctgacctacggcgtgcagtgcttcagccgctaccccgaccacatgaagcagcacgacttcttcaagtccgcca
tgcccgaaggctacgtccaggagcgcaccatcttcttcaaggacgacggcaactacaagacccgcgccgaggtgaag
ttcgagggcgacaccctggtgaaccgcatcgagctgaagggcatcgacttcaaggaggacggcaacatcctggggca
caagctggagtacaactacaacagccacaacgtctatatcatggccgacaagcagaagaacggcatcaaggtgaact
tcaagatccgccacaacatcgaggacggcagcgtgcagctcgccgaccactaccagcagaacacccccatcggcgac
ggccccgtgctgctgcccgacaaccactacctgagcacccagtccgccctgagcaaagaccccaacgagaagcgcga
tcacatggtcctgctggagttcgtgaccgccgccgggatcactctcggcatggacgagctgtacaagGTcactAcgt
gTgaaaacctgtacttccaaggtatggtgagcaagggcgaggagctgttcaccggggtggtgcccatcctggtcgag
ctggacggcgacgtaaacggccacaagttcagcgtgtccggcgagggcgagggcgatgccacctacggcaagctgac
cctgaagttcatctgcaccaccggcaagctgcccgtgccctggcccaccctcgtgaccaccctgacctacggcgtgc
agtgcttcagccgctaccccgaccacatgaagcagcacgacttcttcaagtccgccatgcccgaaggctacgtccag
gagcgcaccatcttcttcaaggacgacggcaactacaagacccgcgccgaggtgaagttcgagggcgacaccctggt
gaaccgcatcgagctgaagggcatcgacttcaaggaggacggcaacatcctggggcacaagctggagtacaactaca
acagccacaacgtctatatcatggccgacaagcagaagaacggcatcaaggtgaacttcaagatccgccacaacatc
gaggacggcagcgtgcagctcgccgaccactaccagcagaacacccccatcggcgacggccccgtgctgctgcccga
caaccactacctgagcacccagtccgccctgagcaaagaccccaacgagaagcgcgatcacatggtcctgctggagt
tcgtgaccgccgccgggatcactctcggcatggacgagctgtacaagACcactTcgtgCgaaaacctgtacttccaa
ggtatggtgagcaagggcgaggagctgttcaccggggtggtgcccatcctggtcgagctggacggcgacgtaaacgg
ccacaagttcagcgtgtccggcgagggcgagggcgatgccacctacggcaagctgaccctgaagttcatctgcacca
ccggcaagctgcccgtgccctggcccaccctcgtgaccaccctgacctacggcgtgcagtgcttcagccgctacccc
gaccacatgaagcagcacgacttcttcaagtccgccatgcccgaaggctacgtccaggagcgcaccatcttcttcaa
ggacgacggcaactacaagacccgcgccgaggtgaagttcgagggcgacaccctggtgaaccgcatcgagctgaagg
gcatcgacttcaaggaggacggcaacatcctggggcacaagctggagtacaactacaacagccacaacgtctatatc
atggccgacaagcagaagaacggcatcaaggtgaacttcaagatccgccacaacatcgaggacggcagcgtgcagct
cgccgaccactaccagcagaacacccccatcggcgacggccccgtgctgctgcccgacaaccactacctgagcaccc
agtccgccctgagcaaagaccccaacgagaagcgcgatcacatggtcctgctggagttcgtgaccgccgccgggatc
actctcggcatggacgagctgtacaaggaaaacctgtacttccaaAgtggagaAAGCTTgtttaaggaccacgtga
ttacaacccgatatcgagcaccatttgtcatttgacgaatgaatctgatgggcacacaacatcgttgtatggtattg
gatttggtcccttcatcattacaaacaagcacttgtttagaagaaataatggaacactgttggtccaatcactacat
ggtgtattcaaggtcaagaacaccacgactttgcaacaacacctcattgatgggagggacatgataattattcgcat
gcctaaggatttcccaccatttcctcaaaagctgaaatttagagagccacaaagggaagagcgcatatgtcttgtga
caaccaacttccaaactaagagcatgtctagcatggtgtcagacactagttgcacattcccttcatctgatggcata
ttctggaagcattggattcaaaccaaggatgggcagtgtggcagtccattagtatcaactagagatgggttcattgt
tggtatacactcagcatcgaatttcaccaacacaaacaattatttcacaagcgtgccgaaaaacttcatggaattgt
tgacaaatcaggaggcgcagcagtgggttagtggttggcgattaaatgctgactcagtattgtgggggggccataaa
gttttcatgagcaaacctgaagagccttttcagccagttaaggaagcgactcaactcatgaatgaattggtgtactc
gcaagaaaacctgtacttccaaagtATGCATAGATCACGAGATATCAGCCATGGCTTCCCGCCGGCGGTGGCGGCGC
AGGATGATGGCACGCTGCCCATGTCTTGTGCCCAGGAGAGCGGGATGGACCGTCACCCTGCAGCCTGTGCTTCTGCT
AGGATCAATGTGGAGCTCGTGCACCTGACTCCTGAGGAGAAGTCTGCCGTTACTGCCCTGTGGGGCAAGGTGAACGT
GGATGAAGTTGGTGGTGAGGCCCTGGGCAGGCTGCTGGTGGTCTACCCTTGGACCCAGAGGTTCTTTGAGTCCTTTG
GGGATCTGTCCACTCCTGATGCTGTTATGGGCAACCCTAAGGTGAAGGCTCATGGCAAGAAAGTGCTCGGTGCCTTT

AGTGATGGCCTGGCTCACCTGGACAACCTCAAGGGCACCTTTGCCACACTGAGTGAGCTGCACTGTGACAAGCTGCA
CGTGGATCCTGAGAACTTCAGGGTGAGTCTATGGGACCCTTGATGTTTTCTTTCCCCTTCTTTTCTATGGTTAAGTT
CATGTCATAGGAAGGGGATAAGTAACAGGGTACAGTTTAGAATGGGAAACAGACGAATGATTGCATCAGTGTGGAAG
TCTCAGGATCGTTTTAGTTTCTTTTATTTGCTGTTCATAACAATTGTTTTCTTTTGTTTAATTCTTGCTTTCTTTTT
TTTTCTTCTCCGCAATTTTTACTATTATACTTAATGCCTTAACATTGTGTATAACAAAAGGAAATATCTCTGAGATA
CATTAAGTAACTTAAAAAAAAAACTTTACACAGTCTGCCTAGTACATTACTATTTGGAATATATGTGTGCTTATTTG
CATATTCATAATCTCCCTACTTTATTTTCTTTTATTTTTAATTGATACATAATCATTATACATATTTATGGGTTAAA
GTGTAATGTTTTAATATGTGTACACATATTGACCAAATCAGGGTAATTTTGCATTTGTAATTTTAAAAAATGCTTTC
TTCTTTTAATATACTTTTTTGTTTATCTTATTTCTAATACTTTCCCTAATCTCTTTCTTTCAGGGCAATAATGATAC
AATGTATCATGCCTCTTTGCACCATTCTAAAGAATAACAGTGATAATTTCTGGGTTAAGGCAATAGCAATATTTCTG
CATATAAATATTTCTGCATATAAATTGTAACTGATGTAAGAGGTTTCATATTGCTAATAGCAGCTACAATCCAGCTA
CCATTCTGCTTTTATTTTATGGTTGGGATAAGGCTGGATTATTCTGAGT**CCAAGCTAGGCCCTTTTGCTAATCATG**T
TCATACCTCTTATCTTCCTCCCACAGCTCCTGGGCAACGTGCTGGTCTGTGTGCTGGCCCATCACTTTGGCAAAGAA
TTCACCCCACCAGTGCAGGCTGCCTATCAGAAAGTGGTGGCTGGTGTGGCTAATGCCCTGGCCCACAAGTATCAC**TA**
**A**GC**GGCCGC**TCGCTTTCTTGCTGTCCAATTTCTATTAAAGGTTCCTTTGTTCCCTAAGTCCAACTACTAAACTGGGG
GATATTATGAAGGGCCTCGAGtagggtcatgaaggtttttcttttcctgagaaaacaacacgtattgttttctcagg
ttttgcttttggcctttttctagcttaaaaaaaaaaaaaggaaaagatgctggtggttggcactcctggtttccag
gacggggttcaaatccctgcggcgtctttgctttgactCTCGAGGCT**GATC**A***GCCTCGACTGTGCCTTCTAG***TTGCC
AGCCATCTGTTGTTTGCCCCTCCCCCGTGCCTTCCTTGACCCTGGAAGGTGCCACTCCCACTGTCCTTTCCTAATAA
AATGAGGAAATTGCATCGCATTGTCTGAGTAGGTGTCATTCTATTCTGGGGGGTGGGGTGGGGCAGGACAGCAAGGG
GGAGGATTGGGAAGACAATAGCAGGCATGCTGGGGATGCGGTGGGCTCTATGGCTTCTGAGGCGGAAAGAACCAGCT
GGGGCTCTAGGGGGTATCCCCACGCGCCCTGTAGCGGCGCATTAAGCGCGGCGGGTGTGGTGGTTACGCGCAGCGTG
ACCGCTACACTTGCCAGCGCCCTAGCGCCCGCTCCTTTCGCTTTCTTCCCTTCCTTTCTCGCCACGTTCGCCGGCTT
TCCCCGTCAAGCTCTAAATCGGGGGCTCCCTTTAGGGTTCCGATTTAGTGCTTTACGGCACCTCGACCCCAAAAAAC
TTGATTAGGGTGATGGTTCACGTAGGTAC**G**ctagcaccgttggtttccgtagtgtagtggttatcacgttcgcctaa
cacgcgaaaggtccccggttcgaaaccgggcactacaaaccaacaacgctGGTACCCCTA*GAAGTTCCTATTCCGAA*
*GTTCCTATTC**TCTAGAAAGTATAGGAACTTC***CTTGGCCAAAAAGCCTGAAagatcacgagatatcagccatggcttc
ccgccggcggtggcggcgcaggatgatggcacgctgcccaCgtcttgtgcccaggagagcgggaCggaccgtcaccc
tgcagcctgtgcttctgctaggatcaatgtggaaaacctgtacttccaaGgtaccgagtacaagcccacggtgcgcc
tcgccacccgcgacgacgtccccagggccgtCcgcaccctcgccgccgcgttcgccgactaccccgccacgcgccac
accgtcgatccggaccgccacatcgagcg*ggtcaccgagctgcaagaactcttcctcacgcgcgtcgggctcgacat*
*cggcaaggtgtgggtcgcggacgacggcgccgcggtggcggtctggaccacgccggagagcgtcgaagcgggggcgg*
*tgttcgccgagatcggcccgcgcatggccgagttgagcggttcccggctggccgcgcagcaacagatggaaggcctc*
*ctggcgccgcaccggcccaaggagcccgcgtggttcctggccaccgtcggcgtctcgcccgaccaccagggcaaggg*
*tctgggcagcgccgtcgtgctccccggagtggaggcggccgagcgcgccggggtgcccgccttcctggagacctccg*
*cgccccgcaacctccccttctacg*agcggctcggcttcaccgtcaccgccgacgtcgaggtgcccgaaggaccgcg**c**
**acctggtg**catgacccgcaagcccggtgcc**tga**ggtaccgctcgctgatcagcctcgactgtgccttctagttgcca
gccatctgttgtttgcccctcccccgtgccttccttgaccctggaaggtgccactcccactgtcctttcctaataaa
atgaggaaattgcatcgcattgtctgagtaggtgtcattctattctggggggtggggtggggcaggacagcaagggg
gaggattgggaagacaatagcaggcatgctggggaGTCGACaggtggcacttttcggggaaatgtgcgcggaacccc
tatttgtttatttttctaaatacattcaaatatgtatccgctcatgagacaataaccctgataaatgcttcaataat
attgaaaaaggaagagtatgagtattcaacatttccgtgtcgcccttattcccttttttgcggcattttgccttcct
gtttttgctcacccagaaacgctggtgaaagtaaaagatgctgaagatcagttgggtgcacgagtgggttacatcga
actggatctcaacagcggtaagatccttgagagttttcgccccgaagaacgttttccaatgatgagcacttttaaag
ttctgctatgtggcgcggtattatcccgtattgacgccgggcaagagcaactcggtcgccgcatacactattctcag
aatgacttggttgagtactcaccagtcacagaaaagcatcttacggatggcatgacagtaagagaattatgcagtgc
tgccataaccatgagtgataacactgcggccaacttacttctgacaacgatcggaggaccgaaggagctaaccgctt
ttttgcacaacatgggggatcatgtaactcgccttgatcgttgggaaccggagctgaatgaagccataccaaacgac
gagcgtgacaccacgatgcctgtagcaatggcaacaacgttgcgcaaactattaactggcgaactacttactctagc
ttcccggcaacaattaatagactggatggaggcggataaagttgcaggaccacttctgcgctcg**gcccttccggc**tg
gctggtttattgctgataaatctggagccggtgagcgtgggtctcgcggtatcattgcagcactggggccagatggt
aagccctcccgtatcgtagttatctacacgacggggagtcaggcaactatggatgaacgaaatagacagatcgctga
gataggtgcctcactgattaagcattggtaactgtcagaccaagtttactcatatatactttagattgatttaaa

**C) pAVA2995 [RFP-ENE(C8351G)-mascRNA, Hygromycin$^R$]**

**tDNA1: 1045-1141**
**EF1alfa promoter: 1176-2679**
**5xtdTomato: 2702-9994**
**TEV protease: 9995-10720**
**PEST (degradation sequence): 10748-10882**
**Human beta-globin(ΔI1): 10889-12271**
**MALAT1-ENE(C8351G)-mascRNA: 12278-12437**
**BGH polyA: 12470-12694**
**tDNA2: 12980-13076**
**FRT: 13087-13113**
**C8351G: 12357**
**Hygromycin$^R$ (lacks promoter and the 1$^{st}$ Methionine): 13154-14430**

acttcattttaatttaaaaggatctaggtgaagatcctttttgataatctcatgaccaaaatcccttaacgtgagt
tttcgttccactgagcgtcagaccccgtagaaaagatcaaaggatcttcttgagatcctttttttctgcgcgtaatc
tgctgcttgcaaacaaaaaaaccaccgctaccagcggtggtttgtttgccggatcaagagctaccaactctttttcc
gaaggtaactggcttcagcagagcgcagataccaaatactgtccttctagtgtagccgtagttaggccaccacttca
agaactctgtagcaccgcctacatacctcgctctgctaatcctgttaccagtggctgctgccagtggcgataagtcg
tgtcttaccgggttggactcaagacgatagttaccggataaggcgcagcggtcgggctgaacggggggttcgtgcac
acagcccagcttggagcgaacgacctacaccgaactgagatacctacagcgtgagctatgagaaagcgccacgcttc
ccgaagggagaaaggcggacaggtatccggtaagcggcagggtcggaacaggagagcgcacgagggagcttccaggg
ggaaacgcctggtatctttatagtcctgtcgggtttcgccacctctgacttgagcgtcgatttttgtgatgctcgtc
aggggggcggagcctatggaaaaacgccagcaacgcggcctttttacggttcctggccttttgctggccttttgctc
acatgttctttcctgcgttatcccctgattctgtggataaccgtattaccgcctttgagtgagctgataccgctcgc
cgcagccgaacgaccgagcgcagcgagtcagtgagcgaggaagcggaagagcgcccaatacgcaaaccgcctctccc
cgcgcgttggccgattcattaatgcagctggcacgacaggtttcccgactggaaagcgggcagtgagcgcaacgcaa
ttaatgtgagttagctcactcattaggcaccccaggcACGCGAagcgttgttggtttgtagtgcccggtttcgaacc
ggggacctttcgcgtgttaggcgaacgtgataaccactacactacggaaaccaacggtgctagACGCGTattgtcta
ttctgactcggatcCGTACGaattctcatgtttgacagcttatcatcgattagctttggagctaagccagcaatggt
agagggaagattctgcacgtcccttccaggcggcctccccgtcaccaccccccccaacccgccccgaccggagctga
gagtaattcatacaaaaggactcgcccctgccttggggaatcccagggaccgtcgttaaactcccactaacgtagaa
cccagagatcgctgcgttcccgcccctcacccgcccgctctcgtcatcactgaggtggagaagagcatgcgtgagg
ctccggtgcccgtcagtgggcagagcgcacatcgcccacagtccccgagaagttggggggaggggtcggcaattgaa
ccggtgcctagagaaagtggcgcggggtaaactgggaaagtgatgtcgtgtactggctccgcctttttcccgagggt
gggggagaaccgtatataagtgcagtagtcgccgtgaacgttctttttcgcaacgggtttgccgccagaacacaggt
aagtgccgtgtgtggttcccgcgggcctggcctctttacgggttatggcccttgcgtgccttgaattacttccacgc
ccctggctgcagtacgtgattcttgatcccgagcttcgggttggaagtgggtgggagagttcgaggccttgcgctta
aggagccccttcgcctcgtgcttgagttgaggcctggcctgggcgctggggccgccgcgtgcgaatctggtggcacc
ttcgcgcctgtctcgctgctttcgataagtctctagccatttaaaatttttgatgacctgctgcgacgctttttttc
tggcaagatagtcttgtaaatgcgggccaagatctgcacactggtatttcggtttttggggccgcgggcggcgacgg
ggcccgtgcgtcccagcgcacatgttcggcgaggcggggcctgcgagcgcggccaccgagaatcggacgggggtagt
ctcaagctggccggcctgctctggtgcctggcctcgcgccgccgtgtatcgccccgccctgggcggcaaggctgg
cccggtcggcaccagttgcgtgagcggaaagatggccgcttcccggccctgctgcagggagctcaaaatggaggacg
cggcgctcgggagagcgggcgggtgagtcacccacacaaaggaaaagggcctttccgtcctcagccgtcgcttcatg
tgactccacggagtaccgggcgccgtccaggcacctcgattagttctcgagcttttggagtacgtcgtctttaggtt
ggggggaggggttttatgcgatggagtttccccacactgagtgggtggagactgaagttaggccagcttggcacttg
atgtaattctccttggaatttgccctttttgagtttggatcttggttcattctcaagcctcagacagtggttcaaag
tttttttcttccatttcaggtgtcgtgaaaactctagcgtttccgctctagaactagtccaGCTAGCGCTACCGGTC
GCCACCatggtgagcaagggcgaggaggtcatcaaagagttcatgcgcttcaaggtgcgcatggagggctccatgaa
cggccacgagttcgagatcgagggcgagggcgagggccgcccctacgagggcacccagaccgccaagctgaaggtga
ccaagggcggccccctgcccttcgcctgggacatcctgtcccccagttcatgtacggctccaaggcgtacgtgaag
caccccgccgacatccccgattacaagaagctgtccttccccgagggcttcaagtgggagcgcgtgatgaacttcga
ggacggcggtctggtgaccgtgacccaggactcctccctgcaggacggcacgctgatctacaaggtgaagatgcgcg

gcaccaacttcccccccgacggccccgtaatgcagaagaagaccatgggctgggaggcctccaccgagcgcctgtac
ccccgcgacggcgtgctgaagggcgagatccaccaggccctgaagctgaaggacggcggccactacctggtggagtt
caagaccatctacatggccaagaagcccgtgcaactgcccggctactactacgtggacaccaagctggacatcacct
cccacaacgaggactacaccatcgtggaacagtacgagcgctccgagggccgccaccacctgttcctggggcatggc
accggcagcaccggcagcggcagctccggcaccgcctcctccgaggacaacaacatggccgtcatcaaagagttcat
gcgcttcaaggtgcgcatggagggctccatgaacggccacgagttcgagatcgagggcgagggcgagggccgcccct
acgagggcacccagaccgccaagctgaaggtgaccaagggcggccccctgcccttcgcctgggacatcctgtccccc
cagttcatgtacggctccaaggcgtacgtgaagcaccccgccgacatccccgattacaagaagctgtccttccccga
gggcttcaagtgggagcgcgtgatgaacttcgaggacggcggtctggtgaccgtgacccaggactcctccctgcagg
acggcacgctgatctacaaggtgaagatgcgcggcaccaacttcccccccgacggccccgtaatgcagaagaagacc
atgggctgggaggcctccaccgagcgcctgtaccccgcgacggcgtgctgaagggcgagatccaccaggccctgaa
gctgaaggacggcggccactacctggtggagttcaagaccatctacatggccaagaagcccgtgcaactgcccggct
actactacgtggacaccaagctggacatcacctcccacaacgaggactacaccatcgtggaacagtacgagcgctcc
gagggccgccaccacctgttcctgtacggcatggacgagctgtacaag**GTcactCcgtgTg**aaaacctgtacttcca
aggt**atg**gtgagcaagggcgaggaggtcatcaaagagttcatgcgcttcaaggtgcgcatggagggctccatgaacg
gccacgagttcgagatcgagggcgagggcgagggccgcccctacgagggcacccagaccgccaagctgaaggtgacc
aagggcggccccctgcccttcgcctgggacatcctgtcccccagttcatgtacggctccaaggcgtacgtgaagca
ccccgccgacatccccgattacaagaagctgtccttccccgagggcttcaagtgggagcgcgtgatgaacttcgagg
acggcggtctggtgaccgtgacccaggactcctccctgcaggacggcacgctgatctacaaggtgaagatgcgcggc
accaacttcccccccgacggccccgtaatgcagaagaagaccatgggctgggaggcctccaccgagcgcctgtaccc
ccgcgacggcgtgctgaagggcgagatccaccaggccctgaagctgaaggacggcggccactacctggtggagttca
agaccatctacatggccaagaagcccgtgcaactgcccggctactactacgtggacaccaagctggacatcacctcc
cacaacgaggactacaccatcgtggaacagtacgagcgctccgagggccgccaccacctgttcctggggcatggcac
cggcagcaccggcagcggcagctccggcaccgcctcctccgaggacaacaacatggccgtcatcaaagagttcatgc
gcttcaaggtgcgcatggagggctccatgaacggccacgagttcgagatcgagggcgagggcgagggccgcccctac
gagggcacccagaccgccaagctgaaggtgaccaagggcggccccctgcccttcgcctgggacatcctgtcccccca
gttcatgtacggctccaaggcgtacgtgaagcaccccgccgacatccccgattacaagaagctgtccttccccgagg
gcttcaagtgggagcgcgtgatgaacttcgaggacggcggtctggtgaccgtgacccaggactcctccctgcaggac
ggcacgctgatctacaaggtgaagatgcgcggcaccaacttcccccccgacggccccgtaatgcagaagaagaccat
gggctgggaggcctccaccgagcgcctgtaccccgcgacggcgtgctgaagggcgagatccaccaggccctgaagc
tgaaggacggcggccactacctggtggagttcaagaccatctacatggccaagaagcccgtgcaactgcccggctac
tactacgtggacaccaagctggacatcacctcccacaacgaggactacaccatcgtggaacagtacgagcgctccga
gggccgccaccacctgttcctgtacggcatggacgagctgtacaag**ACcactGcgtgC**gaaaacctgtacttccaag
gt**atg**gtgagcaagggcgaggaggtcatcaaagagttcatgcgcttcaaggtgcgcatggagggctccatgaacggc
cacgagttcgagatcgagggcgagggcgagggccgcccctacgagggcacccagaccgccaagctgaaggtgaccaa
gggcggccccctgcccttcgcctgggacatcctgtcccccagttcatgtacggctccaaggcgtacgtgaagcacc
ccgccgacatccccgattacaagaagctgtccttccccgagggcttcaagtgggagcgcgtgatgaacttcgaggac
ggcggtctggtgaccgtgacccaggactcctccctgcaggacggcacgctgatctacaaggtgaagatgcgcggcac
caacttcccccccgacggccccgtaatgcagaagaagaccatgggctgggaggcctccaccgagcgcctgtaccccc
gcgacggcgtgctgaagggcgagatccaccaggccctgaagctgaaggacggcggccactacctggtggagttcaag
accatctacatggccaagaagcccgtgcaactgcccggctactactacgtggacaccaagctggacatcacctccca
caacgaggactacaccatcgtggaacagtacgagcgctccgagggccgccaccacctgttcctggggcatggcaccg
gcagcaccggcagcggcagctccggcaccgcctcctccgaggacaacaacatggccgtcatcaaagagttcatgcgc
ttcaaggtgcgcatggagggctccatgaacggccacgagttcgagatcgagggcgagggcgagggccgcccctacga
gggcacccagaccgccaagctgaaggtgaccaagggcggccccctgcccttcgcctgggacatcctgtcccccccagt
tcatgtacggctccaaggcgtacgtgaagcaccccgccgacatccccgattacaagaagctgtccttccccgagggc
ttcaagtgggagcgcgtgatgaacttcgaggacggcggtctggtgaccgtgacccaggactcctccctgcaggacgg
cacgctgatctacaaggtgaagatgcgcggcaccaacttcccccccgacggccccgtaatgcagaagaagaccatgg
gctgggaggcctccaccgagcgcctgtaccccgcgacggcgtgctgaagggcgagatccaccaggccctgaagctg
aaggacggcggccactacctggtggagttcaagaccatctacatggccaagaagcccgtgcaactgcccggctacta
ctacgtggacaccaagctggacatcacctcccacaacgaggactacaccatcgtggaacagtacgagcgctccgagg
gccgccaccacctgttcctgtacggcatggacgagctgtacaag**GTcactAcgtgTg**aaaacctgtacttccaaggt
**atg**gtgagcaagggcgaggaggtcatcaaagagttcatgcgcttcaaggtgcgcatggagggctccatgaacggcca
cgagttcgagatcgagggcgagggcgagggccgcccctacgagggcacccagaccgccaagctgaaggtgaccaagg
gcggccccctgcccttcgcctgggacatcctgtcccccagttcatgtacggctccaaggcgtacgtgaagcacccc
gccgacatccccgattacaagaagctgtccttccccgagggcttcaagtgggagcgcgtgatgaacttcgaggacgg
cggtctggtgaccgtgacccaggactcctccctgcaggacggcacgctgatctacaaggtgaagatgcgcggcacca

acttccccccgacggccccgtaatgcagaagaagaccatgggctgggaggcctccaccgagcgcctgtacccccgc
gacggcgtgctgaagggcgagatccaccaggccctgaagctgaaggacggcggccactacctggtggagttcaagac
catctacatggccaagaagcccgtgcaactgcccggctactactacgtggacaccaagctggacatcacctcccaca
acgaggactacaccatcgtggaacagtacgagcgctccgagggccgccaccacctgttcctggggcatggcaccggc
agcaccggcagcggcagctccggcaccgcctcctccgaggacaacaacatggccgtcatcaaagagttcatgcgctt
caaggtgcgcatggagggctccatgaacggccacgagttcgagatcgagggcgagggcgagggccgcccctacgagg
gcacccagaccgccaagctgaaggtgaccaagggcggcccccctgcccttcgcctgggacatcctgtcccccccagttc
atgtacggctccaaggcgtacgtgaagcaccccgccgacatccccgattacaagaagctgtccttccccgagggctt
caagtgggagcgcgtgatgaacttcgaggacggcggtctggtgaccgtgacccaggactcctccctgcaggacggca
cgctgatctacaaggtgaagatgcgcggcaccaacttccccccccgacggccccgtaatgcagaagaagaccatgggc
tgggaggcctccaccgagcgcctgtacccccgcgacggcgtgctgaagggcgagatccaccaggccctgaagctgaa
ggacggcggccactacctggtggagttcaagaccatctacatggccaagaagcccgtgcaactgcccggctactact
acgtggacaccaagctggacatcacctcccacaacgaggactacaccatcgtggaacagtacgagcgctccgagggc
cgccaccacctgttcctgtacggcatggacgagctgtacaagACcacTTcgtgCgaaaacctgtacttccaaggtat
ggtgagcaagggcgaggaggtcatcaaagagttcatgcgcttcaaggtgcgcatggagggctccatgaacggccacg
agttcgagatcgagggcgagggcgagggccgcccctacgagggcacccagaccgccaagctgaaggtgaccaagggc
ggcccccctgcccttcgcctgggacatcctgtcccccccagttcatgtacggctccaaggcgtacgtgaagcaccccgc
cgacatccccgattacaagaagctgtccttccccgagggcttcaagtgggagcgcgtgatgaacttcgaggacggcg
gtctggtgaccgtgacccaggactcctccctgcaggacggcacgctgatctacaaggtgaagatgcgcggcaccaac
ttccccccccgacggccccgtaatgcagaagaagaccatgggctgggaggcctccaccgagcgcctgtacccccgcga
cggcgtgctgaagggcgagatccaccaggccctgaagctgaaggacggcggccactacctggtggagttcaagacca
tctacatggccaagaagcccgtgcaactgcccggctactactacgtggacaccaagctggacatcacctcccacaac
gaggactacaccatcgtggaacagtacgagcgctccgagggccgccaccacctgttcctggggcatggcaccggcag
caccggcagcggcagctccggcaccgcctcctccgaggacaacaacatggccgtcatcaaagagttcatgcgcttca
aggtgcgcatggagggctccatgaacggccacgagttcgagatcgagggcgagggcgagggccgcccctacgagggc
acccagaccgccaagctgaaggtgaccaagggcggcccccctgcccttcgcctgggacatcctgtcccccccagttcat
gtacggctccaaggcgtacgtgaagcaccccgccgacatccccgattacaagaagctgtccttccccgagggcttca
agtgggagcgcgtgatgaacttcgaggacggcggtctggtgaccgtgacccaggactcctccctgcaggacggcacg
ctgatctacaaggtgaagatgcgcggcaccaacttccccccccgacggccccgtaatgcagaagaagaccatgggctg
ggaggcctccaccgagcgcctgtacccccgcgacggcgtgctgaagggcgagatccaccaggccctgaagctgaagg
acggcggccactacctggtggagttcaagaccatctacatggccaagaagcccgtgcaactgcccggctactactac
gtggacaccaagctggacatcacctcccacaacgaggactacaccatcgtggaacagtacgagcgctccgagggccg
ccaccacctgttcctgtacggcatggacgagctgtacaaggaaaacctgtacttccaaAgtggagaAAGCTTgttta
agggaccacgtgattacaacccgatatcgagcaccatttgtcatttgacgaatgaatctgatgggcacacaacatcg
ttgtatggtattggatttggtcccttcatcattacaaacaagcacttgtttagaagaaataatggaacactgttggt
ccaatcactacatggtgtattcaaggtcaagaacaccacgactttgcaacaacacctcattgatgggagggacatga
taattattcgcatgcctaaggatttcccaccatttcctcaaaagctgaaatttagagagccacaaagggaagagcgc
atatgtcttgtgacaaccaacttccaaactaagagcatgtctagcatggtgtcagacactagttgcacattcccttc
atctgatggcatattctggaagcattggattcaaaccaaggatgggcagtgtggcagtccattagtatcaactagag
atgggttcattgttggtatacactcagcatcgaatttcaccaacacaaacaattatttcacaagcgtgccgaaaaac
ttcatggaattgttgacaaatcaggaggcgcagcagtgggttagtggttggcgattaaatgctgactcagtattgtg
gggggccataaagttttcatgagcaaacctgaagagccttttcagccagttaaggaagcgactcaactcatgaatg
aattggtgtactcgcaagaaaacctgtacttccaaagtATGCATAGATCACGAGATATCAGCCATGGCTTCCCGCCG
GCGGTGGCGGCGCAGGATGATGGCACGCTGCCCATGTCTTGTGCCCAGGAGAGCGGGATGGACCGTCACCCTGCAGC
CTGTGCTTCTGCTAGGATCAATGTGGAGCTCGTGCACCTGACTCCTGAGGAGAAGTCTGCCGTTACTGCCCTGTGGG
GCAAGGTGAACGTGGATGAAGTTGGTGGTGAGGCCCTGGGCAGGCTGCTGGTGGTCTACCCTTGGACCCAGAGGTTC
TTTGAGTCCTTTGGGGATCTGTCCACTCCTGATGCTGTTATGGGCAACCCTAAGGTGAAGGCTCATGGCAAGAAAGT
GCTCGGTGCCTTTAGTGATGGCCTGGCTCACCTGGACAACCTCAAGGGCACCTTTGCCACACTGAGTGAGCTGCACT
GTGACAAGCTGCACGTGGATCCTGAGAACTTCAGGGTGAGTCTATGGGACCCTTGATGTTTTCTTTCCCCTTCTTTT
CTATGGTTAAGTTCATGTCATAGGAAGGGGATAAGTAACAGGGTACAGTTTAGAATGGGAAACAGACGAATGATTGC
ATCAGTGTGGAAGTCTCAGGATCGTTTTAGTTTCTTTTATTTGCTGTTCATAACAATTGTTTTCTTTTGTTTAATTC
TTGCTTTCTTTTTTTTTCTTCTCCGCAATTTTTACTATTATACTTAATGCCTTAACATTGTGTATAACAAAAGGAAA
TATCTCTGAGATACATTAAGTAACTTAAAAAAAAACTTTACACAGTCTGCCTAGTACATTACTATTTGGAATATAT
GTGTGCTTATTTGCATATTCATAATCTCCCTACTTTATTTTCTTTTATTTTTAATTGATACATAATCATTATACATA
TTTATGGGTTAAAGTGTAATGTTTTAATATGTGTACACATATTGACCAAATCAGGGTAATTTTGCATTTGTAATTTT
AAAAAATGCTTTCTTCTTTTAATATACTTTTTTGTTTATCTTATTTCTAATACTTTCCCTAATCTCTTTCTTTCAGG
GCAATAATGATACAATGTATCATGCCTCTTTGCACCATTCTAAAGAATAACAGTGATAATTTCTGGGTTAAGGCAAT

AGCAATATTTCTGCATATAAATATTTCTGCATATAAATTGTAACTGATGTAAGAGGTTTCATATTGCTAATAGCAGC
TACAATCCAGCTACCATTCTGCTTTTATTTTATGGTTGGGATAAGGCTGGATTATTCTGAGT**CCAAGCTAGGCCCTT**
**TTGCTAATCATG**TTCATACCTCTTATCTTCCTCCCACAGCTCCTGGGCAACGTGCTGGTCTGTGTGCTGGCCCATCA
CTTTGGCAAAGAATTCACCCCACCAGTGCAGGCTGCCTATCAGAAAGTGGTGGCTGGTGTGGCTAATGCCCTGGCCC
ACAAGTATCAC**TAA**GC**GGCCGC**TCGCTTTCTTGCTGTCCAATTTCTATTAAAGGTTCCTTTGTTCCCTAAGTCCAAC
TACTAAACTGGGGGATATTATGAAGGGCCTCGAGtagggtcatgaaggtttttcttttcctgagaaaacaacacgta
ttgttttctcaggttttgctttttggcctttttctagcttaaaaaaaaaaaaaggaaaagatgctggtggttggcac
tcctggtttccaggacggggttcaaatccctgcggcgtctttgctttgactCTCGAGGCT**GATC**A***GCCTCGACTGTG***
***CCTTCTAG***TTGCCAGCCATCTGTTGTTTGCCCCTCCCCCGTGCCTTCCTTGACCCTGGAAGGTGCCACTCCCACTGT
CCTTTCCTAATAAAATGAGGAAATTGCATCGCATTGTCTGAGTAGGTGTCATTCTATTCTGGGGGGTGGGGTGGGGC
AGGACAGCAAGGGGGAGGATTGGGAAGACAATAGCAGGCATGCTGGGGATGCGGTGGGCTCTATGGCTTCTGAGGCG
GAAAGAACCAGCTGGGGCTCTAGGGGGTATCCCCACGCGCCCTGTAGCGGCGCATTAAGCGCGGCGGGTGTGGTGGT
TACGCGCAGCGTGACCGCTACACTTGCCAGCGCCCTAGCGCCCGCTCCTTTCGCTTTCTTCCCTTCCTTTCTCGCCA
CGTTCGCCGGCTTTCCCCGTCAAGCTCTAAATCGGGGGCTCCCTTTAGGGTTCCGATTTAGTGCTTTACGGCACCTC
GACCCCAAAAAACTTGATTAGGGTGATGGTTCACGTAGGTAC**G**ctagcaccgttggtttccgtagtgtagtggttat
cacgttcgcctaacacgcgaaaggtccccggttcgaaaccgggcactacaaaccaacaacgctGGTACCCCTA*GAAG*
*TTCCTATTCCGAAGTTCCTATTCTCTAGAAAGTATAGGAACTTC*CTTGGCCAAAAAGCCTGAACTCACCGCGACGTC
TGTCGAGAAGTTTCTGATCGAAAAGTTCGACAGCGTCTCCGACCTGATGCAGCTCTCGGAGGGCGAAGAATCTCGTG
CTTTCAGCTTCGATGTAGGAGGGCGTGGATATGTCCTGCGGGTAAATAGCTGCGCCGATGGTTTCTACAAAGATCGT
TATGTTTATCGGCACTTTGCATCGGCCGCGCTCCCGATTCCGGAAGTGCTTGACATTGGGGAATTCAGCGAGAGCCT
GACCTATTGCATCTCCCGCCGTGCACAGGGTGTCACGTTGCAAGACCTGCCTGAAACCGAACTGCCCGCTGTTCTGC
AGCCGGTCGCGGAGGCCATGGATGCGATCGCTGCGGCCGATCTTAGCCAGACGAGCGGGTTCGGCCCATTCGGACCG
CAAGGAATCGGTCAATACACTACATGGCGTGATTTCATATGCGCGATTGCTGATCCCCATGTGTATCACTGGCAAAC
TGTGATGGACGACACCGTCAGTGCGTCCGTCGCGCAGGCTCTCGATGAGCTGATGCTTTGGGCCGAGGACTGCCCCG
AAGTCCGGCACCTCGTGCACGCGGATTTCGGCTCCAACAATGTCCTGACGGACAATGGCCGCATAACAGCGGTCATT
GACTGGAGCGAGGCGATGTTCGGGGATTCCCAATACGAGGTCGCCAACATCTTCTTCTGGAGGCCGTGGTTGGCTTG
TATGGAGCAGCAGACGCGCTACTTCGAGCGGAGGCATCCGGAGCTTGCAGGATCGCCGCGGCTCCGGGCGTATATGC
TCCGCATTGGTCTTGACCAACTCTATCAGAGCTTGGTTGACGGCAATTTCGATGATGCAGCTTGGGCGCAGGGTCGA
TGCGACGCAATCGTCCGATCCGGAGCCGGGACTGTCGGGCGTACACAAATCGCCCGCAGAAGCGCGGCCGTCTGGAC
CGATGGCTGTGTAGAAGTACTCGCCGATAGTGGAAACCGACGCCCCAGCACTCGTCCGAGGGCAAAGGAA**TAG**CACG
TACTACGAGATTTCGATTCCACCGCCGCCTTCTATGAAAGGTTGGGCTTCGGAATCGTTTTCCGGGACGCCGGCTGG
ATGATCCTCCAGCGCGGGGATCTCATGCTGGAGTTCTTCGCCCACCCCAACTTGTTTATTGCAGCTTATAATGGTTA
CAAATAAAGCAATAGCATCACAAATTTCACAAATAAAGCATTTTTTTCACTGCATTCTAGTTGTGGTTTGTCCAAAC
TCATCAATGTATCTTATCATGTCTGTATACCGTCGACaggtggcacttttcggggaaatgtgcgcggaacccctatt
tgtttatttttctaaatacattcaaatatgtatccgctcatgagacaataaccctgataaatgcttcaataatattg
aaaaaggaagagtatgagtattcaacatttccgtgtcgcccttattcccttttttgcggcattttgccttcctgttt
ttgctcacccagaaacgctggtgaaagtaaaagatgctgaagatcagttgggtgcacgagtgggttacatcgaactg
gatctcaacagcggtaagatccttgagagttttcgccccgaagaacgttttccaatgatgagcacttttaaagttct
gctatgtggcgcggtattatcccgtattgacgccgggcaagagcaactcggtcgccgcatacactattctcagaatg
acttggttgagtactcaccagtcacagaaaagcatcttacggatggcatgacagtaagagaattatgcagtgctgcc
ataaccatgagtgataacactgcggccaacttacttctgacaacgatcggaggaccgaaggagctaaccgcttttt
gcacaacatgggggatcatgtaactcgccttgatcgttgggaaccggagctgaatgaagccataccaaacgacgagc
gtgacaccacgatgcctgtagcaatggcaacaacgttgcgcaaactattaactggcgaactacttactctagcttcc
cggcaacaattaatagactggatggaggcggataaagttgcaggaccacttctgcgctcg**gcccttccggc**tggctg
gtttattgctgataaatctggagccggtgagcgtgggtctcgcggtatcattgcagcactggggccagatggtaagc
cctcccgtatcgtagttatctacacgacggggagtcaggcaactatggatgaacgaaatagacagatcgctgagata
ggtgcctcactgattaagcattggtaactgtcagaccaagtttactcatatatactttagattgatttaaa

**D) pAVA2965 [GFP-ENE(WT)-mascRNA, Puromycin$^R$]**

**tDNA1: 1045-1141**
**EF1alfa promoter: 1176-2679**
**5xEGFP: 2702-6439**
**TEV protease: 6440-7165**
**PEST (degradation sequence): 7193-7327**
**Human beta-globin (HBB): 7334-8718**
**MALAT1-ENE(WT)-mascRNA: 8723-8882**
**BGH polyA: 8963-9141**
**tDNA2: 9425-9521**
**FRT: 9532-9558**
**Puromycin$^R$ (lacks promoter and the 1st Methionine): 9755-10584**

acttcatttttaatttaaaaggatctaggtgaagatcctttttgataatctcatgaccaaaatcccttaacgtgagt
tttcgttccactgagcgtcagaccccgtagaaaagatcaaaggatcttcttgagatcctttttttctgcgcgtaatc
tgctgcttgcaaacaaaaaaaccaccgctaccagcggtggtttgtttgccggatcaagagctaccaactctttttcc
gaaggtaactggcttcagcagagcgcagataccaaatactgtccttctagtgtagccgtagttaggccaccacttca
agaactctgtagcaccgcctacatacctcgctctgctaatcctgttaccagtggctgctgccagtggcgataagtcg
tgtcttaccgggttggactcaagacgatagttaccggataaggcgcagcggtcgggctgaacggggggttcgtgcac
acagcccagcttggagcgaacgacctacaccgaactgagatacctacagcgtgagctatgagaaagcgccacgcttc
ccgaagggagaaaggcggacaggtatccggtaagcggcagggtcggaacaggagagcgcacgagggagcttccaggg
ggaaacgcctggtatctttatagtcctgtcgggtttcgccacctctgacttgagcgtcgatttttgtgatgctcgtc
aggggggcggagcctatggaaaaacgccagcaacgcggcctttttacggttcctggccttttgctggccttttgctc
acatgttctttcctgcgttatcccctgattctgtggataaccgtattaccgcctttgagtgagctgataccgctcgc
cgcagccgaacgaccgagcgcagcgagtcagtgagcgaggaagcggaagagcgcccaatacgcaaaccgcctctccc
cgcgcgttggccgattcattaatgcagctggcacgacaggtttcccgactggaaagcgggcagtgagcgcaacgcaa
ttaatgtgagttagctcactcattaggcaccccaggcACGCGAagcgttgttggtttgtagtgcccggtttcgaacc
ggggacctttcgcgtgttaggcgaacgtgataaccactacactacggaaaccaacggtgctagACGCGTattgtcta
ttctgactcggatcCGTACGaattctcatgtttgacagcttatcatcgattagctttggagctaagccagcaatggt
agagggaagattctgcacgtcccttccaggcggcctccccgtcaccaccccccccaacccgccccgaccggagctga
gagtaattcatacaaaaggactcgcccctgccttggggaatcccagggaccgtcgttaaactcccactaacgtagaa
cccagagatcgctgcgttcccgcccctcacccgcccgctctcgtcatcactgaggtggagaagagcatgcgtgagg
ctccggtgcccgtcagtgggcagagcgcacatcgcccacagtccccgagaagttggggggaggggtcggcaattgaa
ccggtgcctagagaaagtggcgcggggtaaactgggaaagtgatgtcgtgtactggctccgcctttttcccgagggt
gggggagaaccgtatataagtgcagtagtcgccgtgaacgttctttttcgcaacgggtttgccgccagaacacaggt
aagtgccgtgtgtggttcccgcgggcctggcctctttacgggttatggcccttgcgtgccttgaattacttccacgc
ccctggctgcagtacgtgattcttgatcccgagcttcgggttggaagtgggtgggagagttcgaggccttgcgctta
aggagccccttcgcctcgtgcttgagttgaggcctggcctgggcgctggggccgccgcgtgcgaatctggtggcacc
ttcgcgcctgtctcgctgctttcgataagtctctagccatttaaaatttttgatgacctgctgcgacgctttttttc
tggcaagatagtcttgtaaatgcgggccaagatctgcacactggtatttcggtttttggggccgcgggcggcgacgg
ggcccgtgcgtcccagcgcacatgttcggcgaggcggggcctgcgagcgcggccaccgagaatcggacgggggtagt
ctcaagctggccggcctgctctggtgcctggcctcgcgccgccgtgtatcgccccgccctgggcggcaaggctgg
cccggtcggcaccagttgcgtgagcggaaagatggccgcttcccggccctgctgcagggagctcaaaatggaggacg
cggcgctcgggagagcgggcgggtgagtcacccacacaaaggaaaagggcctttccgtcctcagccgtcgcttcatg
tgactccacggagtaccgggcgccgtccaggcacctcgattagttctcgagcttttggagtacgtcgtctttaggtt
ggggggaggggttttatgcgatggagtttccccacactgagtgggtggagactgaagttaggccagcttggcacttg
atgtaattctccttggaatttgccctttttgagtttggatcttggttcattctcaagcctcagacagtggttcaaag
tttttttcttccatttcaggtgtcgtgaaaactctagcgtttccgctctagaactagtccaGCTAGCGCTACCGGTC
GCCACCatggtgagcaagggcgaggagctgttcaccggggtggtgcccatcctggtcgagctggacggcgacgtaaa
cggccacaagttcagcgtgtccggcgagggcgagggcgatgccacctacggcaagctgaccctgaagttcatctgca
ccaccggcaagctgcccgtgccctggcccaccctcgtgaccaccctgacctacggcgtgcagtgcttcagccgctac
cccgaccacatgaagcagcacgacttcttcaagtccgccatgcccgaaggctacgtccaggagcgcaccatcttctt
caaggacgacggcaactacaagacccgcgccgaggtgaagttcgagggcgacaccctggtgaaccgcatcgagctga
agggcatcgacttcaaggaggacggcaacatcctggggcacaagctggagtacaactacaacagccacaacgtctat

atcatggccgacaagcagaagaacggcatcaaggtgaacttcaagatccgccacaacatcgaggacggcagcgtgca
gctcgccgaccactaccagcagaacacccccatcggcgacggccccgtgctgctgcccgacaaccactacctgagca
cccagtccgccctgagcaaagaccccaacgagaagcgcgatcacatggtcctgctggagttcgtgaccgccgccggg
atcactctcggcatggacgagctgtacaagGTcactCcgtgTgaaaacctgtacttccaaggtatggtgagcaaggg
cgaggagctgttcaccggggtggtgcccatcctggtcgagctggacggcgacgtaaacggccacaagttcagcgtgt
ccggcgagggcgagggcgatgccacctacggcaagctgaccctgaagttcatctgcaccaccggcaagctgcccgtg
ccctggcccaccctcgtgaccaccctgacctacggcgtgcagtgcttcagccgctaccccgaccacatgaagcagca
cgacttcttcaagtccgccatgcccgaaggctacgtccaggagcgcaccatcttcttcaaggacgacggcaactaca
agacccgcgccgaggtgaagttcgagggcgacaccctggtgaaccgcatcgagctgaagggcatcgacttcaaggag
gacggcaacatcctggggcacaagctggagtacaactacaacagccacaacgtctatatcatggccgacaagcagaa
gaacggcatcaaggtgaacttcaagatccgccacaacatcgaggacggcagcgtgcagctcgccgaccactaccagc
agaacacccccatcggcgacggccccgtgctgctgcccgacaaccactacctgagcacccagtccgccctgagcaaa
gaccccaacgagaagcgcgatcacatggtcctgctggagttcgtgaccgccgccgggatcactctcggcatggacga
gctgtacaagACcactGcgtgCgaaaacctgtacttccaaggtatggtgagcaagggcgaggagctgttcaccgggg
tggtgcccatcctggtcgagctggacggcgacgtaaacggccacaagttcagcgtgtccggcgagggcgagggcgat
gccacctacggcaagctgaccctgaagttcatctgcaccaccggcaagctgcccgtgccctggcccaccctcgtgac
caccctgacctacggcgtgcagtgcttcagccgctaccccgaccacatgaagcagcacgacttcttcaagtccgcca
tgcccgaaggctacgtccaggagcgcaccatcttcttcaaggacgacggcaactacaagacccgcgccgaggtgaag
ttcgagggcgacaccctggtgaaccgcatcgagctgaagggcatcgacttcaaggaggacggcaacatcctggggca
caagctggagtacaactacaacagccacaacgtctatatcatggccgacaagcagaagaacggcatcaaggtgaact
tcaagatccgccacaacatcgaggacggcagcgtgcagctcgccgaccactaccagcagaacacccccatcggcgac
ggccccgtgctgctgcccgacaaccactacctgagcacccagtccgccctgagcaaagaccccaacgagaagcgcga
tcacatggtcctgctggagttcgtgaccgccgccgggatcactctcggcatggacgagctgtacaagGTcactAcgt
gTgaaaacctgtacttccaaggtatggtgagcaagggcgaggagctgttcaccggggtggtgcccatcctggtcgag
ctggacggcgacgtaaacggccacaagttcagcgtgtccggcgagggcgagggcgatgccacctacggcaagctgac
cctgaagttcatctgcaccaccggcaagctgcccgtgccctggcccaccctcgtgaccaccctgacctacggcgtgc
agtgcttcagccgctaccccgaccacatgaagcagcacgacttcttcaagtccgccatgcccgaaggctacgtccag
gagcgcaccatcttcttcaaggacgacggcaactacaagacccgcgccgaggtgaagttcgagggcgacaccctggt
gaaccgcatcgagctgaagggcatcgacttcaaggaggacggcaacatcctggggcacaagctggagtacaactaca
acagccacaacgtctatatcatggccgacaagcagaagaacggcatcaaggtgaacttcaagatccgccacaacatc
gaggacggcagcgtgcagctcgccgaccactaccagcagaacacccccatcggcgacggccccgtgctgctgcccga
caaccactacctgagcacccagtccgccctgagcaaagaccccaacgagaagcgcgatcacatggtcctgctggagt
tcgtgaccgccgccgggatcactctcggcatggacgagctgtacaagACcactTcgtgCgaaaacctgtacttccaa
ggtatggtgagcaagggcgaggagctgttcaccggggtggtgcccatcctggtcgagctggacggcgacgtaaacgg
ccacaagttcagcgtgtccggcgagggcgagggcgatgccacctacggcaagctgaccctgaagttcatctgcacca
ccggcaagctgcccgtgccctggcccaccctcgtgaccaccctgacctacggcgtgcagtgcttcagccgctacccc
gaccacatgaagcagcacgacttcttcaagtccgccatgcccgaaggctacgtccaggagcgcaccatcttcttcaa
ggacgacggcaactacaagacccgcgccgaggtgaagttcgagggcgacaccctggtgaaccgcatcgagctgaagg
gcatcgacttcaaggaggacggcaacatcctggggcacaagctggagtacaactacaacagccacaacgtctatatc
atggccgacaagcagaagaacggcatcaaggtgaacttcaagatccgccacaacatcgaggacggcagcgtgcagct
cgccgaccactaccagcagaacacccccatcggcgacggccccgtgctgctgcccgacaaccactacctgagcaccc
agtccgccctgagcaaagaccccaacgagaagcgcgatcacatggtcctgctggagttcgtgaccgccgccgggatc
actctcggcatggacgagctgtacaaggaaaacctgtacttccaaAgtggagaAAGCTTgtttaaggaccacgtga
ttacaacccgatatcgagcaccatttgtcatttgacgaatgaatctgatgggcacacaacatcgttgtatggtattg
gatttggtcccttcatcattacaaacaagcacttgtttagaagaaataatggaacactgttggtccaatcactacat
ggtgtattcaaggtcaagaacaccacgactttgcaacaacacctcattgatgggagggacatgataattattcgcat
gcctaaggatttcccaccatttcctcaaaagctgaaatttagagagccacaaagggaagagcgcatatgtcttgtga
caaccaacttccaaactaagagcatgtctagcatggtgtcagacactagttgcacattcccttcatctgatggcata
ttctggaagcattggattcaaaccaaggatgggcagtgtggcagtccattagtatcaactagagatgggttcattgt
tggtatacactcagcatcgaatttcaccaacacaaacaattatttcacaagcgtgccgaaaaacttcatggaattgt
tgacaaatcaggaggcgcagcagtgggttagtggttggcgattaaatgctgactcagtattgtgggggggccataaa
gttttcatgagcaaacctgaagagccttttcagccagttaaggaagcgactcaactcatgaatgaattggtgtactc
gcaagaaaacctgtacttccaaagtATGCATAGATCACGAGATATCAGCCATGGCTTCCCGCCGGCGGTGGCGGCGC
AGGATGATGGCACGCTGCCCATGTCTTGTGCCCAGGAGAGCGGGATGGACCGTCACCCTGCAGCCTGTGCTTCTGCT
AGGATCAATGTGGAGCTCGTGCACCTGACTCCTGAGGAGAAGTCTGCCGTTACTGCCCTGTGGGGCAAGGTGAACGT
GGATGAAGTTGGTGGTGAGGCCCTGGGCAGGCTGCTGGTGGTCTACCCTTGGACCCAGAGGTTCTTTGAGTCCTTTG
GGGATCTGTCCACTCCTGATGCTGTTATGGGCAACCCTAAGGTGAAGGCTCATGGCAAGAAAGTGCTCGGTGCCTTT

AGTGATGGCCTGGCTCACCTGGACAACCTCAAGGGCACCTTTGCCACACTGAGTGAGCTGCACTGTGACAAGCTGCA
CGTGGATCCTGAGAACTTCAGGGTGAGTCTATGGGACCCTTGATGTTTTCTTTCCCCTTCTTTTCTATGGTTAAGTT
CATGTCATAGGAAGGGGATAAGTAACAGGGTACAGTTTAGAATGGGAAACAGACGAATGATTGCATCAGTGTGGAAG
TCTCAGGATCGTTTTAGTTTCTTTTATTTGCTGTTCATAACAATTGTTTTCTTTTGTTTAATTCTTGCTTTCTTTTT
TTTTCTTCTCCGCAATTTTTACTATTATACTTAATGCCTTAACATTGTGTATAACAAAAGGAAATATCTCTGAGATA
CATTAAGTAACTTAAAAAAAAAACTTTACACAGTCTGCCTAGTACATTACTATTTGGAATATATGTGTGCTTATTTG
CATATTCATAATCTCCCTACTTTATTTTCTTTTATTTTTAATTGATACATAATCATTATACATATTTATGGGTTAAA
GTGTAATGTTTTAATATGTGTACACATATTGACCAAATCAGGGTAATTTTGCATTTGTAATTTTAAAAAATGCTTTC
TTCTTTTAATATACTTTTTTGTTTATCTTATTTCTAATACTTTCCCTAATCTCTTTCTTTCAGGGCAATAATGATAC
AATGTATCATGCCTCTTTGCACCATTCTAAAGAATAACAGTGATAATTTCTGGGTTAAGGCAATAGCAATATTTCTG
CATATAAATATTTCTGCATATAAATTGTAACTGATGTAAGAGGTTTCATATTGCTAATAGCAGCTACAATCCAGCTA
CCATTCTGCTTTTATTTTATGGTTGGGATAAGGCTGGATTATTCTGAGT**CCAAGCTAGGCCCTTTTGCTAATCATG**T
TCATACCTCTTATCTTCCTCCCACAGCTCCTGGGCAACGTGCTGGTCTGTGTGCTGGCCCATCACTTTGGCAAAGAA
TTCACCCCACCAGTGCAGGCTGCCTATCAGAAAGTGGTGGCTGGTGTGGCTAATGCCCTGGCCCACAAGTATCAC**TA**
**A**GC**GGCCGC**TCGCTTTCTTGCTGTCCAATTTCTATTAAAGGTTCCTTTGTTCCCTAAGTCCAACTACTAAACTGGGG
GATATTATGAAGGGCCTCGAGtagggtcatgaaggtttttcttttcctgagaaaacaacacgtattgttttctcagg
ttttgcttttggcctttttctagcttaaaaaaaaaaaaagcaaaagatgctggtggttggcactcctggtttccag
gacggggttcaaatccctgcggcgtctttgctttgactCTCGAGGCT**GATC**A***GCCTCGA******CTGTGCCTTCTAG***TTGCC
AGCCATCTGTTGTTTGCCCCTCCCCCGTGCCTTCCTTGACCCTGGAAGGTGCCACTCCCACTGTCCTTTCCTAATAA
AATGAGGAAATTGCATCGCATTGTCTGAGTAGGTGTCATTCTATTCTGGGGGGTGGGGTGGGGCAGGACAGCAAGGG
GGAGGATTGGGAAGACAATAGCAGGCATGCTGGGGATGCGGTGGGCTCTATGGCTTCTGAGGCGGAAAGAACCAGCT
GGGGCTCTAGGGGGTATCCCCACGCGCCCTGTAGCGGCGCATTAAGCGCGGCGGGTGTGGTGGTTACGCGCAGCGTG
ACCGCTACACTTGCCAGCGCCCTAGCGCCCGCTCCTTTCGCTTTCTTCCCTTCCTTTCTCGCCACGTTCGCCGGCTT
TCCCCGTCAAGCTCTAAATCGGGGGCTCCCTTTAGGGTTCCGATTTAGTGCTTTACGGCACCTCGACCCCAAAAAAC
TTGATTAGGGTGATGGTTCACGTAGGTAC**G**ctagcaccgttggtttccgtagtgtagtggttatcacgttcgcctaa
cacgcgaaaggtccccggttcgaaaccgggcactacaaaccaacaacgctGGTACCCCTA*GAAGTTCCTATTCCGAA*
*GTTCCTATTC****TCTAGAAAGTATAGGAACTTC***CTTGGCCAAAAAGCCTGAAagatcacgagatatcagccatggcttc
ccgccggcggtggcggcgcaggatgatggcacgctgcccaCgtcttgtgcccaggagagcgggaCggaccgtcaccc
tgcagcctgtgcttctgctaggatcaatgtggaaaacctgtacttccaaGgtaccgagtacaagcccacggtgcgcc
tcgccacccgcgacgacgtccccagggccgtCcgcaccctcgccgccgcgttcgccgactaccccgccacgcgccac
accgtcgatccggaccgccacatcgagcg*ggtcaccgagctgcaagaactcttcctcacgcgcgtcgggctcgacat*
*cggcaaggtgtgggtcgcggacgacggcgccgcggtggcggtctggaccacgccggagagcgtcgaagcgggggcgg*
*tgttcgccgagatcggcccgcgcatggccgagttgagcggttcccggctggccgcgcagcaacagatggaaggcctc*
*ctggcgccgcaccggcccaaggagcccgcgtggttcctggccaccgtcggcgtctcgcccgaccaccagggcaaggg*
*tctgggcagcgccgtcgtgctccccggagtggaggcggccgagcgcgccggggtgcccgccttcctggagacctccg*
*cgccccgcaacctccccttctacg*agcggctcggcttcaccgtcaccgccgacgtcgaggtgcccgaaggaccgcg**c**
**acctggtg**catgacccgcaagcccggtgcc**tga**ggtaccgctcgctgatcagcctcgactgtgccttctagttgcca
gccatctgttgtttgcccctcccccgtgccttccttgaccctggaaggtgccactcccactgtcctttcctaataaa
atgaggaaattgcatcgcattgtctgagtaggtgtcattctattctggggggtggggtggggcaggacagcaagggg
gaggattgggaagacaatagcaggcatgctggggaGTCGACaggtggcacttttcggggaaatgtgcgcggaacccc
tatttgtttatttttctaaatacattcaaatatgtatccgctcatgagacaataaccctgataaatgcttcaataat
attgaaaaaggaagagtatgagtattcaacatttccgtgtcgcccttattcccttttttgcggcattttgccttcct
gtttttgctcacccagaaacgctggtgaaagtaaaagatgctgaagatcagttgggtgcacgagtgggttacatcga
actggatctcaacagcggtaagatccttgagagttttcgccccgaagaacgttttccaatgatgagcacttttaaag
ttctgctatgtggcgcggtattatcccgtattgacgccgggcaagagcaactcggtcgccgcatacactattctcag
aatgacttggttgagtactcaccagtcacagaaaagcatcttacggatggcatgacagtaagagaattatgcagtgc
tgccataaccatgagtgataacactgcggccaacttacttctgacaacgatcggaggaccgaaggagctaaccgctt
ttttgcacaacatgggggatcatgtaactcgccttgatcgttgggaaccggagctgaatgaagccataccaaacgac
gagcgtgacaccacgatgcctgtagcaatggcaacaacgttgcgcaaactattaactggcgaactacttactctagc
ttcccggcaacaattaatagactggatggaggcggataaagttgcaggaccacttctgcgctcg**gcccttccggc**tg
gctggtttattgctgataaatctggagccggtgagcgtgggtctcgcggtatcattgcagcactggggccagatggt
aagccctcccgtatcgtagttatctacacgacggggagtcaggcaactatggatgaacgaaatagacagatcgctga
gataggtgcctcactgattaagcattggtaactgtcagaccaagtttactcatatatactttagattgatttaa

**E) pAVA3871 [GFP-ENE(C8351G)-mascRNA(mut 8356-8370), Puromycin$^R$]**

**tDNA1: 1045-1141**
**EF1alfa promoter: 1176-2679**
**5xEGFP: 2702-6439**
**TEV protease: 6440-7165**
**PEST (degradation sequence): 7193-7327**
**Human beta-globin (HBB): 7334-8718**
**MALAT1-ENE(C8351G)-mascRNA(mut 8356-8370): 8723-8882**
**BGH polyA: 8963-9141**
**tDNA2: 9425-9521**
**FRT: 9532-9558**
**Puromycin$^R$ (lacks promoter and the 1$^{st}$ Methionine): 9755-10584**

acttcattttaatttaaaaggatctaggtgaagatcctttttgataatctcatgaccaaaatcccttaacgtgagt
tttcgttccactgagcgtcagaccccgtagaaaagatcaaaggatcttcttgagatcctttttttctgcgcgtaatc
tgctgcttgcaaacaaaaaaaccaccgctaccagcggtggtttgtttgccggatcaagagctaccaactctttttcc
gaaggtaactggcttcagcagagcgcagataccaaatactgtccttctagtgtagccgtagttaggccaccacttca
agaactctgtagcaccgcctacatacctcgctctgctaatcctgttaccagtggctgctgccagtggcgataagtcg
tgtcttaccgggttggactcaagacgatagttaccggataaggcgcagcggtcgggctgaacggggggttcgtgcac
acagcccagcttggagcgaacgacctacaccgaactgagatacctacagcgtgagctatgagaaagcgccacgcttc
ccgaagggagaaaggcggacaggtatccggtaagcggcagggtcggaacaggagagcgcacgagggagcttccaggg
ggaaacgcctggtatctttatagtcctgtcgggtttcgccacctctgacttgagcgtcgatttttgtgatgctcgtc
aggggggcggagcctatggaaaaacgccagcaacgcggcctttttacggttcctggccttttgctggccttttgctc
acatgttctttcctgcgttatcccctgattctgtggataaccgtattaccgcctttgagtgagctgataccgctcgc
cgcagccgaacgaccgagcgcagcgagtcagtgagcgaggaagcggaagagcgcccaatacgcaaaccgcctctccc
cgcgcgttggccgattcattaatgcagctggcacgacaggtttcccgactggaaagcgggcagtgagcgcaacgcaa
ttaatgtgagttagctcactcattaggcaccccaggcACGCGAagcgttgttggtttgtagtgcccggtttcgaacc
ggggacctttcgcgtgttaggcgaacgtgataaccactacactacggaaaccaacggtgctagACGCGTattgtcta
ttctgactcggatcCGTACGaattctcatgtttgacagcttatcatcgattagctttggagctaagccagcaatggt
agagggaagattctgcacgtcccttccaggcggcctccccgtcaccaccccccccaacccgccccgaccggagctga
gagtaattcatacaaaaggactcgcccctgccttggggaatcccagggaccgtcgttaaactcccactaacgtagaa
cccagagatcgctgcgttcccgcccctcacccgcccgctctcgtcatcactgaggtggagaagagcatgcgtgagg
ctccggtgcccgtcagtgggcagagcgcacatcgcccacagtccccgagaagttggggggaggggtcggcaattgaa
ccggtgcctagagaaagtggcgcggggtaaactgggaaagtgatgtcgtgtactggctccgcctttttcccgagggt
gggggagaaccgtatataagtgcagtagtcgccgtgaacgttctttttcgcaacgggtttgccgccagaacacaggt
aagtgccgtgtgtggttcccgcgggcctggcctctttacgggttatggcccttgcgtgccttgaattacttccacgc
ccctggctgcagtacgtgattcttgatcccgagcttcgggttggaagtgggtgggagagttcgaggccttgcgctta
aggagccccttcgcctcgtgcttgagttgaggcctggcctgggcgctggggccgccgcgtgcgaatctggtggcacc
ttcgcgcctgtctcgctgctttcgataagtctctagccatttaaaatttttgatgacctgctgcgacgctttttttc
tggcaagatagtcttgtaaatgcgggccaagatctgcacactggtatttcggtttttggggccgcgggcggcgacgg
ggcccgtgcgtcccagcgcacatgttcggcgaggcggggcctgcgagcgcggccaccgagaatcggacgggggtagt
ctcaagctggccggcctgctctggtgcctggcctcgcgccgccgtgtatcgccccgccctgggcggcaaggctgg
cccggtcggcaccagttgcgtgagcggaaagatggccgcttcccggccctgctgcagggagctcaaaatggaggacg
cggcgctcgggagagcgggcgggtgagtcacccacacaaaggaaaagggcctttccgtcctcagccgtcgcttcatg
tgactccacggagtaccgggcgccgtccaggcacctcgattagttctcgagcttttggagtacgtcgtctttaggtt
ggggggaggggttttatgcgatggagtttccccacactgagtgggtggagactgaagttaggccagcttggcacttg
atgtaattctccttggaatttgccctttttgagtttggatcttggttcattctcaagcctcagacagtggttcaaag
tttttttcttccatttcaggtgtcgtgaaaactctagcgtttccgctctagaactagtccaGCTAGCGCTACCGGTC
GCCACCatggtgagcaagggcgaggagctgttcaccggggtggtgcccatcctggtcgagctggacggcgacgtaaa
cggccacaagttcagcgtgtccggcgagggcgagggcgatgccacctacggcaagctgaccctgaagttcatctgca
ccaccggcaagctgcccgtgccctggcccaccctcgtgaccaccctgacctacggcgtgcagtgcttcagccgctac
cccgaccacatgaagcagcacgacttcttcaagtccgccatgcccgaaggctacgtccaggagcgcaccatcttctt
caaggacgacggcaactacaagacccgcgccgaggtgaagttcgagggcgacaccctggtgaaccgcatcgagctga
agggcatcgacttcaaggaggacggcaacatcctggggcacaagctggagtacaactacaacagccacaacgtctat

atcatggccgacaagcagaagaacggcatcaaggtgaacttcaagatccgccacaacatcgaggacggcagcgtgca
gctcgccgaccactaccagcagaacacccccatcggcgacggccccgtgctgctgcccgacaaccactacctgagca
cccagtccgccctgagcaaagaccccaacgagaagcgcgatcacatggtcctgctggagttcgtgaccgccgccggg
atcactctcggcatggacgagctgtacaagGTcactCcgtgTgaaaacctgtacttccaaggtatggtgagcaaggg
cgaggagctgttcaccggggtggtgcccatcctggtcgagctggacggcgacgtaaacggccacaagttcagcgtgt
ccggcgagggcgagggcgatgccacctacggcaagctgaccctgaagttcatctgcaccaccggcaagctgcccgtg
ccctggcccaccctcgtgaccaccctgacctacggcgtgcagtgcttcagccgctaccccgaccacatgaagcagca
cgacttcttcaagtccgccatgcccgaaggctacgtccaggagcgcaccatcttcttcaaggacgacggcaactaca
agacccgcgccgaggtgaagttcgagggcgacaccctggtgaaccgcatcgagctgaagggcatcgacttcaaggag
gacggcaacatcctggggcacaagctggagtacaactacaacagccacaacgtctatatcatggccgacaagcagaa
gaacggcatcaaggtgaacttcaagatccgccacaacatcgaggacggcagcgtgcagctcgccgaccactaccagc
agaacacccccatcggcgacggccccgtgctgctgcccgacaaccactacctgagcacccagtccgccctgagcaaa
gaccccaacgagaagcgcgatcacatggtcctgctggagttcgtgaccgccgccgggatcactctcggcatggacga
gctgtacaagACcactGcgtgCgaaaacctgtacttccaaggtatggtgagcaagggcgaggagctgttcaccgggg
tggtgcccatcctggtcgagctggacggcgacgtaaacggccacaagttcagcgtgtccggcgagggcgagggcgat
gccacctacggcaagctgaccctgaagttcatctgcaccaccggcaagctgcccgtgccctggcccaccctcgtgac
caccctgacctacggcgtgcagtgcttcagccgctaccccgaccacatgaagcagcacgacttcttcaagtccgcca
tgcccgaaggctacgtccaggagcgcaccatcttcttcaaggacgacggcaactacaagacccgcgccgaggtgaag
ttcgagggcgacaccctggtgaaccgcatcgagctgaagggcatcgacttcaaggaggacggcaacatcctggggca
caagctggagtacaactacaacagccacaacgtctatatcatggccgacaagcagaagaacggcatcaaggtgaact
tcaagatccgccacaacatcgaggacggcagcgtgcagctcgccgaccactaccagcagaacacccccatcggcgac
ggccccgtgctgctgcccgacaaccactacctgagcacccagtccgccctgagcaaagaccccaacgagaagcgcga
tcacatggtcctgctggagttcgtgaccgccgccgggatcactctcggcatggacgagctgtacaagGTcactAcgt
gTgaaaacctgtacttccaaggtatggtgagcaagggcgaggagctgttcaccggggtggtgcccatcctggtcgag
ctggacggcgacgtaaacggccacaagttcagcgtgtccggcgagggcgagggcgatgccacctacggcaagctgac
cctgaagttcatctgcaccaccggcaagctgcccgtgccctggcccaccctcgtgaccaccctgacctacggcgtgc
agtgcttcagccgctaccccgaccacatgaagcagcacgacttcttcaagtccgccatgcccgaaggctacgtccag
gagcgcaccatcttcttcaaggacgacggcaactacaagacccgcgccgaggtgaagttcgagggcgacaccctggt
gaaccgcatcgagctgaagggcatcgacttcaaggaggacggcaacatcctggggcacaagctggagtacaactaca
acagccacaacgtctatatcatggccgacaagcagaagaacggcatcaaggtgaacttcaagatccgccacaacatc
gaggacggcagcgtgcagctcgccgaccactaccagcagaacacccccatcggcgacggccccgtgctgctgcccga
caaccactacctgagcacccagtccgccctgagcaaagaccccaacgagaagcgcgatcacatggtcctgctggagt
tcgtgaccgccgccgggatcactctcggcatggacgagctgtacaagACcactTcgtgCgaaaacctgtacttccaa
ggtatggtgagcaagggcgaggagctgttcaccggggtggtgcccatcctggtcgagctggacggcgacgtaaacgg
ccacaagttcagcgtgtccggcgagggcgagggcgatgccacctacggcaagctgaccctgaagttcatctgcacca
ccggcaagctgcccgtgccctggcccaccctcgtgaccaccctgacctacggcgtgcagtgcttcagccgctacccc
gaccacatgaagcagcacgacttcttcaagtccgccatgcccgaaggctacgtccaggagcgcaccatcttcttcaa
ggacgacggcaactacaagacccgcgccgaggtgaagttcgagggcgacaccctggtgaaccgcatcgagctgaagg
gcatcgacttcaaggaggacggcaacatcctggggcacaagctggagtacaactacaacagccacaacgtctatatc
atggccgacaagcagaagaacggcatcaaggtgaacttcaagatccgccacaacatcgaggacggcagcgtgcagct
cgccgaccactaccagcagaacacccccatcggcgacggccccgtgctgctgcccgacaaccactacctgagcaccc
agtccgccctgagcaaagaccccaacgagaagcgcgatcacatggtcctgctggagttcgtgaccgccgccgggatc
actctcggcatggacgagctgtacaaggaaaacctgtacttccaaAgtggagaAAGCTTgtttaaggaccacgtga
ttacaacccgatatcgagcaccatttgtcatttgacgaatgaatctgatgggcacacaacatcgttgtatggtattg
gatttggtcccttcatcattacaaacaagcacttgtttagaagaaataatggaacactgttggtccaatcactacat
ggtgtattcaaggtcaagaacaccacgactttgcaacaacacctcattgatgggagggacatgataattattcgcat
gcctaaggatttcccaccatttcctcaaaagctgaaatttagagagccacaaagggaagagcgcatatgtcttgtga
caaccaacttccaaactaagagcatgtctagcatggtgtcagacactagttgcacattcccttcatctgatggcata
ttctggaagcattggattcaaaccaaggatgggcagtgtggcagtccattagtatcaactagagatgggttcattgt
tggtatacactcagcatcgaatttcaccaacacaaacaattatttcacaagcgtgccgaaaaacttcatggaattgt
tgacaaatcaggaggcgcagcagtgggttagtggttggcgattaaatgctgactcagtattgtgggggggccataaa
gttttcatgagcaaacctgaagagccttttcagccagttaaggaagcgactcaactcatgaatgaattggtgtactc
gcaagaaaacctgtacttccaaagtATGCATAGATCACGAGATATCAGCCATGGCTTCCCGCCGGCGGTGGCGGCGC
AGGATGATGGCACGCTGCCCATGTCTTGTGCCCAGGAGAGCGGGATGGACCGTCACCCTGCAGCCTGTGCTTCTGCT
AGGATCAATGTGGAGCTCGTGCACCTGACTCCTGAGGAGAAGTCTGCCGTTACTGCCCTGTGGGGCAAGGTGAACGT
GGATGAAGTTGGTGGTGAGGCCCTGGGCAGGCTGCTGGTGGTCTACCCTTGGACCCAGAGGTTCTTTGAGTCCTTTG
GGGATCTGTCCACTCCTGATGCTGTTATGGGCAACCCTAAGGTGAAGGCTCATGGCAAGAAAGTGCTCGGTGCCTTT

AGTGATGGCCTGGCTCACCTGGACAACCTCAAGGGCACCTTTGCCACACTGAGTGAGCTGCACTGTGACAAGCTGCA
CGTGGATCCTGAGAACTTCAGGGTGAGTCTATGGGACCCTTGATGTTTTCTTTCCCCTTCTTTTCTATGGTTAAGTT
CATGTCATAGGAAGGGGATAAGTAACAGGGTACAGTTTAGAATGGGAAACAGACGAATGATTGCATCAGTGTGGAAG
TCTCAGGATCGTTTTAGTTTCTTTTATTTGCTGTTCATAACAATTGTTTTCTTTTGTTTAATTCTTGCTTTCTTTTT
TTTTCTTCTCCGCAATTTTTACTATTATACTTAATGCCTTAACATTGTGTATAACAAAAGGAAATATCTCTGAGATA
CATTAAGTAACTTAAAAAAAAAACTTTACACAGTCTGCCTAGTACATTACTATTTGGAATATATGTGTGCTTATTTG
CATATTCATAATCTCCCTACTTTATTTTCTTTTATTTTTAATTGATACATAATCATTATACATATTTATGGGTTAAA
GTGTAATGTTTTAATATGTGTACACATATTGACCAAATCAGGGTAATTTTGCATTTGTAATTTTAAAAAATGCTTTC
TTCTTTTAATATACTTTTTTGTTTATCTTATTTCTAATACTTTCCCTAATCTCTTTCTTTCAGGGCAATAATGATAC
AATGTATCATGCCTCTTTGCACCATTCTAAAGAATAACAGTGATAATTTCTGGGTTAAGGCAATAGCAATATTTCTG
CATATAAATATTTCTGCATATAAATTGTAACTGATGTAAGAGGTTTCATATTGCTAATAGCAGCTACAATCCAGCTA
CCATTCTGCTTTTATTTTATGGTTGGGATAAGGCTGGATTATTCTGAGTCCAAGCTAGGCCCTTTTGCTAATCATGT
TCATACCTCTTATCTTCCTCCCACAGCTCCTGGGCAACGTGCTGGTCTGTGTGCTGGCCCATCACTTTGGCAAAGAA
TTCACCCCACCAGTGCAGGCTGCCTATCAGAAAGTGGTGGCTGGTGTGGCTAATGCCCTGGCCCACAAGTATCACTA
AGCGGCCGCTCGCTTTCTTGCTGTCCAATTTCTATTAAAGGTTCCTTTGTTCCCTAAGTCCAACTACTAAACTGGGG
GATATTATGAAGGGCCTCGAGtagggtcatgaaggtttttcttttcctgagaaaacaacacgtattgttttctcagg
ttttgctttttggcctttttctagcttaaaaaaaaaaaaaggaaaactacgaccaccaacccactcctggtttccag
gacggggttcaaatccctgcggcgtctttgctttgactCTCGAGGCTGATCAGCCTCGACTGTGCCTTCTAGTTGCC
AGCCATCTGTTGTTTGCCCCTCCCCCGTGCCTTCCTTGACCCTGGAAGGTGCCACTCCCACTGTCCTTTCCTAATAA
AATGAGGAAATTGCATCGCATTGTCTGAGTAGGTGTCATTCTATTCTGGGGGGTGGGGTGGGGCAGGACAGCAAGGG
GGAGGATTGGGAAGACAATAGCAGGCATGCTGGGGATGCGGTGGGCTCTATGGCTTCTGAGGCGGAAAGAACCAGCT
GGGGCTCTAGGGGGTATCCCCACGCGCCCTGTAGCGGCGCATTAAGCGCGGCGGGTGTGGTGGTTACGCGCAGCGTG
ACCGCTACACTTGCCAGCGCCCTAGCGCCCGCTCCTTTCGCTTTCTTCCCTTCCTTTCTCGCCACGTTCGCCGGCTT
TCCCCGTCAAGCTCTAAATCGGGGGCTCCCTTTAGGGTTCCGATTTAGTGCTTTACGGCACCTCGACCCCAAAAAAC
TTGATTAGGGTGATGGTTCACGTAGGTACGctagcaccgttggtttccgtagtgtagtggttatcacgttcgcctaa
cacgcgaaaggtccccggttcgaaaccgggcactacaaaccaacaacgctGGTACCCCTAGAAGTTCCTATTCCGAA
GTTCCTATTCTCTAGAAAGTATAGGAACTTCCTTGGCCAAAAAGCCTGAAagatcacgagatatcagccatggcttc
ccgccggcggtggcggcgcaggatgatggcacgctgcccaCgtcttgtgcccaggagagcgggaCggaccgtcaccc
tgcagcctgtgcttctgctaggatcaatgtggaaaacctgtacttccaaGgtaccgagtacaagcccacggtgcgcc
tcgccacccgcgacgacgtccccagggccgtCcgcaccctcgccgccgcgttcgccgactaccccgccacgcgccac
accgtcgatccggaccgccacatcgagcgggtcaccgagctgcaagaactcttcctcacgcgcgtcgggctcgacat
cggcaaggtgtgggtcgcggacgacggcgccgcggtggcggtctggaccacgccggagagcgtcgaagcgggggcgg
tgttcgccgagatcggcccgcgcatggccgagttgagcggttcccggctggccgcgcagcaacagatggaaggcctc
ctggcgccgcaccggcccaaggagcccgcgtggttcctggccaccgtcggcgtctcgcccgaccaccagggcaaggg
tctgggcagcgccgtcgtgctccccggagtggaggcggccgagcgcgccggggtgcccgccttcctggagacctccg
cgccccgcaacctccccttctacgagcggctcggcttcaccgtcaccgccgacgtcgaggtgcccgaaggaccgcgc
acctggtgcatgacccgcaagcccggtgcctgaggtaccgctcgctgatcagcctcgactgtgccttctagttgcca
gccatctgttgtttgcccctcccccgtgccttccttgaccctggaaggtgccactcccactgtcctttcctaataaa
atgaggaaattgcatcgcattgtctgagtaggtgtcattctattctggggggtggggtggggcaggacagcaagggg
gaggattgggaagacaatagcaggcatgctggggaGTCGACaggtggcacttttcggggaaatgtgcgcggaacccc
tatttgtttatttttctaaatacattcaaatatgtatccgctcatgagacaataaccctgataaatgcttcaataat
attgaaaaaggaagagtatgagtattcaacatttccgtgtcgcccttattcccttttttgcggcattttgccttcct
gtttttgctcacccagaaacgctggtgaaagtaaaagatgctgaagatcagttgggtgcacgagtgggttacatcga
actggatctcaacagcggtaagatccttgagagttttcgccccgaagaacgttttccaatgatgagcacttttaaag
ttctgctatgtggcgcggtattatcccgtattgacgccgggcaagagcaactcggtcgccgcatacactattctcag
aatgacttggttgagtactcaccagtcacagaaaagcatcttacggatggcatgacagtaagagaattatgcagtgc
tgccataaccatgagtgataacactgcggccaacttacttctgacaacgatcggaggaccgaaggagctaaccgctt
ttttgcacaacatgggggatcatgtaactcgccttgatcgttgggaaccggagctgaatgaagccataccaaacgac
gagcgtgacaccacgatgcctgtagcaatggcaacaacgttgcgcaaactattaactggcgaactacttactctagc
ttcccggcaacaattaatagactggatggaggcggataaagttgcaggaccacttctgcgctcggcccttccggctg
gctggtttattgctgataaatctggagccggtgagcgtgggtctcgcggtatcattgcagcactggggccagatggt
aagccctcccgtatcgtagttatctacacgacggggagtcaggcaactatggatgaacgaaatagacagatcgctga
gataggtgcctcactgattaagcattggtaactgtcagaccaagtttactcatatatactttagattgatttaaa

**F) pAVA3874 [GFP-ENE(WT)-mascRNA(mut 8356-8370), Puromycin$^R$]**

**tDNA1: 1045-1141**
**EF1alfa promoter: 1176-2679**
**5xEGFP: 2702-6439**
**TEV protease: 6440-7165**
**PEST (degradation sequence): 7193-7327**
**Human beta-globin (HBB): 7334-8718**
**MALAT1-ENE(WT)-mascRNA(mut 8356-8370): 8723-8882**
**BGH polyA: 8963-9141**
**tDNA2: 9425-9521**
**FRT: 9532-9558**
**Puromycin$^R$ (lacks promoter and the 1$^{st}$ Methionine): 9755-10584**

acttcattttaatttaaaaggatctaggtgaagatcctttttgataatctcatgaccaaaatcccttaacgtgagt
tttcgttccactgagcgtcagaccccgtagaaaagatcaaaggatcttcttgagatcctttttttctgcgcgtaatc
tgctgcttgcaaacaaaaaaaccaccgctaccagcggtggtttgtttgccggatcaagagctaccaactctttttcc
gaaggtaactggcttcagcagagcgcagataccaaatactgtccttctagtgtagccgtagttaggccaccacttca
agaactctgtagcaccgcctacatacctcgctctgctaatcctgttaccagtggctgctgccagtggcgataagtcg
tgtcttaccgggttggactcaagacgatagttaccggataaggcgcagcggtcgggctgaacggggggttcgtgcac
acagcccagcttggagcgaacgacctacaccgaactgagatacctacagcgtgagctatgagaaagcgccacgcttc
ccgaagggagaaaggcggacaggtatccggtaagcggcagggtcggaacaggagagcgcacgagggagcttccaggg
ggaaacgcctggtatctttatagtcctgtcgggtttcgccacctctgacttgagcgtcgatttttgtgatgctcgtc
aggggggcggagcctatggaaaaacgccagcaacgcggcctttttacggttcctggccttttgctggccttttgctc
acatgttctttcctgcgttatcccctgattctgtggataaccgtattaccgcctttgagtgagctgataccgctcgc
cgcagccgaacgaccgagcgcagcgagtcagtgagcgaggaagcggaagagcgcccaatacgcaaaccgcctctccc
cgcgcgttggccgattcattaatgcagctggcacgacaggtttcccgactggaaagcgggcagtgagcgcaacgcaa
ttaatgtgagttagctcactcattaggcaccccaggcACGCGAagcgttgttggtttgtagtgcccggtttcgaacc
ggggacctttcgcgtgttaggcgaacgtgataaccactacactacggaaaccaacggt**gctag**ACGCGTattgtcta
ttctgactcggatcCGTACG**aattctcatgtttgacagcttatcatcgat*tagctt*tggagctaagccagcaatggt**
**agagggaagattctgcacgtcccttccaggcggcctccccgtcaccaccccccccaacccgccccgaccggagctga**
**gagtaattcatacaaaaggactcgcccctgccttggggaatcccagggaccgtcgttaaactcccactaacgtagaa**
**cccagagatcgctgcgttcccgccccctcacccgcccgctctcgtcatcactgaggtggagaagagcatgcgtgagg**
**ctccggtgcccgtcagtgggcagagcgcacatcgcccacagtccccgagaagttggggggaggggtcggcaattgaa**
**ccggtgcctagagaaagtggcgcggggtaaactgggaaagtgatgtcgtgtactggctccgcctttttcccgagggt**
**gggggagaaccgtatataagtgcagtagtcgccgtgaacgttctttttcgcaacgggtttgccgccagaacacaggt**
**aagtgccgtgtgtggttcccgcgggcctggcctctttacgggttatggcccttgcgtgccttgaattacttccacgc**
**ccctggctgcagtacgtgattcttgatcccgagcttcgggttggaagtgggtgggagagttcgaggccttgcgctta**
**aggagccccttcgcctcgtgcttgagttgaggcctggcctgggcgctggggccgccgcgtgcgaatctggtggcacc**
**ttcgcgcctgtctcgctgctttcgataagtctctagccatttaaaatttttgatgacctgctgcgacgctttttttc**
**tggcaagatagtcttgtaaatgcgggccaagatctgcacactggtatttcggtttttggggccgcgggcggcgacgg**
**ggcccgtgcgtcccagcgcacatgttcggcgaggcggggcctgcgagcgcggccaccgagaatcggacgggggtagt**
**ctcaagctggccggcctgctctggtgcctggcctcgcgccgccgtgtatcgccccgccctgggcggcaacaggctgg**
**cccggtcggcaccagttgcgtgagcggaaagatggccgcttcccggccctgctgcagggagctcaaaatggaggacg**
**cggcgctcgggagagcgggcgggtgagtcacccacacaaaggaaaagggcctttccgtcctcagccgtcgcttcatg**
**tgactccacggagtaccgggcgccgtccaggcacctcgattagttctcgagcttttggagtacgtcgtctttaggtt**
**ggggggaggggttttatgcgatggagtttccccacactgagtgggtggagactgaagttaggccagcttggcacttg**
**atgtaattctccttggaatttgccctttttgagtttggatcttggttcattctcaagcctcagacagtggttcaaag**
**tttttttcttccatttcaggtgtcgtgaaactctagcgtttccgctctagaactagtcca**GCTAGCGCTACCGGTC
GCCACC**atggtgagcaagggcgaggagctgttcaccggggtggtgcccatcctggtcgagctggacggcgacgtaaa**
**cggccacaagttcagcgtgtccggcgagggcgagggcgatgccacctacggcaagctgaccctgaagttcatctgca**
**ccaccggcaagctgcccgtgccctggcccaccctcgtgaccaccctgacctacggcgtgcagtgcttcagccgctac**
**cccgaccacatgaagcagcacgacttcttcaagtccgccatgcccgaaggctacgtccaggagcgcaccatcttctt**
**caaggacgacggcaactacaagacccgcgccgaggtgaagttcgagggcgacaccctggtgaaccgcatcgagctga**
**agggcatcgacttcaaggaggacggcaacatcctggggcacaagctggagtacaactacaacagccacaacgtctat**

atcatggccgacaagcagaagaacggcatcaaggtgaacttcaagatccgccacaacatcgaggacggcagcgtgca
gctcgccgaccactaccagcagaacacccccatcggcgacggccccgtgctgctgcccgacaaccactacctgagca
cccagtccgccctgagcaaagaccccaacgagaagcgcgatcacatggtcctgctggagttcgtgaccgccgccggg
atcactctcggcatggacgagctgtacaagGTcactCcgtgTgaaaacctgtacttccaaggtatggtgagcaaggg
cgaggagctgttcaccggggtggtgcccatcctggtcgagctggacggcgacgtaaacggccacaagttcagcgtgt
ccggcgagggcgagggcgatgccacctacggcaagctgaccctgaagttcatctgcaccaccggcaagctgcccgtg
ccctggcccaccctcgtgaccaccctgacctacggcgtgcagtgcttcagccgctaccccgaccacatgaagcagca
cgacttcttcaagtccgccatgcccgaaggctacgtccaggagcgcaccatcttcttcaaggacgacggcaactaca
agacccgcgccgaggtgaagttcgagggcgacaccctggtgaaccgcatcgagctgaagggcatcgacttcaaggag
gacggcaacatcctggggcacaagctggagtacaactacaacagccacaacgtctatatcatggccgacaagcagaa
gaacggcatcaaggtgaacttcaagatccgccacaacatcgaggacggcagcgtgcagctcgccgaccactaccagc
agaacacccccatcggcgacggccccgtgctgctgcccgacaaccactacctgagcacccagtccgccctgagcaaa
gaccccaacgagaagcgcgatcacatggtcctgctggagttcgtgaccgccgccgggatcactctcggcatggacga
gctgtacaagACcactGcgtgCgaaaacctgtacttccaaggtatggtgagcaagggcgaggagctgttcaccgggg
tggtgcccatcctggtcgagctggacggcgacgtaaacggccacaagttcagcgtgtccggcgagggcgagggcgat
gccacctacggcaagctgaccctgaagttcatctgcaccaccggcaagctgcccgtgccctggcccaccctcgtgac
caccctgacctacggcgtgcagtgcttcagccgctaccccgaccacatgaagcagcacgacttcttcaagtccgcca
tgcccgaaggctacgtccaggagcgcaccatcttcttcaaggacgacggcaactacaagacccgcgccgaggtgaag
ttcgagggcgacaccctggtgaaccgcatcgagctgaagggcatcgacttcaaggaggacggcaacatcctggggca
caagctggagtacaactacaacagccacaacgtctatatcatggccgacaagcagaagaacggcatcaaggtgaact
tcaagatccgccacaacatcgaggacggcagcgtgcagctcgccgaccactaccagcagaacacccccatcggcgac
ggccccgtgctgctgcccgacaaccactacctgagcacccagtccgccctgagcaaagaccccaacgagaagcgcga
tcacatggtcctgctggagttcgtgaccgccgccgggatcactctcggcatggacgagctgtacaagGTcactAcgt
gTgaaaacctgtacttccaaggtatggtgagcaagggcgaggagctgttcaccggggtggtgcccatcctggtcgag
ctggacggcgacgtaaacggccacaagttcagcgtgtccggcgagggcgagggcgatgccacctacggcaagctgac
cctgaagttcatctgcaccaccggcaagctgcccgtgccctggcccaccctcgtgaccaccctgacctacggcgtgc
agtgcttcagccgctaccccgaccacatgaagcagcacgacttcttcaagtccgccatgcccgaaggctacgtccag
gagcgcaccatcttcttcaaggacgacggcaactacaagacccgcgccgaggtgaagttcgagggcgacaccctggt
gaaccgcatcgagctgaagggcatcgacttcaaggaggacggcaacatcctggggcacaagctggagtacaactaca
acagccacaacgtctatatcatggccgacaagcagaagaacggcatcaaggtgaacttcaagatccgccacaacatc
gaggacggcagcgtgcagctcgccgaccactaccagcagaacacccccatcggcgacggccccgtgctgctgcccga
caaccactacctgagcacccagtccgccctgagcaaagaccccaacgagaagcgcgatcacatggtcctgctggagt
tcgtgaccgccgccgggatcactctcggcatggacgagctgtacaagACcactTcgtgCgaaaacctgtacttccaa
ggtatggtgagcaagggcgaggagctgttcaccggggtggtgcccatcctggtcgagctggacggcgacgtaaacgg
ccacaagttcagcgtgtccggcgagggcgagggcgatgccacctacggcaagctgaccctgaagttcatctgcacca
ccggcaagctgcccgtgccctggcccaccctcgtgaccaccctgacctacggcgtgcagtgcttcagccgctacccc
gaccacatgaagcagcacgacttcttcaagtccgccatgcccgaaggctacgtccaggagcgcaccatcttcttcaa
ggacgacggcaactacaagacccgcgccgaggtgaagttcgagggcgacaccctggtgaaccgcatcgagctgaagg
gcatcgacttcaaggaggacggcaacatcctggggcacaagctggagtacaactacaacagccacaacgtctatatc
atggccgacaagcagaagaacggcatcaaggtgaacttcaagatccgccacaacatcgaggacggcagcgtgcagct
cgccgaccactaccagcagaacacccccatcggcgacggccccgtgctgctgcccgacaaccactacctgagcaccc
agtccgccctgagcaaagaccccaacgagaagcgcgatcacatggtcctgctggagttcgtgaccgccgccgggatc
actctcggcatggacgagctgtacaaggaaaacctgtacttccaaAgtggagaAAGCTTgtttaaggaccacgtga
ttacaacccgatatcgagcaccatttgtcatttgacgaatgaatctgatgggcacacaacatcgttgtatggtattg
gatttggtcccttcatcattacaaacaagcacttgtttagaagaaataatggaacactgttggtccaatcactacat
ggtgtattcaaggtcaagaacaccacgactttgcaacaacacctcattgatgggagggacatgataattattcgcat
gcctaaggatttcccaccatttcctcaaaagctgaaatttagagagccacaaagggaagagcgcatatgtcttgtga
caaccaacttccaaactaagagcatgtctagcatggtgtcagacactagttgcacattcccttcatctgatggcata
ttctggaagcattggattcaaaccaaggatgggcagtgtggcagtccattagtatcaactagagatgggttcattgt
tggtatacactcagcatcgaatttcaccaacacaaacaattatttcacaagcgtgccgaaaaacttcatggaattgt
tgacaaatcaggaggcgcagcagtgggttagtggttggcgattaaatgctgactcagtattgtgggggggccataaa
gttttcatgagcaaacctgaagagccttttcagccagttaaggaagcgactcaactcatgaatgaattggtgtactc
gcaagaaaacctgtacttccaaagtATGCATAGATCACGAGATATCAGCCATGGCTTCCCGCCGGCGGTGGCGGCGC
AGGATGATGGCACGCTGCCCATGTCTTGTGCCCAGGAGAGCGGGATGGACCGTCACCCTGCAGCCTGTGCTTCTGCT
AGGATCAATGTGGAGCTCGTGCACCTGACTCCTGAGGAGAAGTCTGCCGTTACTGCCCTGTGGGGCAAGGTGAACGT
GGATGAAGTTGGTGGTGAGGCCCTGGGCAGGCTGCTGGTGGTCTACCCTTGGACCCAGAGGTTCTTTGAGTCCTTTG
GGGATCTGTCCACTCCTGATGCTGTTATGGGCAACCCTAAGGTGAAGGCTCATGGCAAGAAAGTGCTCGGTGCCTTT

AGTGATGGCCTGGCTCACCTGGACAACCTCAAGGGCACCTTTGCCACACTGAGTGAGCTGCACTGTGACAAGCTGCA
CGTGGATCCTGAGAACTTCAGGGTGAGTCTATGGGACCCTTGATGTTTTCTTTCCCCTTCTTTTCTATGGTTAAGTT
CATGTCATAGGAAGGGGATAAGTAACAGGGTACAGTTTAGAATGGGAAACAGACGAATGATTGCATCAGTGTGGAAG
TCTCAGGATCGTTTTAGTTTCTTTTATTTGCTGTTCATAACAATTGTTTTCTTTTGTTTAATTCTTGCTTTCTTTTT
TTTTCTTCTCCGCAATTTTTACTATTATACTTAATGCCTTAACATTGTGTATAACAAAAGGAAATATCTCTGAGATA
CATTAAGTAACTTAAAAAAAAAAACTTTACACAGTCTGCCTAGTACATTACTATTTGGAATATATGTGTGCTTATTTG
CATATTCATAATCTCCCTACTTTATTTTCTTTTATTTTTAATTGATACATAATCATTATACATATTTATGGGTTAAA
GTGTAATGTTTTAATATGTGTACACATATTGACCAAATCAGGGTAATTTTGCATTTGTAATTTTAAAAAATGCTTTC
TTCTTTTAATATACTTTTTTGTTTATCTTATTTCTAATACTTTCCCTAATCTCTTTCTTTCAGGGCAATAATGATAC
AATGTATCATGCCTCTTTGCACCATTCTAAAGAATAACAGTGATAATTTCTGGGTTAAGGCAATAGCAATATTTCTG
CATATAAATATTTCTGCATATAAATTGTAACTGATGTAAGAGGTTTCATATTGCTAATAGCAGCTACAATCCAGCTA
CCATTCTGCTTTTATTTTATGGTTGGGATAAGGCTGGATTATTCTGAGT**CCAAGCTAGGCCCTTTTGCTAATCATG**T
TCATACCTCTTATCTTCCTCCCACAGCTCCTGGGCAACGTGCTGGTCTGTGTGCTGGCCCATCACTTTGGCAAAGAA
TTCACCCCACCAGTGCAGGCTGCCTATCAGAAAGTGGTGGCTGGTGTGGCTAATGCCCTGGCCCACAAGTATCAC**TA**
**A**GC**GGCCGC**TCGCTTTCTTGCTGTCCAATTTCTATTAAAGGTTCCTTTGTTCCCTAAGTCCAACTACTAAACTGGGG
GATATTATGAAGGGCCTCGAGtagggtcatgaaggtttttcttttcctgagaaaacaacacgtattgttttctcagg
ttttgctttttggcctttttctagcttaaaaaaaaaaaaagcaaaactacgaccaccaacccactcctggtttccag
gacggggttcaaatccctgcggcgtctttgctttgactCTCGAGGCT**GATC**A***GCCTCGACTGTGCCTTCTAG***TTGCC
AGCCATCTGTTGTTTGCCCCTCCCCCGTGCCTTCCTTGACCCTGGAAGGTGCCACTCCCACTGTCCTTTCCTAATAA
AATGAGGAAATTGCATCGCATTGTCTGAGTAGGTGTCATTCTATTCTGGGGGGTGGGGTGGGGCAGGACAGCAAGGG
GGAGGATTGGGAAGACAATAGCAGGCATGCTGGGGATGCGGTGGGCTCTATGGCTTCTGAGGCGGAAAGAACCAGCT
GGGGCTCTAGGGGGTATCCCCACGCGCCCTGTAGCGGCGCATTAAGCGCGGCGGGTGTGGTGGTTACGCGCAGCGTG
ACCGCTACACTTGCCAGCGCCCTAGCGCCCGCTCCTTTCGCTTTCTTCCCTTCCTTTCTCGCCACGTTCGCCGGCTT
TCCCCGTCAAGCTCTAAATCGGGGGCTCCCTTTAGGGTTCCGATTTAGTGCTTTACGGCACCTCGACCCCAAAAAAC
TTGATTAGGGTGATGGTTCACGTAGGTAC**G**ctagcaccgttggtttccgtagtgtagtggttatcacgttcgcctaa
cacgcgaaaggtccccggttcgaaaccgggcactacaaaccaacaacgctGGTACCCCTA*GAAGTTCCTATTCCGAA*
*GTTCCTATTC****TCTAGAAAGTATAGGAACTTC***CTTGGCCAAAAAGCCTGAAagatcacgagatatcagccatggcttc
ccgccggcggtggcggcgcaggatgatggcacgctgcccaCgtcttgtgcccaggagagcgggaCggaccgtcaccc
tgcagcctgtgcttctgctaggatcaatgtggaaaacctgtacttccaaGgtaccgagtacaagcccacggtgcgcc
tcgccacccgcgacgacgtccccagggccgtCcgcaccctcgccgccgcgttcgccgactaccccgccacgcgccac
accgtcgatccggaccgccacatcgagcg*ggtcaccgagctgcaagaactcttcctcacgcgcgtcgggctcgacat*
*cggcaaggtgtgggtcgcggacgacggcgccgcggtggcggtctggaccacgccggagagcgtcgaagcgggggcgg*
*tgttcgccgagatcggcccgcgcatggccgagttgagcggttcccggctggccgcgcagcaacagatggaaggcctc*
*ctggcgccgcaccggcccaaggagcccgcgtggttcctggccaccgtcggcgtctcgcccgaccaccagggcaaggg*
*tctgggcagcgccgtcgtgctccccggagtggaggcggccgagcgcgccggggtgcccgccttcctggagacctccg*
*cgccccgcaacctccccttctacg*agcggctcggcttcaccgtcaccgccgacgtcgaggtgcccgaaggaccgcg**c**
**acctggtg**catgacccgcaagcccggtgcc**tga**ggtaccgctcgctgatcagcctcgactgtgccttctagttgcca
gccatctgttgtttgcccctcccccgtgccttccttgaccctggaaggtgccactcccactgtcctttcctaataaa
atgaggaaattgcatcgcattgtctgagtaggtgtcattctattctggggggtggggtggggcaggacagcaagggg
gaggattgggaagacaatagcaggcatgctggggaGTCGACaggtggcacttttcggggaaatgtgcgcggaacccc
tatttgtttatttttctaaatacattcaaatatgtatccgctcatgagacaataaccctgataaatgcttcaataat
attgaaaaaggaagagtatgagtattcaacatttccgtgtcgcccttattcccttttttgcggcattttgccttcct
gtttttgctcacccagaaacgctggtgaaagtaaaagatgctgaagatcagttgggtgcacgagtgggttacatcga
actggatctcaacagcggtaagatccttgagagttttcgccccgaagaacgttttccaatgatgagcacttttaaag
ttctgctatgtggcgcggtattatcccgtattgacgccgggcaagagcaactcggtcgccgcatacactattctcag
aatgacttggttgagtactcaccagtcacagaaaagcatcttacggatggcatgacagtaagagaattatgcagtgc
tgccataaccatgagtgataacactgcggccaacttacttctgacaacgatcggaggaccgaaggagctaaccgctt
ttttgcacaacatgggggatcatgtaactcgccttgatcgttgggaaccggagctgaatgaagccataccaaacgac
gagcgtgacaccacgatgcctgtagcaatggcaacaacgttgcgcaaactattaactggcgaactacttactctagc
ttcccggcaacaattaatagactggatggaggcggataaagttgcaggaccacttctgcgctcg**gcccttccggc**tg
gctggtttattgctgataaatctggagccggtgagcgtgggtctcgcggtatcattgcagcactggggccagatggt
aagccctcccgtatcgtagttatctacacgacggggagtcaggcaactatggatgaacgaaatagacagatcgctga
gataggtgcctcactgattaagcattggtaactgtcagaccaagtttactcatatatactttagattgatttaaa

**G) pAVA3169 [expressing full-length MALAT1 driven by its own promoter with stability-compromised ENE(C8351G)]**

**SV40-polyA (reverse complement): 2126-2384**
**Hygromycin$^R$ (lacks promoter and the 1st Methionine, reverse complement): 2431-3441**
**FRT (reverse complement): 3461-3508**
**XhoI restriction site: 3513-3518**
**10,843 bp genomic locus of MALAT1: 3519-14361**
**Components of the MALAT1 genomic locus:**
**MALAT1 endogenous upstream sequence (promoter): 3520-4698**
**MALAT1 RNA: 4698-14361**
**11 nt qPCR insert: 13040-13050**
**MALAT1-ENE(C8351G)-mascRNA): 13085-13255**
**Position of the ENE mutation G8531G: 13182**
**RNase P cleavage site: between nucleotides 13186 and 13187**
**MALAT1 endogenous downstream sequence: 13256-14361**
**BglII restriction site: 14362-14367**

ATTCAAATATGTATCCGCTCATGAGACAATAACCCTGATAAATGCTTCAATAATATTGAAAAAGGAAGAGTATGAGT
ATTCAACATTTCCGTGTCGCCCTTATTCCCTTTTTTGCGGCATTTTGCCTTCCTGTTTTTGCTCACCCAGAAACGCT
GGTGAAAGTAAAAGATGCTGAAGATCAGTTGGGTGCACGAGTGGGTTACATCGAACTGGATCTCAACAGCGGTAAGA
TCCTTGAGAGTTTTCGCCCCGAAGAACGTTTTCCAATGATGAGCACTTTTAAAGTTCTGCTATGTGGCGCGGTATTA
TCCCGTATTGACGCCGGGCAAGAGCAACTCGGTCGCCGCATACACTATTCTCAGAATGACTTGGTTGAGTACTCACC
AGTCACAGAAAAGCATCTTACGGATGGCATGACAGTAAGAGAATTATGCAGTGCTGCCATAACCATGAGTGATAACA
CTGCGGCCAACTTACTTCTGACAACGATCGGAGGACCGAAGGAGCTAACCGCTTTTTTGCACAACATGGGGGATCAT
GTAACTCGCCTTGATCGTTGGGAACCGGAGCTGAATGAAGCCATACCAAACGACGAGCGTGACACCACGATGCCTGT
AGCAATGGCAACAACGTTGCGCAAACTATTAACTGGCGAACTACTTACTCTAGCTTCCCGGCAACAATTAATAGACT
GGATGGAGGCGGATAAAGTTGCAGGACCACTTCTGCGCTCGGCCCTTCCGGCTGGCTGGTTTATTGCTGATAAATCT
GGAGCCGGTGAGCGTGGGTCTCGCGGTATCATTGCAGCACTGGGGCCAGATGGTAAGCCCTCCCGTATCGTAGTTAT
CTACACGACGGGGAGTCAGGCAACTATGGATGAACGAAATAGACAGATCGCTGAGATAGGTGCCTCACTGATTAAGC
ATTGGTAACTGTCAGACCAAGTTTACTCATATATACTTTAGATTGATTTAAAACTTCATTTTTAATTTAAAAGGATC
TAGGTGAAGATCCTTTTTGATAATCTCATGACCAAAATCCCTTAACGTGAGTTTTCGTTCCACTGAGCGTCAGACCC
CGTAGAAAAGATCAAAGGATCTTCTTGAGATCCTTTTTTTCTGCGCGTAATCTGCTGCTTGCAAACAAAAAAACCAC
CGCTACCAGCGGTGGTTTGTTTGCCGGATCAAGAGCTACCAACTCTTTTTCCGAAGGTAACTGGCTTCAGCAGAGCG
CAGATACCAAATACTGTCCTTCTAGTGTAGCCGTAGTTAGGCCACCACTTCAAGAACTCTGTAGCACCGCCTACATA
CCTCGCTCTGCTAATCCTGTTACCAGTGGCTGCTGCCAGTGGCGATAAGTCGTGTCTTACCGGGTTGGACTCAAGAC
GATAGTTACCGGATAAGGCGCAGCGGTCGGGCTGAACGGGGGGTTCGTGCACACAGCCCAGCTTGGAGCGAACGACC
TACACCGAACTGAGATACCTACAGCGTGAGCATTGAGAAAGCGCCACGCTTCCCGAAGGGAGAAAGGCGGACAGGTA
TCCGGTAAGCGGCAGGGTCGGAACAGGAGAGCGCACGAGGGAGCTTCCAGGGGGAAACGCCTGGTATCTTTATAGTC
CTGTCGGGTTTCGCCACCTCTGACTTGAGCGTCGATTTTTGTGATGCTCGTCAGGGGGGCGGAGCCTATGGAAAAAC
GCCAGCAACGCGGCCTTTTTACGGTTCCTGGCCTTTTGCTGGCCTTTTGCTCACATGTTCTTTCCTGCGTTATCCCC
TGATTCTGTGGATAACCGTATTACCGCCTTTGAGTGAGCTGATACCGCTCGCCGCAGCCGAACGACCGAGCGCAGCG
AGTCAGTGAGCGAGGAAGCGGAAGAGCGCCCAATACGCAAACCGCCTCTCCCCGCGCGTTGGCCGATTCATTAATGC
AGCTGGCACGACAGGTTTCCCGACTGGAAAGCGGGCAGTGAGCGCAACGCAATTAATGTGAGTTAGCTCACTCATTA
GGCACCCCAGGCTTTACACTTTATGCTTCCGGCTCGTATGTTGTGTGGAATTGTGAGCGGATAACAATTTCACACAG
GAAACAGCTATGACCATGATTACGCCAAGCTCTAGCTAGAGGTCGACGGTATACAGACATGATAAGATACATTGATG
AGTTTGGACAAACCACAACTA**GAATGC**AGTGAAAAAAATGCTTTATTTGTGAAATTTGTGATGCTATTGCTTTATTT
GTAACCATTATAAGCTGCAATAAACAAGTTGGGGTGGGCGAAGAACTCCAGCATGAGATCCCCGCGCTGGAGGATCA
TCCAGCCGGCGTCCCGGAAAACGATTCCGAAGCCCAACCTTTCATAGAAGGCGGCGGTGGAATCGAAATCTCGTGAT
GGCAGGTTGGGCGTCGCTTGGTCGGTCATTTCGAAAGTACGTG**CTA**TTCCTTTGCCCTCGGACGAGTGCTGGGGCGT
CGGTTTCCACTATCGGCGAGTACTTCTACACAGCCATCGGTCCAGACGGCCGCGCTTCTGCGGGCGATTTGTGTACG
CCCGACAGTCCCGGCTCCGGATCGGACGATTGCGTCGCATCGACCCTGCGCCCAAGCTGCATCATCGAAATTGCCGT
CAACCAAGCTCTGATAGAGTTGGTCAAGACCAATGCGGAGCATATACGCCCGGAGCCGCGGCGATCCTGCAAGCTCC
GGATGCCTCCGCTCGAAGTAGCGCGTCTGCTGCTCCATACAAGCCAACCACGGCCTCCAGAAGAAGATGTTGGCGAC

CTCGTATTGGGAATCCCCGAACATCGCCTCGCTCCAGTCAATGACCGCTGTTATGCGGCCATTGTCCGTCAGGACAT
TGTTGGAGCCGAAATCCGCGTGCACGAGGTGCCGGACTTCGGGGCAGTCCTCGGCCCAAAGCATCAGCTCATCGAGA
GCCTGCGCGACGGACGCACTGACGGTGTCGTCCATCACAGTTTGCCAGTGATACACATGGGGATCAGCAATCGCGCA
TATGAAATCACGCCATGTAGTGTATTGACCGATTCCTTGCGGTCCGAATGGGCCGAACCCGCTCGTCTGGCTAAGAT
CGGCCGCAGCGATCGCATCCATGGCCTCCGCGACCGGCTGCAGAACAGCGGGCAGTTCGGTTTCAGGCAGGTCTTGC
AACGTGACACCCTGTGCACGGCGGGAGATGCAATAGGTCAGGCTCTCGCTGAATTCCCCAATGTCAAGCACTTCCGG
AATCGGGAGCGCGGCCGATGCAAAGTGCCGATAAACATAACGATCTTTGTAGAAACCATCGGCGCAGCTATTTACCC
GCAGGACATATCCACGCCCTCCTACATCGAAGCTGAAAGCACGAGATTCTTCGCCCTCCGAGAGCTGCATCAGGTCG
GAGACGCTGTCGAACTTTTCGATCAGAAACTTCTCGACAGACGTCGCGGTGAGTTCAGGCTTTTTGGCCAAG*GAAGT*
*TCCTATACTTTCTAGAGAATAGGAACTTCGGAATAGGAACTTC*TAGGCTCGAGGATTCAAGAACAGCTTCTGCTATT
AATTAGCTGTGTGTCATTTCAGGCAAATCACAAAATCTCCAGGCCCCCAAATCCCCATTTGCGAAACGAGGTGACCT
GCCTCAGTTTTCCTTCATCCAGCGATAAAATTCTACTAAAAAATTTCCAACCCCAGAGCAAGCATCAGGACTGTGCC
TGAAGGGATCCGGCTGCATCTCAGTAATATTCCACTCTTACAGGCTTAAAAATAAAAAAGAAAAGAAAAGAAAACAA
GAACAGTTTCACCAGCGTCAATTGAGAACAGCTGCTTCAACAGGCCCTGCTTTATGTGGGCAGGTGGGGACAGGGAG
TTGGCCCAGAAAACAGGACCCTCATTTCCTGGAGCCCCCAGGGCAGCTCCCCAACACCGTACACAACCTGCATCATT
TACAGGAGCCAAAGGAGTTTTAGAAATCAACTTCATAGAGTTGCTGTATCATTGTGTTAGTTTTGCCATCCTAACCT
ATACAGCGTCACTAATCTCTCCCCTCGGAGTTGACTGCCTAAAAACAAGCCATGGATACAGGTTCCAAAGACCCGGG
GGTGAGGGAGCGGAGAGGGTGGGTGTCACCACCCCGTCCAGCTCCAAGCTTTGTGTGCCCTGGAACTCTCCATTTTA
GGTCATTGCTTCAGTTTCTTTTCTAAAAAATTAGGCTGCTGCAAGGTCAGCCTGAGACCACTTCTGCCCCGAGAATT
CTAGACTAGTAAGACCTGGTGACATACAAACGACGAAGATATTTTACAATAATGCCATGGCCCTTGATAGCTACACG
AGGTTTGTGTTCTGATTTTAAATTAATGGATGACGTCGAGACTATATGGAGGAAATGACAAAGGACAGGAGAGAGGT
GGGAAAGGAAGACCTAGACTGAAAATGGAAGTTGGGCAGCAGCTCCACGAAAGAAAGACCAGCCCCCAAGTGCAGTG
ACAGCGCAGAGTAGCGACCGAGAAGTTCCCAGGCCAGCTGCCACCCCGCCCCCATGCCATTCCCCAGAACAGGCACA
GGCGTTAGGGCGGGGCGCGCGTGCGCAGTCACGCGCTGCGCCAACCGCCACAGCTCCGGGAAGGCGGCCAGGACCGG
CTAGAGCCGGTTAGAACCAGTGGCGCCCGCCCACGAGCCAGCGCCTCACAAAGGGAGGGCGGCTCACGGCCCTCGCG
TATCCCTGCGCGGCGCTCGCGAGCCGCCCCTCCCCCGGCGTTTGTCCCTGACGCAGCCCCACCGGTTGCGCAGTCCC
TCCCCGCCCCCGCTCTCCCCTCCGCAGCCTGCAGCCCGAGACTTCTGTAAAGGACTGGGGCCCCGCAACTGGCCTCT
CCTGCCCTCTTAAGCGCAGCGCCATTTTAGCAACGCAGAAGCCCGGCGCCGGGAAGCCTCAGCTCGCCTGAAGGCAG
GTCCCCTCTGACGCCTCCGGGAGCCCAGGTTTCCCAGAGTCCTTGGGACGCAGCGACGAGTTGTGCTGCTATCTTAG
CTGTCCTTATAGGCTGGCCATTCCAGGTGGTGGTATTTAGATAAAACCACTCAAACTCTGCAGTTTGGTCTTGGGGT
TTGGAGGAAAGCTTTTATTTTTCTTCCTGCTCCGGTTCAGAAGGTCTGAAGCTCATACCTAACCAGGCATAACACAG
AATCTGCAAAACAAAAACCCCTAAAAAAGCAGACCCAGAGCAGTGTAAACACTTCTGGGTGTGTCCCTGACTGGCTG
CCCAAGGTCTCTGTGTCTTCGGAGACAAAGCCATTCGCTTAGTTGGTCTACTTTAAAAGGCCACTTGAACTCGCTTT
CCATGGCGATTTGCCTTGTGAGCACTTTCAGGAGAGCCTGGAAGCTGAAAAACGGTAGAAAAATTTCCGTGCGGGCC
GTGGGGGGCTGGCGGCAACTGGGGGGCCGCAGATCAGAGTGGGCCACTGGCAGCCAACGGCCCCCGGGGCTCAGGCG
GGGAGCAGCTCTGTGGTGTGGGATTGAGGCGTTTTCCAAGAGTGGGTTTTCACGTTTCTAAGATTTCCCAAGCAGAC
AGCCCGTGCTGCTCCGATTTCTCGAACAAAAAAGCAAAACGTGTGGCTGTCTTGGGAGCAAGTCGCAGGACTGCAAG
CAGTTGGGGGAGAAAGTCCGCCATTTTGCCACTTCTCAACCGTCCCTGCAAGGCTGGGGCTCAGTTGCGTAATGGAA
AGTAAAGCCCTGAACTATCACACTTTAATCTTCCTTCAAAAGGTGGTAAACTATACCTACTGTCCCTCAAGAGAACA
CAAGAAGTGCTTTAAGAGGTATTTTAAAAGTTCCGGGGGTTTTGTGAGGTGTTTGATGACCCGTTTAAAATATGATT
TCCATGTTTCTTTTGTCTAAAGTTTGCAGCTCAAATCTTTCCACACGCTAGTAATTTAAGTATTTCTGCATGTGTAG
TTTGCATTCAAGTTCCATAAGCTGTTAAGAAAAATCTAGAAAAGTAAAACTAGAACCTATTTTTAACCGAAGAACTA
CTTTTTGCCTCCCTCACAAAGGCGGCGGAAGGTGATCGAATTCCGGTGATGCGAGTTGTTCTCCGTCTATAAATACG
CCTCGCCCGAGCTGTGCGGTAGGCATTGAGGCAGCCAGCGCAGGGGCTTCTGCTGAGGGGGCAGGCGGAGCTTGAGG
AAACCGCAGATAAGTTTTTTTCTCTTTGAAAGATAGAGATTAATACAACTACTTAAAAAATATAGTCAATAGGTTAC
TAAGATATTGCTTAGCGTTAAGTTTTTAACGTAATTTTAATAGCTTAAGATTTTAAGAGAAAATATGAAGACTTAGA
AGAGTAGCATGAGGAAGGAAAAGATAAAAGGTTTCTAAAACATGACGGAGGTTGAGATGAAGCTTCTTCATGGAGTA
AAAAATGTATTTAAAAGAAAATTGAGAGAAAGGACTACAGAGCCCCGAATTAATACCAATAGAAGGGCAATGCTTTT
AGATTAAAATGAAGGTGACTTAAACAGCTTAAAGTTTAGTTTAAAAGTTGTAGGTGATTAAAATAATTTGAAGGCGA
TCTTTTAAAAAGAGATTAAACCGAAGGTGATTAAAAGACCTTGAAATCCATGACGCAGGGAGAATTGCGTCATTTAA
AGCCTAGTTAACGCATTTACTAAACGCAGACGAAAATGGAAAGATTAATTGGGAGTGGTAGGATGAAACAATTTGGA
GAAGATAGAAGTTTGAAGTGGAAAACTGGAAGACAGAAGTACGGGAAGGCGAAGAAAAGAATAGAGAAGATAGGGAA
ATTAGAAGATAAAAACATACTTTTAGAAGAAAAAAGATAAATTTAAACCTGAAAAGTAGGAAGCAGAAGAAAAAAGA
CAAGCTAGGAAACAAAAAGCTAAGGGCAAAATGTACAAACTTAGAAGAAAATTGGAAGATAGAAACAAGATAGAAAA
TGAAAATATTGTCAAGAGTTTCAGATAGAAAATGAAAAACAAGCTAAGACAAGTATTGGAGAAGTATAGAAGATAGA
AAAATATAAAGCCAAAAATTGGATAAAATAGCACTGAAAAAATGAGGAAATTATTGGTAACCAATTTATTTTAAAAG
CCCATCAATTTAATTTCTGGTGGTGCAGAAGTTAGAAGGTAAAGCTTGAGAAGATGAGGGTGTTTACGTAGACCAGA

ACCAATTTAGAAGAATACTTGAAGCTAGAAGGGGAAGTTGGTTAAAAATCACATCAAAAAGCTACTAAAAGGACTGG
TGTAATTTAAAAAAAACTAAGGCAGAAGGCTTTTGGAAGAGTTAGAAGAATTTGGAAGGCCTTAAATATAGTAGCTT
AGTTTGAAAAATGTGAAGGACTTTCGTAACGGAAGTAATTCAAGATCAAGAGTAATTACCAACTTAATGTTTTTGCA
TTGGACTTTGAGTTAAGATTATTTTTTAAATCCTGAGGACTAGCATTAATTGACAGCTGACCCAGGTGCTACACAGA
AGTGGATTCAGTGAATCTAGGAAGACAGCAGCAGACAGGATTCCAGGAACCAGTGTTTGATGAAGCTAGGACTGAGG
AGCAAGCGAGCAAGCAGCAGTTCGTGGTGAAGATAGGAAAAGAGTCCAGGAGCCAGTGCGATTTGGTGAAGGAAGCT
AGGAAGAAGGAAGGAGCGCTAACGATTTGGTGGTGAAGCTAGGAAAAAGGATTCCAGGAAGGAGCGAGTGCAATTTG
GTGATGAAGGTAGCAGGCGGCTTGGCTTGGCAACCACACGGAGGAGGCGAGCAGGCGTTGTGCGTAGAGGATCCTAG
ACCAGCATGCCAGTGTGCCAAGGCCACAGGGAAAGCGAGTGGTTGGTAAAAATCCGTGAGGTCGGCAATATGTTGTT
TTTCTGGAACTTACTTATGGTAACCTTTTATTTATTTTCTAATATAATGGGGGAGTTTCGTACTGAGGTGTAAAGGG
ATTTATATGGGGACGTAGGCCGATTTCCGGGTGTTGTAGGTTTCTCTTTTTCAGGCTTATACTCATGAATCTTGTCT
GAAGCTTTTGAGGGCAGACTGCCAAGTCCTGGAGAAATAGTAGATGGCAAGTTTGTGGGTTTTTTTTTTTTACACGA
ATTTGAGGAAAACCAAATGAATTTGATAGCCAAATTGAGACAATTTCAGCAAATCTGTAAGCAGTTTGTATGTTTAG
TTGGGGTAATGAAGTATTTCAGTTTTGTGAATAGATGACCTGTTTTTACTTCCTCACCCTGAATTCGTTTTGTAAAT
GTAGAGTTTGGATGTGTAACTGAGGCGGGGGGGAGTTTTCAGTATTTTTTTTTGTGGGGGTGGGGGCAAAATATGTT
TTCAGTTCTTTTTCCCTTAGGTCTGTCTAGAATCCTAAAGGCAAATGACTCAAGGTGTAACAGAAAACAAGAAAATC
CAATATCAGGATAATCAGACCACCACAGGTTTACAGTTTATAGAAACTAGAGCAGTTCTCACGTTGAGGTCTGTGGA
AGAGATGTCCATTGGAGAAATGGCTGGTAGTTACTCTTTTTTCCCCCCACCCCCTTAATCAGACTTTAAAAGTGCTT
AACCCCTTAAACTTGTTATTTTTTACTTGAAGCATTTTGGGATGGTCTTAACAGGGAAGAGAGAGGGTGGGGGAGAA
AATGTTTTTTTCTAAGATTTTCCACAGATGCTATAGTACTATTGACAAACTGGGTTAGAGAAGGAGTGTACCGCTGT
GCTGTTGGCACGAACACCTTCAGGGACTGGAGCTGCTTTTATCCTTGGAAGAGTATTCCCAGTTGAAGCTGAAAAGT
ACAGCACAGTGCAGCTTTGGTTCATATTCAGTCATCTCAGGAGAACTTCAGAAGAGCTTGAGTAGGCCAAATGTTGA
AGTTAAGTTTTCCAATAATGTGACTTCTTAAAAGTTTTATTAAAGGGGAGGGGCAAATATTGGCAATTAGTTGGCAG
TGGCCTGTTACGGTTGGGATTGGTGGGGTGGGTTTAGGTAATTGTTTAGTTTATGATTGCAGATAAACTCATGCCAG
AGAACTTAAAGTCTTAGAATGGAAAAAGTAAAGAAATATCAACTTCCAAGTTGGCAAGTAACTCCCAATGATTTAGT
TTTTTTCCCCCCAGTTTGAATTGGGAAGCTGGGGGAAGTTAAATATGAGCCACTGGGTGTACCAGTGCATTAATTTG
GGCAAGGAAAGTGTCATAATTTGATACTGTATCTGTTTTCCTTCAAAGTATAGAGCTTTTGGGGAAGGAAAGTATTG
AACTGGGGGTTGGTCTGGCCTACTGGGCTGACATTAACTACAATTATGGGAAATGCAAAAGTTGTTTGGATATGGTA
GTGTGTGGTTCTCTTTTGGAATTTTTTTCAGGTGATTTAATAATAATTTAAAACTACTATAGAAACTGCAGAGCAAA
GGAAGTGGCTTAATGATCCTGAAGGGATTTCTTCTGATGGTAGCTTTTGTATTATCAAGTAAGATTCTATTTTCAGT
TGTGTGTAAGCAAGTTTTTTTTTAGTGTAGGAGAAATACTTTTCCATTGTTTAACTGCAAAACAAGATGTTAAGGTA
TGCTTCAAAAATTTTGTAAATTGTTTATTTTAAACTTATCTGTTTGTAAATTGTAACTGATTAAGAATTGTGATAGT
TCAGCTTGAATGTCTCTTAGAGGGTGGGCTTTTGTTGATGAGGGAGGGGAAACTTTTTTTTTTTCTATAGACTTTTT
TCAGATAACATCTTCTGAGTCATAACCAGCCTGGCAGTATGATGGCCTAGATGCAGAGAAAACAGCTCCTTGGTGAA
TTGATAAGTAAAGGCAGAAAAGATTATATGTCATACCTCCATTGGGGAATAAGCATAACCCTGAGATTCTTACTACT
GATGAGAACATTATCTGCATATGCCAAAAAATTTTAAGCAAATGAAAGCTACCAATTTAAAGTTACGGAATCTACCA
TTTTAAAGTTAATTGCTTGTCAAGCTATAACCACAAAAATAATGAATTGATGAGAAATACAATGAAGAGGCAATGTC
CATCTCAAAATACTGCTTTTACAAAAGCAGAATAAAAGCGAAAAGAAATGAAAATGTTACACTACATTAATCCTGGA
ATAAAAGAAGCCGAAATAAATGAGAGATGAGTTGGGATCAAGTGGATTGAGGAGGCTGTGCTGTGTGCCAATGTTTC
GTTTGCCTCAGACAGGTATCTCTTCGTTATCAGAAGAGTTGCTTCATTTCATCTGGGAGCAGAAAACAGCAGGCAGC
TGTTAACAGATAAGTTTAACTTGCATCTGCAGTATTGCATGTTAGGGATAAGTGCTTATTTTTAAGAGCTGTGGAGT
TCTTAAATATCAACCATGGCACTTTCTCCTGACCCCTTCCCTAGGGGATTTCAGGATTGAGAAATTTTTCCATCGAG
CCTTTTTAAAATTGTAGGACTTGTTCCTGTGGGCTTCAGTGATGGGATAGTACACTTCACTCAGAGGCATTTGCATC
TTTAAATAATTTCTTAAAAGCCTCTAAAGTGATCAGTGCCTTGATGCCAACTAAGGAAATTTGTTTAGCATTGAATC
TCTGAAGGCTCTATGAAAGGAATAGCATGATGTGCTGTTAGAATCAGATGTTACTGCTAAAATTTACATGTTGTGAT
GTAAATTGTGTAGAAAACCATTAAATCATTCAAAATAATAAACTATTTTTATTAGAGAATGTATACTTTTAGAAAGC
TGTCTCCTTATTTAAATAAAATAGTGTTTGTCTGTAGTTCAGTGTTGGGGCAATCTTGGGGGGGATTCTTCTCTAAT
CTTTCAGAAACTTTGTCTGCGAACACTCTTTAATGGACCAGATCAGGATTTGAGCGGAAGAACGAATGTAACTTTAA
GGCAGGAAAGACAAATTTTATTCTTCATAAAGTGATGAGCATATAATAATTCCAGGCACATGGCAATAGAGGCCCTC
TAAATAAGGAATAAATAACCTCTTAGACAGGTGGGAGATTATGATCAGAGTAAAAGGTAATTACACATTTTATTTCC
AGAAAGTCAGGGGTCTATAAATTGACAGTGATTAGAGTAATACTTTTTCACATTTCCAAAGTTTGCATGTTAACTTT
AAATGCTTACAATCTTAGAGTGGTAGGCAATGTTTTACACTATTGACCTTATATAGGGAAGGGAGGGGGTGCCTGTG
GGGTTTTAAAGAATTTTCCTTTGCAGAGGCATTTCATCCTTCATGAAGCCATTCAGGATTTTGAATTGCATATGAGT
GCTTGGCTCTTCCTTCTGTTCTAGTGAGTGTATGAGACCTTGCAGTGAGTTTATCAGCATACTCAAAATTTTTTTCC
TGGAATTTGGAGGGATGGGAGGAGGGGGTGGGGCTTACTTGTTGTAGCTTTTTTTTTTTTTACAGACTTCACAGAGA
ATGCAGTTGTCTTGACTTCAGGTCTGTCTGTTCTGTTGGCAAGTAAATGCAGTACTGTTCTGATCCCGCTGCTATTA
GAATGCATTGTGAAACGACTGGAGTATGATTAAAAGTTGTGTTCCCCAATGCTTGGAGTAGTGATTGTTGAAGGAAA

AAATCCAGCTGAGTGATAAAGGCTGAGTGTTGAGGAAATTTCTGCAGTTTTAAGCAGTCGTATTTGTGATTGAAGCT
GAGTACATTTTGCTGGTGTATTTTTAGGTAAAATGCTTTTTGTTCATTTCTGGTGGTGGGAGGGGACTGAAGCCTTT
AGTCTTTTCCAGATGCAACCTTAAAATCAGTGACAAGAAACATTCCAAACAAGCAACAGTCTTCAAGAAATTAAACT
GGCAAGTGGAAATGTTTAAACAGTTCAGTGATCTTTAGTGCATTGTTTATGTGTGGGTTTCTCTCTCCCCTCCCTTG
GTCTTAATTCTTACATGCAGGAACACTCAGCAGACACACGTATGCGAAGGGCCAGAGAAGCCAGACCCAGTAAGAAA
AAATAGCCTATTTACTTTAAATAAACCAAACATTCCATTTTAAATGTGGGGATTGGGAACCACTAGTTCTTTCAGAT
GGTATTCTTCAGACTATAGAAGGAGCTTCCAGTTGAATTCACCAGTGGACAAAATGAGGAAAACAGGTGAACAAGCT
TTTTCTGTATTTACATACAAAGTCAGATCAGTTATGGGACAATAGTATTGAATAGATTTCAGCTTTATGCTGGAGTA
ACTGGCATGTGAGCAAACTGTGTTGGCGTGGGGGTGGAGGGGTGAGGTGGGCGCTAAGCCTTTTTTTAAGATTTTTC
AGGTACCCCTCACTAAAGGCACCGAAGGCTTAAAGTAGGACAACCATGGAGCCTTCCTGTGGCAGGAGAGACAACAA
AGCGCTATTATCCTAAGGTCAAGAGAAGTGTCAGCCTCACCTGATTTTTATTAGTAATGAGGACTTGCCTCAACTCC
CTCTTTCTGGAGTGAAGCATCCGAAGGAATGCTTGAAGTACCCCTGGGCTTCTCTTAACATTTAAGCAAGCTGTTTT
TATAGCAGCTCTTAATAATAAAGCCCAAATCTCAAGCGGTGCTTGAAGGGGAGGGAAAGGGGGAAAGCGGGCAACCA
CTTTTCCCTAGCTTTTCCAGAAGCCTGTTAAAAGCAAGGTCTCCCCACAAGCAACTTCTCTGCCACATCGCCACCCC
GTGCCTTTTGATCTAGCACAGACCCTTCACCCCTCACCTCGATGCAGCCAGTAGCTTGGATCCTTGTGGGCATGATC
CATAATCGGTTTCAAGGTAACGATGGTGTCGAGGTCTTTGGTGGGTTGAACTATGTTAGAAAAGGCCATTAATTTGC
CTGCAAATTGTTAACAGAAGGGTATTAAAACCACAGCTAAGTAGCTCTATTATAATACTTATCCAGTGACTAAAACC
AACTTAAACCAGTAAGTGGAGAAATAACATGTTCAAGAACTGTAATGCTGGGTGGGAACATGTAACTTGTAGACTGG
AGAAGATAGGCATTTGAGTGGCTGAGAGGGCTTTTGGGTGGGAATGCAAAAATTCTCTGCTAAGACTTTTTCAGGTG
AACATAACAGACTTGGCCAAGCTAGC**cgctcgacgta**ATCTTAGCGGAAGCTGATCTCCAATGCTCTTCAG**TAGGGT**
**CATGAAGGTTTTTCTTTTCCTGAGAAAACAACACGTATTGTTTTCTCAGGTTTTGCTTTTTGGCCTTTTTCTAGCTT**
**AAAAAAAAAAAAAGGAAAA**GATGCTGGTGGTTGGCACTCCTGGTTTCCAGGACGGGGTTCAAATCCCTGCGGCGTCT
TTGCTTTGACTACTAATCTGTCTTCAGGACTCTTTCTGTATTTCTCCTTTTCTCTGCAGGTGCTAGTTCTTGGAGTT
TTGGGGAGGTGGGAGGTAACAGCACAATATCTTTGAACTATATACATCCTTGATGTATAATTTGTCAGGAGCTTGAC
TTGATTGTATATTCATATTTACACGAGAACCTAATATAACTGCCTTGTCTTTTTCAGGTAATAGCCTGCAGCTGGTG
TTTTGAGAAGCCCTACTGCTGAAAACTTAACAATTTTGTGTAATAAAAATGGAGAAGCTCTAAATTGTTGTGGTTCT
TTTGTGAATAAAAAAATCTTGATTGGGGAAAAAAGATGGGTGTTCTGTGGGCTTGTTCTGTTAAATCTGTGGTCTAT
AAACACAGCACCCATAATTACAGCATAATCTTCAAGTAGGGTACGGACTTTGGGGGATTGGTGCGAGGGTAGTGGGT
GAGTGGCCTACTAAAAAGCCCAGTAACCCCCACAGGAAAATAGGGAACTTCTTTTTAAGTAGCCTCCTTTCCACTAT
TTAGTAATTGGCTGTGAGCTGGGCTGGGGGAGAAATGGGGCGGGGTGTGTGTGTCATTGGAAAGCTCTCTTTTTTGT
TTTTTTGAGACAGTCTCACTTTGTCCCCCAGGCTGGAGTGTAGTGGCATGATCTCTGCAAACTGCAACCTCCACTTC
TGGGGTCCAAGTGGTTGTCCTGCTTCACCCTCCCTGTAGCTGGGACTACAGGTGCACACCACCACGCCTGGCTAATT
TTTGTATTTTCAGTTAGAGACGTGGTTTTACCATATTGGCCAGGCTGGTCTCAAACTCCTGACCTCGTGTGATCCAC
CCGCCTGGGCCTCTGAAAGTGCTGGGATTACAGGTGTGAGCCACCAAGCCTGGCCGATCCTTTTAAGTTTTTAAACC
AGTTAAGCTCTTTGGTTCCCCCTCAGAGTCCCAAGGTCCTGGGTCACTCAGGATCACATTTTCCTTTAATCAGTTGT
CACTGGTCCCCTTTTGTCCCTTTGAACGTGCTGTGGGATTAGTAGCAGCATCTGGCTGTTGGAAGGACTGGCTGGGA
TCTCAGGTGAAATACCCTCCCTGGCCCTCCTTA**CCCTTAAGATCTCCC**GATCCGTCGACGTCAGGTGGCACTTTTCG
GGGAAATGTGCGCGGAACCCCTATTTGTTTATTTTTCTAAATAC

**H) pAVA3171 [expressing full-length endogenous MALAT1 driven by its own promoter]**

**SV40-polyA (reverse complement): 2126-2384**
**Hygromycin$^{R}$ (lacks promoter and the 1st Methionine, reverse complement): 2431-3441**
**FRT (reverse complement): 3461-3508**
**XhoI restriction site: 3513-3518**
**10,843 bp genomic locus of MALAT1: 3519-14361**
**Components of the MALAT1 genomic locus:**
**MALAT1 endogenous upstream sequence (promoter): 3520-4698**
**MALAT1 RNA: 4698-14361**
**11 nt qPCR insert: 13040-13050**
**MALAT1 (ENE(WT)-mascRNA): 13085-13255**
**RNase P cleavage site: between nucleotides 13186 and 13187**
**MALAT1 endogenous downstream sequence: 13256-14361**
**BglII restriction site: 14362-14367**

ATTCAAATATGTATCCGCTCATGAGACAATAACCCTGATAAATGCTTCAATAATATTGAAAAAGGAAGAGTATGAGT
ATTCAACATTTCCGTGTCGCCCTTATTCCCTTTTTTGCGGCATTTTGCCTTCCTGTTTTTGCTCACCCAGAAACGCT
GGTGAAAGTAAAAGATGCTGAAGATCAGTTGGGTGCACGAGTGGGTTACATCGAACTGGATCTCAACAGCGGTAAGA
TCCTTGAGAGTTTTCGCCCCGAAGAACGTTTTCCAATGATGAGCACTTTTAAAGTTCTGCTATGTGGCGCGGTATTA
TCCCGTATTGACGCCGGGCAAGAGCAACTCGGTCGCCGCATACACTATTCTCAGAATGACTTGGTTGAGTACTCACC
AGTCACAGAAAAGCATCTTACGGATGGCATGACAGTAAGAGAATTATGCAGTGCTGCCATAACCATGAGTGATAACA
CTGCGGCCAACTTACTTCTGACAACGATCGGAGGACCGAAGGAGCTAACCGCTTTTTTGCACAACATGGGGGATCAT
GTAACTCGCCTTGATCGTTGGGAACCGGAGCTGAATGAAGCCATACCAAACGACGAGCGTGACACCACGATGCCTGT
AGCAATGGCAACAACGTTGCGCAAACTATTAACTGGCGAACTACTTACTCTAGCTTCCCGGCAACAATTAATAGACT
GGATGGAGGCGGATAAAGTTGCAGGACCACTTCTGCGCTCGGCCCTTCCGGCTGGCTGGTTTATTGCTGATAAATCT
GGAGCCGGTGAGCGTGGGTCTCGCGGTATCATTGCAGCACTGGGGCCAGATGGTAAGCCCTCCCGTATCGTAGTTAT
CTACACGACGGGGAGTCAGGCAACTATGGATGAACGAAATAGACAGATCGCTGAGATAGGTGCCTCACTGATTAAGC
ATTGGTAACTGTCAGACCAAGTTTACTCATATATACTTTAGATTGATTTAAAACTTCATTTTTAATTTAAAAGGATC
TAGGTGAAGATCCTTTTTGATAATCTCATGACCAAAATCCCTTAACGTGAGTTTTCGTTCCACTGAGCGTCAGACCC
CGTAGAAAAGATCAAAGGATCTTCTTGAGATCCTTTTTTTCTGCGCGTAATCTGCTGCTTGCAAACAAAAAAACCAC
CGCTACCAGCGGTGGTTTGTTTGCCGGATCAAGAGCTACCAACTCTTTTTCCGAAGGTAACTGGCTTCAGCAGAGCG
CAGATACCAAATACTGTCCTTCTAGTGTAGCCGTAGTTAGGCCACCACTTCAAGAACTCTGTAGCACCGCCTACATA
CCTCGCTCTGCTAATCCTGTTACCAGTGGCTGCTGCCAGTGGCGATAAGTCGTGTCTTACCGGGTTGGACTCAAGAC
GATAGTTACCGGATAAGGCGCAGCGGTCGGGCTGAACGGGGGGTTCGTGCACACAGCCCAGCTTGGAGCGAACGACC
TACACCGAACTGAGATACCTACAGCGTGAGCATTGAGAAAGCGCCACGCTTCCCGAAGGGAGAAAGGCGGACAGGTA
TCCGGTAAGCGGCAGGGTCGGAACAGGAGAGCGCACGAGGGAGCTTCCAGGGGGAAACGCCTGGTATCTTTATAGTC
CTGTCGGGTTTCGCCACCTCTGACTTGAGCGTCGATTTTTGTGATGCTCGTCAGGGGGGCGGAGCCTATGGAAAAAC
GCCAGCAACGCGGCCTTTTTACGGTTCCTGGCCTTTTGCTGGCCTTTTGCTCACATGTTCTTTCCTGCGTTATCCCC
TGATTCTGTGGATAACCGTATTACCGCCTTTGAGTGAGCTGATACCGCTCGCCGCAGCCGAACGACCGAGCGCAGCG
AGTCAGTGAGCGAGGAAGCGGAAGAGCGCCCAATACGCAAACCGCCTCTCCCCGCGCGTTGGCCGATTCATTAATGC
AGCTGGCACGACAGGTTTCCCGACTGGAAAGCGGGCAGTGAGCGCAACGCAATTAATGTGAGTTAGCTCACTCATTA
GGCACCCCAGGCTTTACACTTTATGCTTCCGGCTCGTATGTTGTGTGGAATTGTGAGCGGATAACAATTTCACACAG
GAAACAGCTATGACCATGATTACGCCAAGCTCTAGCTAGAGGTCGACGGTATACAGACATGATAAGATACATTGATG
AGTTTGGACAAACCACAACTA**GAATGC**AGTGAAAAAAATGCTTTATTTGTGAAATTTGTGATGCTATTGCTTTATTT
GTAACCATTATAAGCTGCAATAAACAAGTTGGGGTGGGCGAAGAACTCCAGCATGAGATCCCCGCGCTGGAGGATCA
TCCAGCCGGCGTCCCGGAAAACGATTCCGAAGCCCAACCTTTCATAGAAGGCGGCGGTGGAATCGAAATCTCGTGAT
GGCAGGTTGGGCGTCGCTTGGTCGGTCATTTCGAAAGTACGTG**CTA**TTCCTTTGCCCTCGGACGAGTGCTGGGGCGT
CGGTTTCCACTATCGGCGAGTACTTCTACACAGCCATCGGTCCAGACGGCCGCGCTTCTGCGGGCGATTTGTGTACG
CCCGACAGTCCCGGCTCCGGATCGGACGATTGCGTCGCATCGACCCTGCGCCCAAGCTGCATCATCGAAATTGCCGT
CAACCAAGCTCTGATAGAGTTGGTCAAGACCAATGCGGAGCATATACGCCCGGAGCCGCGGCGATCCTGCAAGCTCC
GGATGCCTCCGCTCGAAGTAGCGCGTCTGCTGCTCCATACAAGCCAACCACGGCCTCCAGAAGAAGATGTTGGCGAC
CTCGTATTGGGAATCCCCGAACATCGCCTCGCTCCAGTCAATGACCGCTGTTATGCGGCCATTGTCCGTCAGGACAT
TGTTGGAGCCGAAATCCGCGTGCACGAGGTGCCGGACTTCGGGGCAGTCCTCGGCCCAAAGCATCAGCTCATCGAGA
GCCTGCGCGACGGACGCACTGACGGTGTCGTCCATCACAGTTTGCCAGTGATACACATGGGGATCAGCAATCGCGCA

TATGAAATCACGCCATGTAGTGTATTGACCGATTCCTTGCGGTCCGAATGGGCCGAACCCGCTCGTCTGGCTAAGAT
CGGCCGCAGCGATCGCATCCATGGCCTCCGCGACCGGCTGCAGAACAGCGGGCAGTTCGGTTTCAGGCAGGTCTTGC
AACGTGACACCCTGTGCACGGCGGGAGATGCAATAGGTCAGGCTCTCGCTGAATTCCCCAATGTCAAGCACTTCCGG
AATCGGGAGCGCGGCCGATGCAAAGTGCCGATAAACATAACGATCTTTGTAGAAACCATCGGCGCAGCTATTTACCC
GCAGGACATATCCACGCCCTCCTACATCGAAGCTGAAAGCACGAGATTCTTCGCCCTCCGAGAGCTGCATCAGGTCG
GAGACGCTGTCGAACTTTTCGATCAGAAACTTCTCGACAGACGTCGCGGTGAGTTCAGGCTTTTTGGCCAAGGAAGT
TCCTATACTTTCTAGAGAATAGGAACTTCGGAATAGGAACTTCTAGGCTCGAGGATTCAAGAACAGCTTCTGCTATT
AATTAGCTGTGTGTCATTTCAGGCAAATCACAAAATCTCCAGGCCCCCAAATCCCCATTTGCGAAACGAGGTGACCT
GCCTCAGTTTTCCTTCATCCAGCGATAAAATTCTACTAAAAAATTTCCAACCCCAGAGCAAGCATCAGGACTGTGCC
TGAAGGGATCCGGCTGCATCTCAGTAATATTCCACTCTTACAGGCTTAAAAATAAAAAAGAAAAGAAAAGAAAACAA
GAACAGTTTCACCAGCGTCAATTGAGAACAGCTGCTTCAACAGGCCCTGCTTTATGTGGGCAGGTGGGGACAGGGAG
TTGGCCCAGAAAACAGGACCCTCATTTCCTGGAGCCCCCAGGGCAGCTCCCCAACACCGTACACAACCTGCATCATT
TACAGGAGCCAAAGGAGTTTTAGAAATCAACTTCATAGAGTTGCTGTATCATTGTGTTAGTTTTGCCATCCTAACCT
ATACAGCGTCACTAATCTCTCCCCTCGGAGTTGACTGCCTAAAAACAAGCCATGGATACAGGTTCCAAAGACCCGGG
GGTGAGGGAGCGGAGAGGGTGGGTGTCACCACCCCGTCCAGCTCCAAGCTTTGTGTGCCCTGGAACTCTCCATTTTA
GGTCATTGCTTCAGTTTCTTTTCTAAAAAATTAGGCTGCTGCAAGGTCAGCCTGAGACCACTTCTGCCCCGAGAATT
CTAGACTAGTAAGACCTGGTGACATACAAACGACGAAGATATTTTACAATAATGCCATGGCCCTTGATAGCTACACG
AGGTTTGTGTTCTGATTTTAAATTAATGGATGACGTCGAGACTATATGGAGGAAATGACAAAGGACAGGAGAGAGGT
GGGAAAGGAAGACCTAGACTGAAAATGGAAGTTGGGCAGCAGCTCCACGAAAGAAAGACCAGCCCCCAAGTGCAGTG
ACAGCGCAGAGTAGCGACCGAGAAGTTCCCAGGCCAGCTGCCACCCCGCCCCCATGCCATTCCCCAGAACAGGCACA
GGCGTTAGGGCGGGGCGCGCGTGCGCAGTCACGCGCTGCGCCAACCGCCACAGCTCCGGGAAGGCGGCCAGGACCGG
CTAGAGCCGGTTAGAACCAGTGGCGCCCGCCCACGAGCCAGCGCCTCACAAAGGGAGGGCGGCTCACGGCCCTCGCG
TATCCCTGCGCGGCGCTCGCGAGCCGCCCCTCCCCCGGCGTTTGTCCCTGACGCAGCCCCACCGGTTGCGCAGTCCC
TCCCCGCCCCCGCTCTCCCCTCCGCAGCCTGCAGCCCGAGACTTCTGTAAAGGACTGGGGCCCCGCAACTGGCCTCT
CCTGCCCTCTTAAGCGCAGCGCCATTTTAGCAACGCAGAAGCCCGGCGCCGGGAAGCCTCAGCTCGCCTGAAGGCAG
GTCCCCTCTGACGCCTCCGGGAGCCCAGGTTTCCCAGAGTCCTTGGGACGCAGCGACGAGTTGTGCTGCTATCTTAG
CTGTCCTTATAGGCTGGCCATTCCAGGTGGTGGTATTTAGATAAAACCACTCAAACTCTGCAGTTTGGTCTTGGGGT
TTGGAGGAAAGCTTTTATTTTTCTTCCTGCTCCGGTTCAGAAGGTCTGAAGCTCATACCTAACCAGGCATAACACAG
AATCTGCAAAACAAAAACCCCTAAAAAAGCAGACCCAGAGCAGTGTAAACACTTCTGGGTGTGTCCCTGACTGGCTG
CCCAAGGTCTCTGTGTCTTCGGAGACAAAGCCATTCGCTTAGTTGGTCTACTTTAAAAGGCCACTTGAACTCGCTTT
CCATGGCGATTTGCCTTGTGAGCACTTTCAGGAGAGCCTGGAAGCTGAAAAACGGTAGAAAAATTTCCGTGCGGGCC
GTGGGGGGCTGGCGGCAACTGGGGGGCCGCAGATCAGAGTGGGCCACTGGCAGCCAACGGCCCCCGGGGCTCAGGCG
GGGAGCAGCTCTGTGGTGTGGGATTGAGGCGTTTTCCAAGAGTGGGTTTTCACGTTTCTAAGATTTCCCAAGCAGAC
AGCCCGTGCTGCTCCGATTTCTCGAACAAAAAAGCAAAACGTGTGGCTGTCTTGGGAGCAAGTCGCAGGACTGCAAG
CAGTTGGGGGAGAAAGTCCGCCATTTTGCCACTTCTCAACCGTCCCTGCAAGGCTGGGGCTCAGTTGCGTAATGGAA
AGTAAAGCCCTGAACTATCACACTTTAATCTTCCTTCAAAAGGTGGTAAACTATACCTACTGTCCCTCAAGAGAACA
CAAGAAGTGCTTTAAGAGGTATTTTAAAAGTTCCGGGGGTTTTGTGAGGTGTTTGATGACCCGTTTAAAATATGATT
TCCATGTTTCTTTTGTCTAAAGTTTGCAGCTCAAATCTTTCCACACGCTAGTAATTTAAGTATTTCTGCATGTGTAG
TTTGCATTCAAGTTCCATAAGCTGTTAAGAAAAATCTAGAAAAGTAAAACTAGAACCTATTTTTAACCGAAGAACTA
CTTTTTGCCTCCCTCACAAAGGCGGCGGAAGGTGATCGAATTCCGGTGATGCGAGTTGTTCTCCGTCTATAAATACG
CCTCGCCCGAGCTGTGCGGTAGGCATTGAGGCAGCCAGCGCAGGGGCTTCTGCTGAGGGGGCAGGCGGAGCTTGAGG
AAACCGCAGATAAGTTTTTTTCTCTTTGAAAGATAGAGATTAATACAACTACTTAAAAAATATAGTCAATAGGTTAC
TAAGATATTGCTTAGCGTTAAGTTTTTAACGTAATTTTAATAGCTTAAGATTTTAAGAGAAAATATGAAGACTTAGA
AGAGTAGCATGAGGAAGGAAAAGATAAAAGGTTTCTAAAACATGACGGAGGTTGAGATGAAGCTTCTTCATGGAGTA
AAAAATGTATTTAAAAGAAAATTGAGAGAAAGGACTACAGAGCCCCGAATTAATACCAATAGAAGGGCAATGCTTTT
AGATTAAAATGAAGGTGACTTAAACAGCTTAAAGTTTAGTTTAAAAGTTGTAGGTGATTAAAATAATTTGAAGGCGA
TCTTTTAAAAAGAGATTAAACCGAAGGTGATTAAAAGACCTTGAAATCCATGACGCAGGGAGAATTGCGTCATTTAA
AGCCTAGTTAACGCATTTACTAAACGCAGACGAAAATGGAAAGATTAATTGGGAGTGGTAGGATGAAACAATTTGGA
GAAGATAGAAGTTTGAAGTGGAAAACTGGAAGACAGAAGTACGGGAAGGCGAAGAAAAGAATAGAGAAGATAGGGAA
ATTAGAAGATAAAAACATACTTTTAGAAGAAAAAAGATAAATTTAAACCTGAAAAGTAGGAAGCAGAAGAAAAAAGA
CAAGCTAGGAAACAAAAAGCTAAGGGCAAAATGTACAAACTTAGAAGAAAATTGGAAGATAGAAACAAGATAGAAAA
TGAAAATATTGTCAAGAGTTTCAGATAGAAAATGAAAAACAAGCTAAGACAAGTATTGGAGAAGTATAGAAGATAGA
AAAATATAAAGCCAAAAATTGGATAAAATAGCACTGAAAAAATGAGGAAATTATTGGTAACCAATTTATTTTAAAAG
CCCATCAATTTAATTTCTGGTGGTGCAGAAGTTAGAAGGTAAAGCTTGAGAAGATGAGGGTGTTTACGTAGACCAGA
ACCAATTTAGAAGAATACTTGAAGCTAGAAGGGGAAGTTGGTTAAAAATCACATCAAAAAGCTACTAAAAGGACTGG
TGTAATTTAAAAAAAACTAAGGCAGAAGGCTTTTGGAAGAGTTAGAAGAATTTGGAAGGCCTTAAATATAGTAGCTT
AGTTTGAAAAATGTGAAGGACTTTCGTAACGGAAGTAATTCAAGATCAAGAGTAATTACCAACTTAATGTTTTTGCA

TTGGACTTTGAGTTAAGATTATTTTTTAAATCCTGAGGACTAGCATTAATTGACAGCTGACCCAGGTGCTACACAGA
AGTGGATTCAGTGAATCTAGGAAGACAGCAGCAGACAGGATTCCAGGAACCAGTGTTTGATGAAGCTAGGACTGAGG
AGCAAGCGAGCAAGCAGCAGTTCGTGGTGAAGATAGGAAAAGAGTCCAGGAGCCAGTGCGATTTGGTGAAGGAAGCT
AGGAAGAAGGAAGGAGCGCTAACGATTTGGTGGTGAAGCTAGGAAAAAGGATTCCAGGAAGGAGCGAGTGCAATTTG
GTGATGAAGGTAGCAGGCGGCTTGGCTTGGCAACCACACGGAGGAGGCGAGCAGGCGTTGTGCGTAGAGGATCCTAG
ACCAGCATGCCAGTGTGCCAAGGCCACAGGGAAAGCGAGTGGTTGGTAAAAATCCGTGAGGTCGGCAATATGTTGTT
TTTCTGGAACTTACTTATGGTAACCTTTTATTTATTTTCTAATATAATGGGGGAGTTTCGTACTGAGGTGTAAAGGG
ATTTATATGGGGACGTAGGCCGATTTCCGGGTGTTGTAGGTTTCTCTTTTTCAGGCTTATACTCATGAATCTTGTCT
GAAGCTTTTGAGGGCAGACTGCCAAGTCCTGGAGAAATAGTAGATGGCAAGTTTGTGGGTTTTTTTTTTTTACACGA
ATTTGAGGAAAACCAAATGAATTTGATAGCCAAATTGAGACAATTTCAGCAAATCTGTAAGCAGTTTGTATGTTTAG
TTGGGGTAATGAAGTATTTCAGTTTTGTGAATAGATGACCTGTTTTTACTTCCTCACCCTGAATTCGTTTTGTAAAT
GTAGAGTTTGGATGTGTAACTGAGGCGGGGGGGAGTTTTCAGTATTTTTTTTTGTGGGGGTGGGGGCAAAATATGTT
TTCAGTTCTTTTTCCCTTAGGTCTGTCTAGAATCCTAAAGGCAAATGACTCAAGGTGTAACAGAAAACAAGAAAATC
CAATATCAGGATAATCAGACCACCACAGGTTTACAGTTTATAGAAACTAGAGCAGTTCTCACGTTGAGGTCTGTGGA
AGAGATGTCCATTGGAGAAATGGCTGGTAGTTACTCTTTTTTCCCCCCACCCCCTTAATCAGACTTTAAAAGTGCTT
AACCCCTTAAACTTGTTATTTTTTACTTGAAGCATTTTGGGATGGTCTTAACAGGGAAGAGAGAGGGTGGGGGAGAA
AATGTTTTTTTCTAAGATTTTCCACAGATGCTATAGTACTATTGACAAACTGGGTTAGAGAAGGAGTGTACCGCTGT
GCTGTTGGCACGAACACCTTCAGGGACTGGAGCTGCTTTTATCCTTGGAAGAGTATTCCCAGTTGAAGCTGAAAAGT
ACAGCACAGTGCAGCTTTGGTTCATATTCAGTCATCTCAGGAGAACTTCAGAAGAGCTTGAGTAGGCCAAATGTTGA
AGTTAAGTTTTCCAATAATGTGACTTCTTAAAAGTTTTATTAAAGGGGAGGGGCAAATATTGGCAATTAGTTGGCAG
TGGCCTGTTACGGTTGGGATTGGTGGGGTGGGTTTAGGTAATTGTTTAGTTTATGATTGCAGATAAACTCATGCCAG
AGAACTTAAAGTCTTAGAATGGAAAAAGTAAAGAAATATCAACTTCCAAGTTGGCAAGTAACTCCCAATGATTTAGT
TTTTTTCCCCCCAGTTTGAATTGGGAAGCTGGGGGAAGTTAAATATGAGCCACTGGGTGTACCAGTGCATTAATTTG
GGCAAGGAAAGTGTCATAATTTGATACTGTATCTGTTTTCCTTCAAAGTATAGAGCTTTTGGGGAAGGAAAGTATTG
AACTGGGGGTTGGTCTGGCCTACTGGGCTGACATTAACTACAATTATGGGAAATGCAAAAGTTGTTTGGATATGGTA
GTGTGTGGTTCTCTTTTGGAATTTTTTTCAGGTGATTTAATAATAATTTAAAACTACTATAGAAACTGCAGAGCAAA
GGAAGTGGCTTAATGATCCTGAAGGGATTTCTTCTGATGGTAGCTTTTGTATTATCAAGTAAGATTCTATTTTCAGT
TGTGTGTAAGCAAGTTTTTTTTTAGTGTAGGAGAAATACTTTTCCATTGTTTAACTGCAAAACAAGATGTTAAGGTA
TGCTTCAAAAATTTTGTAAATTGTTTATTTTAAACTTATCTGTTTGTAAATTGTAACTGATTAAGAATTGTGATAGT
TCAGCTTGAATGTCTCTTAGAGGGTGGGCTTTTGTTGATGAGGGAGGGGAAACTTTTTTTTTTCTATAGACTTTTT
TCAGATAACATCTTCTGAGTCATAACCAGCCTGGCAGTATGATGGCCTAGATGCAGAGAAAACAGCTCCTTGGTGAA
TTGATAAGTAAAGGCAGAAAAGATTATATGTCATACCTCCATTGGGGAATAAGCATAACCCTGAGATTCTTACTACT
GATGAGAACATTATCTGCATATGCCAAAAAATTTTAAGCAAATGAAAGCTACCAATTTAAAGTTACGGAATCTACCA
TTTTAAAGTTAATTGCTTGTCAAGCTATAACCACAAAAATAATGAATTGATGAGAAATACAATGAAGAGGCAATGTC
CATCTCAAAATACTGCTTTTACAAAAGCAGAATAAAAGCGAAAAGAAATGAAAATGTTACACTACATTAATCCTGGA
ATAAAAGAAGCCGAAATAAATGAGAGATGAGTTGGGATCAAGTGGATTGAGGAGGCTGTGCTGTGTGCCAATGTTTC
GTTTGCCTCAGACAGGTATCTCTTCGTTATCAGAAGAGTTGCTTCATTTCATCTGGGAGCAGAAAACAGCAGGCAGC
TGTTAACAGATAAGTTTAACTTGCATCTGCAGTATTGCATGTTAGGGATAAGTGCTTATTTTTAAGAGCTGTGGAGT
TCTTAAATATCAACCATGGCACTTTCTCCTGACCCCTTCCCTAGGGGATTTCAGGATTGAGAAATTTTTCCATCGAG
CCTTTTTAAAATTGTAGGACTTGTTCCTGTGGGCTTCAGTGATGGGATAGTACACTTCACTCAGAGGCATTTGCATC
TTTAAATAATTTCTTAAAAGCCTCTAAAGTGATCAGTGCCTTGATGCCAACTAAGGAAATTTGTTTAGCATTGAATC
TCTGAAGGCTCTATGAAAGGAATAGCATGATGTGCTGTTAGAATCAGATGTTACTGCTAAAATTTACATGTTGTGAT
GTAAATTGTGTAGAAAACCATTAAATCATTCAAAATAATAAACTATTTTTATTAGAGAATGTATACTTTTAGAAAGC
TGTCTCCTTATTTAAATAAAATAGTGTTTGTCTGTAGTTCAGTGTTGGGGCAATCTTGGGGGGGATTCTTCTCTAAT
CTTTCAGAAACTTTGTCTGCGAACACTCTTTAATGGACCAGATCAGGATTTGAGCGGAAGAACGAATGTAACTTTAA
GGCAGGAAAGACAAATTTTATTCTTCATAAAGTGATGAGCATATAATAATTCCAGGCACATGGCAATAGAGGCCCTC
TAAATAAGGAATAAATAACCTCTTAGACAGGTGGGAGATTATGATCAGAGTAAAAGGTAATTACACATTTTATTTCC
AGAAAGTCAGGGGTCTATAAATTGACAGTGATTAGAGTAATACTTTTTCACATTTCCAAAGTTTGCATGTTAACTTT
AAATGCTTACAATCTTAGAGTGGTAGGCAATGTTTTACACTATTGACCTTATATAGGGAAGGAGGGGGTGCCTGTG
GGGTTTTAAAGAATTTTCCTTTGCAGAGGCATTTCATCCTTCATGAAGCCATTCAGGATTTTGAATTGCATATGAGT
GCTTGGCTCTTCCTTCTGTTCTAGTGAGTGTATGAGACCTTGCAGTGAGTTTATCAGCATACTCAAAATTTTTTTCC
TGGAATTTGGAGGGATGGGAGGAGGGGGTGGGGCTTACTTGTTGTAGCTTTTTTTTTTTTTACAGACTTCACAGAGA
ATGCAGTTGTCTTGACTTCAGGTCTGTCTGTTCTGTTGGCAAGTAAATGCAGTACTGTTCTGATCCCGCTGCTATTA
GAATGCATTGTGAAACGACTGGAGTATGATTAAAAGTTGTGTTCCCCAATGCTTGGAGTAGTGATTGTTGAAGGAAA
AAATCCAGCTGAGTGATAAAGGCTGAGTGTTGAGGAAATTTCTGCAGTTTTAAGCAGTCGTATTTGTGATTGAAGCT
GAGTACATTTTGCTGGTGTATTTTTAGGTAAAATGCTTTTTGTTCATTTCTGGTGGTGGGAGGGGACTGAAGCCTTT
AGTCTTTTCCAGATGCAACCTTAAAATCAGTGACAAGAAACATTCCAAACAAGCAACAGTCTTCAAGAAATTAAACT

GGCAAGTGGAAATGTTTAAACAGTTCAGTGATCTTTAGTGCATTGTTTATGTGTGGGTTTCTCTCTCCCCTCCCTTG
GTCTTAATTCTTACATGCAGGAACACTCAGCAGACACACGTATGCGAAGGGCCAGAGAAGCCAGACCCAGTAAGAAA
AAATAGCCTATTTACTTTAAATAAACCAAACATTCCATTTTAAATGTGGGGATTGGGAACCACTAGTTCTTTCAGAT
GGTATTCTTCAGACTATAGAAGGAGCTTCCAGTTGAATTCACCAGTGGACAAAATGAGGAAAACAGGTGAACAAGCT
TTTTCTGTATTTACATACAAAGTCAGATCAGTTATGGACAATAGTATTGAATAGATTTCAGCTTTATGCTGGAGTA
ACTGGCATGTGAGCAAACTGTGTTGGCGTGGGGGTGGAGGGGTGAGGTGGGCGCTAAGCCTTTTTTTAAGATTTTTC
AGGTACCCCTCACTAAAGGCACCGAAGGCTTAAAGTAGGACAACCATGGAGCCTTCCTGTGGCAGGAGAGACAACAA
AGCGCTATTATCCTAAGGTCAAGAGAAGTGTCAGCCTCACCTGATTTTTATTAGTAATGAGGACTTGCCTCAACTCC
CTCTTTCTGGAGTGAAGCATCCGAAGGAATGCTTGAAGTACCCCTGGGCTTCTCTTAACATTTAAGCAAGCTGTTTT
TATAGCAGCTCTTAATAATAAAGCCCAAATCTCAAGCGGTGCTTGAAGGGGAGGGAAAGGGGGAAAGCGGGCAACCA
CTTTTCCCTAGCTTTTCCAGAAGCCTGTTAAAAGCAAGGTCTCCCCACAAGCAACTTCTCTGCCACATCGCCACCCC
GTGCCTTTTGATCTAGCACAGACCCTTCACCCCTCACCTCGATGCAGCCAGTAGCTTGGATCCTTGTGGGCATGATC
CATAATCGGTTTCAAGGTAACGATGGTGTCGAGGTCTTTGGTGGGTTGAACTATGTTAGAAAAGGCCATTAATTTGC
CTGCAAATTGTTAACAGAAGGGTATTAAAACCACAGCTAAGTAGCTCTATTATAATACTTATCCAGTGACTAAAACC
AACTTAAACCAGTAAGTGGAGAAATAACATGTTCAAGAACTGTAATGCTGGGTGGGAACATGTAACTTGTAGACTGG
AGAAGATAGGCATTTGAGTGGCTGAGAGGGCTTTTGGGTGGGAATGCAAAAATTCTCTGCTAAGACTTTTTCAGGTG
AACATAACAGACTTGGCCAAGCTAGC**cgctcgacgta**ATCTTAGCGGAAGCTGATCTCCAATGCTCTTCAG**TAGGGT**
**CATGAAGGTTTTTCTTTTCCTGAGAAAACAACACGTATTGTTTTCTCAGGTTTTGCTTTTTGGCCTTTTTCTAGCTT**
**AAAAAAAAAAAAAGCAAAA**GATGCTGGTGGTTGGCACTCCTGGTTTCCAGGACGGGGTTCAAATCCCTGCGGCGTCT
TTGCTTTGACTACTAATCTGTCTTCAGGACTCTTTCTGTATTTCTCCTTTTCTCTGCAGGTGCTAGTTCTTGGAGTT
TTGGGGAGGTGGGAGGTAACAGCACAATATCTTTGAACTATATACATCCTTGATGTATAATTTGTCAGGAGCTTGAC
TTGATTGTATATTCATATTTACACGAGAACCTAATATAACTGCCTTGTCTTTTTCAGGTAATAGCCTGCAGCTGGTG
TTTTGAGAAGCCCTACTGCTGAAAACTTAACAATTTTGTGTAATAAAAATGGAGAAGCTCTAAATTGTTGTGGTTCT
TTTGTGAATAAAAAAATCTTGATTGGGGAAAAAAGATGGGTGTTCTGTGGGCTTGTTCTGTTAAATCTGTGGTCTAT
AAACACAGCACCCATAATTACAGCATAATCTTCAAGTAGGGTACGGACTTTGGGGGATTGGTGCGAGGGTAGTGGGT
GAGTGGCCTACTAAAAAGCCCAGTAACCCCCACAGGAAAATAGGGAACTTCTTTTTAAGTAGCCTCCTTTCCACTAT
TTAGTAATTGGCTGTGAGCTGGGCTGGGGGAGAAATGGGGCGGGGTGTGTGTGTCATTGGAAAGCTCTCTTTTTTGT
TTTTTTGAGACAGTCTCACTTTGTCCCCCAGGCTGGAGTGTAGTGGCATGATCTCTGCAAACTGCAACCTCCACTTC
TGGGGTCCAAGTGGTTGTCCTGCTTCACCCTCCCTGTAGCTGGGACTACAGGTGCACACCACCACGCCTGGCTAATT
TTTGTATTTTCAGTTAGAGACGTGGTTTTACCATATTGGCCAGGCTGGTCTCAAACTCCTGACCTCGTGTGATCCAC
CCGCCTGGGCCTCTGAAAGTGCTGGGATTACAGGTGTGAGCCACCAAGCCTGGCCGATCCTTTTAAGTTTTTAAACC
AGTTAAGCTCTTTGGTTCCCCCTCAGAGTCCCAAGGTCCTGGGTCACTCAGGATCACATTTTCCTTTAATCAGTTGT
CACTGGTCCCCTTTTGTCCCTTTGAACGTGCTGTGGGATTAGTAGCAGCATCTGGCTGTTGGAAGGACTGGCTGGGA
TCTCAGGTGAAATACCCTCCCTGGCCCTCCTTA**CCCTTAAGATCTCCC**GATCCGTCGACGTCAGGTGGCACTTTTCG
GGGAAATGTGCGCGGAACCCCTATTTGTTTATTTTTCTAAATAC

**I) pAVA3129 [lentiviral expression of Cas9 and two sgRNAs: U6-sgRNA1, 7SK-sgRNA2]**

**U6 promoter: 1716-1956**
**sgRNA1: 1967-1986**
**sgRNA Scaffold: 1987-2062**
**7SK promoter: 2075- 2316**
**sgRNA2: 2319-2338**
**sgRNA Scaffold: 2339-2414**
**Filler Sequence (HIV-1 isolate): 2462-2579**
**EF1alfa promoter: 2623-2873**
**Cas9: 2912-7111**
**P2A peptide: 7118-7183**
**Blasticidin$^R$: 7184-7582**

```
TTAATGTAGTCTTATGCAATACTCTTGTAGTCTTGCAACATGGTAACGATGAGTTAGCAACATGCCTTACAAGGAGA
GAAAAAGCACCGTGCATGCCGATTGGTGGAAGTAAGGTGGTACGATCGTGCCTTATTAGGAAGGCAACAGACGGGTC
TGACATGGATTGGACGAACCACTGAATTGCCGCATTGCAGAGATATTGTATTTAAGTGCCTAGCTCGATACATAAAC
GGGTCTCTCTGGTTAGACCAGATCTGAGCCTGGGAGCTCTCTGGCTAACTAGGGAACCCACTGCTTAAGCCTCAATA
AAGCTTGCCTTGAGTGCTTCAAGTAGTGTGTGCCCGTCTGTTGTGTGACTCTGGTAACTAGAGATCCCTCAGACCCT
TTTAGTCAGTGTGGAAAATCTCTAGCAGTGGCGCCCGAACAGGGACTTGAAAGCGAAAGGGAAACCAGAGGAGCTCT
CTCGACGCAGGACTCGGCTTGCTGAAGCGCGCACGGCAAGAGGCGAGGGGCGGCGACTGGTGAGTACGCCAAAAATT
TTGACTAGCGGAGGCTAGAAGGAGAGAGATGGGTGCGAGAGCGTCAGTATTAAGCGGGGGAGAATTAGATCGCGATG
GGAAAAAATTCGGTTAAGGCCAGGGGGAAAGAAAAAATATAAATTAAAACATATAGTATGGGCAAGCAGGGAGCTAG
AACGATTCGCAGTTAATCCTGGCCTGTTAGAAACATCAGAAGGCTGTAGACAAATACTGGGACAGCTACAACCATCC
CTTCAGACAGGATCAGAAGAACTTAGATCATTATATAATACAGTAGCAACCCTCTATTGTGTGCATCAAAGGATAGA
GATAAAAGACACCAAGGAAGCTTTAGACAAGATAGAGGAAGAGCAAAACAAAAGTAAGACCACCGCACAGCAAGCGG
CCGCTGATCTTCAGACCTGGAGGAGGAGATATGAGGGACAATTGGAGAAGTGAATTATATAAATATAAAGTAGTAAA
AATTGAACCATTAGGAGTAGCACCCACCAAGGCAAAGAGAAGAGTGGTGCAGAGAGAAAAAAGAGCAGTGGGAATAG
GAGCTTTGTTCCTTGGGTTCTTGGGAGCAGCAGGAAGCACTATGGGCGCAGCGTCAATGACGCTGACGGTACAGGCC
AGACAATTATTGTCTGGTATAGTGCAGCAGCAGAACAATTTGCTGAGGGCTATTGAGGCGCAACAGCATCTGTTGCA
ACTCACAGTCTGGGGCATCAAGCAGCTCCAGGCAAGAATCCTGGCTGTGGAAAGATACCTAAAGGATCAACAGCTCC
TGGGGATTTGGGGTTGCTCTGGAAAACTCATTTGCACCACTGCTGTGCCTTGGAATGCTAGTTGGAGTAATAAATCT
CTGGAACAGATTTGGAATCACACGACCTGGATGGAGTGGGACAGAGAAATTAACAATTACACAAGCTTAATACACTC
CTTAATTGAAGAATCGCAAAACCAGCAAGAAAAGAATGAACAAGAATTATTGGAATTAGATAAATGGGCAAGTTTGT
GGAATTGGTTTAACATAACAAATTGGCTGTGGTATATAAAATTATTCATAATGATAGTAGGAGGCTTGGTAGGTTTA
AGAATAGTTTTTGCTGTACTTTCTATAGTGAATAGAGTTAGGCAGGGATATTCACCATTATCGTTTCAGACCCACCT
CCCAACCCCGAGGGGACCCAGAGAGGGCCTATTTCCCATGATTCCTTCATATTTGCATATACGATACAAGGCTGTTA
GAGAGATAATTAGAATTAATTTGACTGTAAACACAAAGATATTAGTACAAAATACGTGACGTAGAAAGTAATAATTT
CTTGGGTAGTTTGCAGTTTTAAAATTATGTTTTAAAATGGACTATCATATGCTTACCGTAACTTGAAAGTATTTCGA
TTTCTTGGCTTTATATATCTTGTGGAAAGGACGAAACACCGNNNNNNNNNNNNNNNNNNNGTTTTAGAGCTAGAAA
TAGCAAGTTAAAATAAGGCTAGTCCGTTATCAACTTGAAAAAGTGGCACCGAGTCGGTGCTTTTTTGAATTCctgca
gtatttagcatgccccacccatctgcaaggcattctggatagtgtcaaaacagccggaaatcaagtccgtttatctc
aaactttagcattttgggaataaatgatatttgctatgctggttaaattagattttagttaaatttcctgctgaagc
tctagtacgataagcaacttgacctaagtgtaaagttgagacttccttcaggtttatatagcttgtgcgccgcttgg
gtacctCGNNNNNNNNNNNNNNNNNNNGTTTTAGAGCTAGAAATAGCAAGTTAAAATAAGGCTAGTCCGTTATCAA
CTTGAAAAAGTGGCACCGAGTCGGTGCTTTTTTGAATTCTAGATCTTGAGACAAATGGCAGTATTCATCCACAATTT
TAAAAGAAAAGGGGGGATTGGGGGGTACAGTGCAGGGGAAAGAATAGTAGACATAATAGCAACAGACATACAAACTA
AAGAATTACAAAAACAAATTACAAAAATTCAAAATTTTCGGGTTTATTACAGGGACAGCAGAGATCCACTTTGGCGC
CGGCTCGAGTGGCTCCGGTGCCCGTCAGTGGGCAGAGCGCACATCGCCCACAGTCCCCGAGAAGTTGGGGGGAGGGG
TCGGCAATTGAACCGGTGCCTAGAGAAGGTGGCGCGGGGTAAACTGGGAAAGTGATGTCGTGTACTGGCTCCGCCTT
TTTCCCGAGGGTGGGGGAGAACCGTATATAAGTGCAGTAGTCGCCGTGAACGTTCTTTTTCGCAACGGGTTTGCCGC
CAGAACACAGGTGTCGTGACGCGGGATCCGCCACCATGGATTACAAAGACGATGACGATAAGATGGCCCCAAAGAAG
AAGCGGAAGGTCGGTATCCACGGAGTCCCAGCAGCCGACAAGAAGTACAGCATCGGCCTGGACATCGGCACCAACTC
TGTGGGCTGGGCCGTGATCACCGACGAGTACAAGGTGCCCAGCAAGAAATTCAAGGTGCTGGGCAACACCGACCGGC
ACAGCATCAAGAAGAACCTGATCGGAGCCCTGCTGTTCGACAGCGGCGAAACAGCCGAGGCCACCCGGCTGAAGAGA
```

ACCGCCAGAAGAAGATACACCAGACGGAAGAACCGGATCTGCTATCTGCAAGAGATCTTCAGCAACGAGATGGCCAA
GGTGGACGACAGCTTCTTCCACAGACTGGAAGAGTCCTTCCTGGTGGAAGAGGATAAGAAGCACGAGCGGCACCCCA
TCTTCGGCAACATCGTGGACGAGGTGGCCTACCACGAGAAGTACCCCACCATCTACCACCTGAGAAAGAAACTGGTG
GACAGCACCGACAAGGCCGACCTGCGGCTGATCTATCTGGCCCTGGCCCACATGATCAAGTTCCGGGGCCACTTCCT
GATCGAGGGCGACCTGAACCCCGACAACAGCGACGTGGACAAGCTGTTCATCCAGCTGGTGCAGACCTACAACCAGC
TGTTCGAGGAAAACCCCATCAACGCCAGCGGCGTGGACGCCAAGGCCATCCTGTCTGCCAGACTGAGCAAGAGCAGA
CGGCTGGAAAATCTGATCGCCCAGCTGCCCGGCGAGAAGAAGAATGGCCTGTTCGGCAACCTGATTGCCCTGAGCCT
GGGCCTGACCCCCAACTTCAAGAGCAACTTCGACCTGGCCGAGGATGCCAAACTGCAGCTGAGCAAGGACACCTACG
ACGACGACCTGGACAACCTGCTGGCCCAGATCGGCGACCAGTACGCCGACCTGTTTCTGGCCGCCAAGAACCTGTCC
GACGCCATCCTGCTGAGCGACATCCTGAGAGTGAACACCGAGATCACCAAGGCCCCCCTGAGCGCCTCTATGATCAA
GAGATACGACGAGCACCACCAGGACCTGACCCTGCTGAAAGCTCTCGTGCGGCAGCAGCTGCCTGAGAAGTACAAAG
AGATTTTCTTCGACCAGAGCAAGAACGGCTACGCCGGCTACATTGACGGCGGAGCCAGCCAGGAAGAGTTCTACAAG
TTCATCAAGCCCATCCTGGAAAAGATGGACGGCACCGAGGAACTGCTCGTGAAGCTGAACAGAGAGGACCTGCTGCG
GAAGCAGCGGACCTTCGACAACGGCAGCATCCCCCACCAGATCCACCTGGGAGAGCTGCACGCCATTCTGCGGCGGC
AGGAAGATTTTTACCCATTCCTGAAGGACAACCGGGAAAAGATCGAGAAGATCCTGACCTTCCGCATCCCCTACTAC
GTGGGCCCTCTGGCCAGGGGAAACAGCAGATTCGCCTGGATGACCAGAAAGAGCGAGGAAACCATCACCCCCTGGAA
CTTCGAGGAAGTGGTGGACAAGGGCGCTTCCGCCCAGAGCTTCATCGAGCGGATGACCAACTTCGATAAGAACCTGC
CCAACGAGAAGGTGCTGCCCAAGCACAGCCTGCTGTACGAGTACTTCACCGTGTATAACGAGCTGACCAAAGTGAAA
TACGTGACCGAGGGAATGAGAAAGCCCGCCTTCCTGAGCGGCGAGCAGAAAAAGGCCATCGTGGACCTGCTGTTCAA
GACCAACCGGAAAGTGACCGTGAAGCAGCTGAAAGAGGACTACTTCAAGAAAATCGAGTGCTTCGACTCCGTGGAAA
TCTCCGGCGTGGAAGATCGGTTCAACGCCTCCCTGGGCACATACCACGATCTGCTGAAAATTATCAAGGACAAGGAC
TTCCTGGACAATGAGGAAAACGAGGACATTCTGGAAGATATCGTGCTGACCCTGACACTGTTTGAGGACAGAGAGAT
GATCGAGGAACGGCTGAAAACCTATGCCCACCTGTTCGACGACAAAGTGATGAAGCAGCTGAAGCGGCGGAGATACA
CCGGCTGGGGCAGGCTGAGCCGGAAGCTGATCAACGGCATCCGGGACAAGCAGTCCGGCAAGACAATCCTGGATTTC
CTGAAGTCCGACGGCTTCGCCAACAGAAACTTCATGCAGCTGATCCACGACGACAGCCTGACCTTTAAAGAGGACAT
CCAGAAAGCCCAGGTGTCCGGCCAGGGCGATAGCCTGCACGAGCACATTGCCAATCTGGCCGGCAGCCCCGCCATTA
AGAAGGGCATCCTGCAGACAGTGAAGGTGGTGGACGAGCTCGTGAAAGTGATGGGCCGGCACAAGCCCGAGAACATC
GTGATCGAAATGGCCAGAGAGAACCAGACCACCCAGAAGGGACAGAAGAACAGCCGCGAGAGAATGAAGCGGATCGA
AGAGGGCATCAAAGAGCTGGGCAGCCAGATCCTGAAAGAACACCCCGTGGAAAACACCCAGCTGCAGAACGAGAAGC
TGTACCTGTACTACCTGCAGAATGGGCGGGATATGTACGTGGACCAGGAACTGGACATCAACCGGCTGTCCGACTAC
GATGTGGACCATATCGTGCCTCAGAGCTTTCTGAAGGACGACTCCATCGACAACAAGGTGCTGACCAGAAGCGACAA
GAACCGGGGCAAGAGCGACAACGTGCCCTCCGAAGAGGTCGTGAAGAAGATGAAGAACTACTGGCGGCAGCTGCTGA
ACGCCAAGCTGATTACCCAGAGAAAGTTCGACAATCTGACCAAGGCCGAGAGAGGCGGCCTGAGCGAACTGGATAAG
GCCGGCTTCATCAAGAGACAGCTGGTGGAAACCCGGCAGATCACAAAGCACGTGGCACAGATCCTGGACTCCCGGAT
GAACACTAAGTACGACGAGAATGACAAGCTGATCCGGGAAGTGAAAGTGATCACCCTGAAGTCCAAGCTGGTGTCCG
ATTTCCGGAAGGATTTCCAGTTTTACAAAGTGCGCGAGATCAACAACTACCACCACGCCCACGACGCCTACCTGAAC
GCCGTCGTGGGAACCGCCCTGATCAAAAAGTACCCTAAGCTGGAAAGCGAGTTCGTGTACGGCGACTACAAGGTGTA
CGACGTGCGGAAGATGATCGCCAAGAGCGAGCAGGAAATCGGCAAGGCTACCGCCAAGTACTTCTTCTACAGCAACA
TCATGAACTTTTTCAAGACCGAGATTACCCTGGCCAACGGCGAGATCCGGAAGCGGCCTCTGATCGAGACAAACGGC
GAAACCGGGGAGATCGTGTGGGATAAGGGCCGGGATTTTGCCACCGTGCGGAAAGTGCTGAGCATGCCCCAAGTGAA
TATCGTGAAAAAGACCGAGGTGCAGACAGGCGGCTTCAGCAAAGAGTCTATCCTGCCCAAGAGGAACAGCGATAAGC
TGATCGCCAGAAAGAAGGACTGGGACCCTAAGAAGTACGGCGGCTTCGACAGCCCCACCGTGGCCTATTCTGTGCTG
GTGGTGGCCAAAGTGGAAAAGGGCAAGTCCAAGAAACTGAAGAGTGTGAAAGAGCTGCTGGGGATCACCATCATGGA
AAGAAGCAGCTTCGAGAAGAATCCCATCGACTTTCTGGAAGCCAAGGGCTACAAAGAAGTGAAAAAGGACCTGATCA
TCAAGCTGCCTAAGTACTCCCTGTTCGAGCTGGAAAACGGCCGGAAGAGAATGCTGGCCTCTGCCGGCGAACTGCAG
AAGGGAAACGAACTGGCCCTGCCCTCCAAATATGTGAACTTCCTGTACCTGGCCAGCCACTATGAGAAGCTGAAGGG
CTCCCCCGAGGATAATGAGCAGAAACAGCTGTTTGTGGAACAGCACAAGCACTACCTGGACGAGATCATCGAGCAGA
TCAGCGAGTTCTCCAAGAGAGTGATCCTGGCCGACGCTAATCTGGACAAAGTGCTGTCCGCCTACAACAAGCACCGG
GATAAGCCCATCAGAGAGCAGGCCGAGAATATCATCCACCTGTTTACCCTGACCAATCTGGGAGCCCCTGCCGCCTT
CAAGTACTTTGACACCACCATCGACCGGAAGAGGTACACCAGCACCAAAGAGGTGCTGGACGCCACCCTGATCCACC
AGAGCATCACCGGCCTGTACGAGACACGGATCGACCTGTCTCAGCTGGGAGGCGACAAGCGTCCTGCTGCTACTAAG
AAAGCTGGTCAAGCTAAGAAAAAGAAAGCTAGCGGCAGCGGCGCCACCAACTTCAGCCTGCTGAAGCAGGCCGGCGA
CGTGGAGGAGAACCCCGGCCCC**ATG**GCCAAGCCTTTGTCTCAAGAAGAATCCACCCTCATTGAAAGAGCAACGGCTA
CAATCAACAGCATCCCCATCTCTGAAGACTACAGCGTCGCCAGCGCAGCTCTCTCTAGCGACGGCCGCATCTTCACT
GGTGTCAATGTATATCATTTTACTGGGGGACCTTGTGCAGAACTCGTGGTGCTGGGCACTGCTGCTGCTGCGGCAGC
TGGCAACCTGACTTGTATCGTCGCGATCGGAAATGAGAACAGGGGCATCTTGAGCCCCTGCGGACGGTGCCGACAGG
TGCTTCTCGATCTGCATCCTGGGATCAAAGCCATAGTGAAGGACAGTGATGGACAGCCGACGGCAGTTGGGATTCGT

GAATTGCTGCCCTCTGGTTATGTGTGGGAGGGC**TAA**ACGCGTTAAGTCGACAATCAACCTCTGGATTACAAAATTTG
TGAAAGATTGACTGGTATTCTTAACTATGTTGCTCCTTTTACGCTATGTGGATACGCTGCTTTAATGCCTTTGTATC
ATGCTATTGCTTCCCGTATGGCTTTCATTTTCTCCTCCTTGTATAAATCCTGGTTGCTGTCTCTTTATGAGGAGTTG
TGGCCCGTTGTCAGGCAACGTGGCGTGGTGTGCACTGTGTTTGCTGACGCAACCCCCACTGGTTGGGGCATTGCCAC
CACCTGTCAGCTCCTTTCCGGGACTTTCGCTTTCCCCCTCCCTATTGCCACGGCGGAACTCATCGCCGCCTGCCTTG
CCCGCTGCTGGACAGGGGCTCGGCTGTTGGGCACTGACAATTCCGTGGTGTTGTCGGGGAAATCATCGTCCTTTCCT
TGGCTGCTCGCCTGTGTTGCCACCTGGATTCTGCGCGGGACGTCCTTCTGCTACGTCCCTTCGGCCCTCAATCCAGC
GGACCTTCCTTCCCGCGGCCTGCTGCCGGCTCTGCGGCCTCTTCCGCGTCTTCGCCTTCGCCCTCAGACGAGTCGGA
TCTCCCTTTGGGCCGCCTCCCCGCGTCGACTTTAAGACCAATGACTTACAAGGCAGCTGTAGATCTTAGCCACTTTT
TAAAAGAAAAGGGGGGACTGGAAGGGCTAATTCACTCCCAACGAAGACAAGATCTGCTTTTTGCTTGTACTGGGTCT
CTCTGGTTAGACCAGATCTGAGCCTGGGAGCTCTCTGGCTAACTAGGGAACCCACTGCTTAAGCCTCAATAAAGCTT
GCCTTGAGTGCTTCAAGTAGTGTGTGCCCGTCTGTTGTGTGACTCTGGTAACTAGAGATCCCTCAGACCCTTTTAGT
CAGTGTGGAAAATCTCTAGCAGTACGTATAGTAGTTCATGTCATCTTATTATTCAGTATTTATAACTTGCAAAGAAA
TGAATATCAGAGAGTGAGAGGAACTTGTTTATTGCAGCTTATAATGGTTACAAATAAAGCAATAGCATCACAAATTT
CACAAATAAAGCATTTTTTTCACTGCATTCTAGTTGTGGTTTGTCCAAACTCATCAATGTATCTTATCATGTCTGGC
TCTAGCTATCCCGCCCCTAACTCCGCCCATCCCGCCCCTAACTCCGCCCAGTTCCGCCCATTCTCCGCCCCATGGCT
GACTAATTTTTTTTATTTATGCAGAGGCCGAGGCCGCCTCGGCCTCTGAGCTATTCCAGAAGTAGTGAGGAGGCTTT
TTTGGAGGCCTAGGGACGTACCCAATTCGCCCTATAGTGAGTCGTATTACGCGCGCTCACTGGCCGTCGTTTTACAA
CGTCGTGACTGGGAAAACCCTGGCGTTACCCAACTTAATCGCCTTGCAGCACATCCCCCTTTCGCCAGCTGGCGTAA
TAGCGAAGAGGCCCGCACCGATCGCCCTTCCCAACAGTTGCGCAGCCTGAATGGCGAATGGGACGCGCCCTGTAGCG
GCGCATTAAGCGCGGCGGGTGTGGTGGTTACGCGCAGCGTGACCGCTACACTTGCCAGCGCCCTAGCGCCCGCTCCT
TTCGCTTTCTTCCCTTCCTTTCTCGCCACGTTCGCCGGCTTTCCCCGTCAAGCTCTAAATCGGGGGCTCCCTTTAGG
GTTCCGATTTAGTGCTTTACGGCACCTCGACCCCAAAAAACTTGATTAGGGTGATGGTTCACGTAGTGGGCCATCGC
CCTGATAGACGGTTTTTCGCCCTTTGACGTTGGAGTCCACGTTCTTTAATAGTGGACTCTTGTTCCAAACTGGAACA
ACACTCAACCCTATCTCGGTCTATTCTTTTGATTTATAAGGGATTTTGCCGATTTCGGCCTATTGGTTAAAAAATGA
GCTGATTTAACAAAAATTTAACGCGAATTTTAACAAAATATTAACGCTTACAATTTAGGTGGCACTTTTCGGGGAAA
TGTGCGCGGAACCCCTATTTGTTTATTTTTCTAAATACATTCAAATATGTATCCGCTCATGAGACAATAACCCTGAT
AAATGCTTCAATAATATTGAAAAAGGAAGAGTATGAGTATTCAACATTTCCGTGTCGCCCTTATTCCCTTTTTTGCG
GCATTTTGCCTTCCTGTTTTTGCTCACCCAGAAACGCTGGTGAAAGTAAAAGATGCTGAAGATCAGTTGGGTGCACG
AGTGGGTTACATCGAACTGGATCTCAACAGCGGTAAGATCCTTGAGAGTTTTCGCCCCGAAGAACGTTTTCCAATGA
TGAGCACTTTTAAAGTTCTGCTATGTGGCGCGGTATTATCCCGTATTGACGCCGGGCAAGAGCAACTCGGTCGCCGC
ATACACTATTCTCAGAATGACTTGGTTGAGTACTCACCAGTCACAGAAAAGCATCTTACGGATGGCATGACAGTAAG
AGAATTATGCAGTGCTGCCATAACCATGAGTGATAACACTGCGGCCAACTTACTTCTGACAACGATCGGAGGACCGA
AGGAGCTAACCGCTTTTTTGCACAACATGGGGGATCATGTAACTCGCCTTGATCGTTGGGAACCGGAGCTGAATGAA
GCCATACCAAACGACGAGCGTGACACCACGATGCCTGTAGCAATGGCAACAACGTTGCGCAAACTATTAACTGGCGA
ACTACTTACTCTAGCTTCCCGGCAACAATTAATAGACTGGATGGAGGCGGATAAAGTTGCAGGACCACTTCTGCGCT
CGGCCCTTCCGGCTGGCTGGTTTATTGCTGATAAATCTGGAGCCGGTGAGCGTGGGTCTCGCGGTATCATTGCAGCA
CTGGGGCCAGATGGTAAGCCCTCCCGTATCGTAGTTATCTACACGACGGGGAGTCAGGCAACTATGGATGAACGAAA
TAGACAGATCGCTGAGATAGGTGCCTCACTGATTAAGCATTGGTAACTGTCAGACCAAGTTTACTCATATATACTTT
AGATTGATTTAAAACTTCATTTTTAATTTAAAAGGATCTAGGTGAAGATCCTTTTTGATAATCTCATGACCAAAATC
CCTTAACGTGAGTTTTCGTTCCACTGAGCGTCAGACCCCGTAGAAAAGATCAAAGGATCTTCTTGAGATCCTTTTTT
TCTGCGCGTAATCTGCTGCTTGCAAACAAAAAAACCACCGCTACCAGCGGTGGTTTGTTTGCCGGATCAAGAGCTAC
CAACTCTTTTTCCGAAGGTAACTGGCTTCAGCAGAGCGCAGATACCAAATACTGTTCTTCTAGTGTAGCCGTAGTTA
GGCCACCACTTCAAGAACTCTGTAGCACCGCCTACATACCTCGCTCTGCTAATCCTGTTACCAGTGGCTGCTGCCAG
TGGCGATAAGTCGTGTCTTACCGGGTTGGACTCAAGACGATAGTTACCGGATAAGGCGCAGCGGTCGGGCTGAACGG
GGGGTTCGTGCACACAGCCCAGCTTGGAGCGAACGACCTACACCGAACTGAGATACCTACAGCGTGAGCTATGAGAA
AGCGCCACGCTTCCCGAAGGGAGAAAGGCGGACAGGTATCCGGTAAGCGGCAGGGTCGGAACAGGAGAGCGCACGAG
GGAGCTTCCAGGGGGAAACGCCTGGTATCTTTATAGTCCTGTCGGGTTTCGCCACCTCTGACTTGAGCGTCGATTTT
TGTGATGCTCGTCAGGGGGGCGGAGCCTATGGAAAAACGCCAGCAACGCGGCCTTTTTACGGTTCCTGGCCTTTTGC
TGGCCTTTTGCTCACATGTTCTTTCCTGCGTTATCCCCTGATTCTGTGGATAACCGTATTACCGCCTTTGAGTGAGC
TGATACCGCTCGCCGCAGCCGAACGACCGAGCGCAGCGAGTCAGTGAGCGAGGAAGCGGAAGAGCGCCCAATACGCA
AACCGCCTCTCCCCGCGCGTTGGCCGATTCATTAATGCAGCTGGCACGACAGGTTTCCCGACTGGAAAGCGGGCAGT
GAGCGCAACGCAATTAATGTGAGTTAGCTCACTCATTAGGCACCCCAGGCTTTACACTTTATGCTTCCGGCTCGTAT
GTTGTGTGGAATTGTGAGCGGATAACAATTTCACACAGGAAACAGCTATGACCATGATTACGCCAAGCGCGCAATTA
ACCCTCACTAAAGGGAACAAAAGCTGGAGCTGCAAGC

**J) pAVA3939 [TCTN3 minigene (EFS-6xMyc-TCTN3-Ex10-I10-Ex11-copGFP-3xFLAG)]**
**BGH polyA: 45-269**
**EFS: 2447-2658**
**6xMyc: 2724-2978**
**TCTN3 minigene: Ex10-I10-Ex11: 2991-3520**
**CopGFP: 3521-4201**
**3xFlag: 4214-4288**

tctatagtgtcacctaaatgctagagctcgctgatcagcctcgactgtgccttctagttgccagccatct
gttgtttgcccctcccccgtgccttccttgaccctggaaggtgccactcccactgtcctttcctaataaa
atgaggaaattgcatcgcattgtctgagtaggtgtcattctattctggggggtggggtggggcaggacag
caagggggaggattgggaagacaatagcaggcatgctggggatgcggtgggctctatggctCTGACCTAT
CGCGGTCGTCTTAATgcctggggtgcctaatgagtgagctaactcacattaattgcgttgcgctcactgc
ccgctttccagtcgggaaacctgtcgtgccagctgcattaatgaatcggccaacgcgcggggagaggcgg
tttgcgtattgggcgctcttccgcttcctcgctcactgactcgctgcgctcggtcgttcggctgcggcga
gcggtatcagctcactcaaaggcggtaatacggttatccacagaatcaggggataacgcaggaaagaaca
tgtgagcaaaaggccagcaaaaggccaggaaccgtaaaaaggccgcgttgctggcgtttttccataggct
ccgcccccctgacgagcatcacaaaaatcgacgctcaagtcagaggtggcgaaacccgacaggactataa
agataccaggcgtttccccctggaagctccctcgtgcgctctcctgttccgaccctgccgcttaccggat
acctgtccgcctttctcccttcgggaagcgtggcgctttctcaatgctcacgctgtaggtatctcagttc
ggtgtaggtcgttcgctccaagctgggctgtgtgcacgaaccccccgttcagcccgaccgctgcgcctta
tccggtaactatcgtcttgagtccaacccggtaagacacgacttatcgccactggcagcagccactggta
acaggattagcagagcgaggtatgtaggcggtgctacagagttcttgaagtggtggcctaactacggcta
cactagaaggacagtatttggtatctgcgctctgctgaagccagttaccttcggaaaaagagttggtagc
tcttgatccggcaaacaaaccaccgctggtagcggtggtttttttgtttgcaagcagcagattacgcgca
gaaaaaaaggatctcaagaagatcctttgatcttttctacggggtctgacgctcagtggaacgaaaactc
acgttaagggattttggtcatgagattatcaaaaaggatcttcacctagatccttttaaattaaaaatga
agttttaaatcaatctaaagtatatatgagtaaacttggtctgacagttaccaatgcttaatcagtgagg
cacctatctcagcgatctgtctatttcgttcatccatagttgcctgactccccgtcgtgtagataactac
gatacgggagggcttaccatctggccccagtgctgcaatgataccgcgagacccacgctcaccggctcca
gatttatcagcaataaaccagccagccggaagggccgagcgcagaagtggtcctgcaactttatccgcct
ccatccagtctattaattgttgccgggaagctagagtaagtagttcgccagttaatagtttgcgcaacgt
tgttgccattgctacaggcatcgtggtgtcacgctcgtcgtttggtatggcttcattcagctccggttcc
caacgatcaaggcgagttacatgatcccccatgttgtgcaaaaaagcggttagctccttcggtcctccga
tcgttgtcagaagtaagttggccgcagtgttatcactcatggttatggcagcactgcataattctcttac
tgtcatgccatccgtaagatgcttttctgtgactggtgagtactcaaccaagtcattctgagaatagtgt
atgcggcgaccgagttgctcttgcccggcgtcaatacgggataataccgcgccacatagcagaactttaa
aagtgctcatcattggaaaacgttcttcggggcgaaaactctcaaggatcttaccgctgttgagatccag
ttcgatgtaacccactcgtgcacccaactgatcttcagcatcttttactttcaccagcgtttctgggtga
gcaaaaacaggaaggcaaaatgccgcaaaaaagggaataagggcgacacggaaatgttgaatactcatac
tcttcctttttcaatattattgaagcatttatcagggttattgtctcatgagcggatacatatttgaatg
tatttagaaaaataaacaaataggggttccgcgcacatttccccgaaaagtgccacctgacgtcATTAAG
GATATTCACCATTATCGTCATTTAAATAGTCGCAAACGCGACTGTTCGGCTCCGGTGCCCGTCAGTGGGC
AGAGCGCACATCGCCCACAGTCCCCGAGAAGTTGGGGGGAGGGGTCGGCAATTGAACCGGTGCCTAGAGA
AGGTGGCGCGGGGTAAACTGGAAAGTGATGTCGTGTACTGGCTCCGCCTTTTTCCCGAGGGTGGGGGAG
AACCGTATATAAGTGCAGTAGTCGCCGTGAACGTTCTTTTTCGCAACGGGTTTGCCGCCAGAACACAGGT
GTCGTGACGCGaatacgactcactatagggagacccaagcttGGATCCCATCGATTTAAAGCTATGGAGC
AAAAGCTCATTTCTGAAGAGGACTTGAATGAAATGGAGCAAAAGCTCATTTCTGAAGAGGACTTGAATGA
AATGGAGCAAAAGCTCATTTCTGAAGAGGACTTGAATGAAATGGAGCAAAAGCTCATTTCTGAAGAGGAC
TTGAATGAAATGGAGCAAAAGCTCATTTCTGAAGAGGACTTGAATGAAATGGAGAGCTTGGGCGACCTCA
CCATGGAGCAAAAGCTCATTTCTGAAGAGGACTTGAATTTAAGATCTATGGCTTTTCAACAGAGCACAGC

TGCTTCTCTCACCAGTCCTAGAAGTGGGAATCCTGGCTATATAGTTGGGAAGCCACTCTTGGCTCTGACT
GATGATATAAGTTACTCAgtatccttttagagctgggtggcctgttgcagcctaatgagaaagctgca
aaggctttcaaacttatgttagactagccaaaggtgaggaatttttgttcattataatgaggtacttat
atactaatatatggttaatcattttggaatctagttgtctctctgtggttttctgaggaaaaaatcac
aaatttgtgactcgaacattatgatagtaatacaaaataaatagcattaaaggagaatgagaacataaat
tttgcaatgctgcctcctttttttttctgttatgatgtccttgatcatcacttttccaccttggATGACC
CTCTTACAGAGCCAGGGTAATGGAAGTTGCTCTGTTAAAAGACATGAAGTGCAGTTTGGAGTGAATGCAA
TATCTGGATGCAAGCTCAGG**ATG**CCCGCCATGAAGATCGAGTGCCGCATCACCGGCACCCTGAACGGCGT
GGAGTTCGAGCTGGTGGGCGGCGGAGAGGGCACCCCCGAGCAGGGCCGCATGA**CCAACAAGATGAAGAGC**
**ACCAAAG**GCGCCCTGACCTTCAGCCCCTACCTGCTGAGCCACGTGATGGGC**TA**CGGCTTCTACCACTTCG
GCACCTACCCCAGCGGCTACGAGAACCCCTTCCTGCACGCCATCAACAACGGCGGCTACA**CCAACACCCG**
**CATCGAGAAGTACG**AGGACGGCGGCGTGCTGCAC**GTGAGCTTCAGCTACCGCTACGAG**GCCGGCCGCGTG
ATCGGCGACTTCAAGGTGGTGGGCACCGGCTTCCCCGAGGACAGCGTGAT**CTTCACCGACAAGATCATCC**
**GCAGCAACG**CCACCGTGGAGCACCTGCACCCCATGGGCGATAACGTGCTGGTGGGCAGCTTCGCCCGCAC
CTTCAGCCTGCGCGACGGCGGCTACTACAGCTTCGTGGTGGACAGCCACATGCACTTCAAGAGCGCCATC
CACCCCAGCATCCTGCAGAACGGGGGCCCCATGTTCGCCTTCCGCCGCGTGGAGGAGCTGCACAGCAACA
CCGAGCTGGG**CATCGTGGAGTACCAGCACGCCTTC**AAGACCCCCATCGCCTTCGCC**AGATCTCGAGCTC**G
Aatcgatgtcgaggattacaaggatgacgacgataaggactataaggacgatgatgacaaggactacaaa
gatgatgacgataaatagggcccggtacc

Predicted protein sequence of TCTN3 minigene with spliced minor intron 10:
MEQKLISEEDLNEMEQKLISEEDLNEMEQKLISEEDLNEMEQKLISEEDLNEMEQKLISEEDLNEMESLGDLTMEQK
LISEEDLNLRS**MAFQQSTAASLTSPRSGNPGYIVGKPLLALTDDISYSMTLLQSQGNGSCSVKRHEVQFGVNAISGC**
**KLR**MPAMKIECRITGTLNGVEFELVGGGEGTPEQGRMTNKMKSTKGALTFSPYLLSHVMGYGFYHFGTYPSGYENPF
LHAINNGGYTNTRIEKYEDGGVLHVSFSYRYEAGRVIGDFKVVGTGFPEDSVIFTDKIIRSNATVEHLHPMGDNVLV
GSFARTFSLRDGGYYSFVVDSHMHFKSAIHPSILQNGGPMFAFRRVEELHSNTELGIVEYQHAFKTPIAFARSRARI
DVEDYKDDDDKDYKDDDDKDYKDDDDK-
