## Supplementary figures and images for "Identification of Human Pathways Acting on Nuclear Non-Coding RNAs Using the Mirror Forward Genetic Approach"

### Supplemental Data S6

#### Supplemental Data S6

Schematics of the employed FACS gates

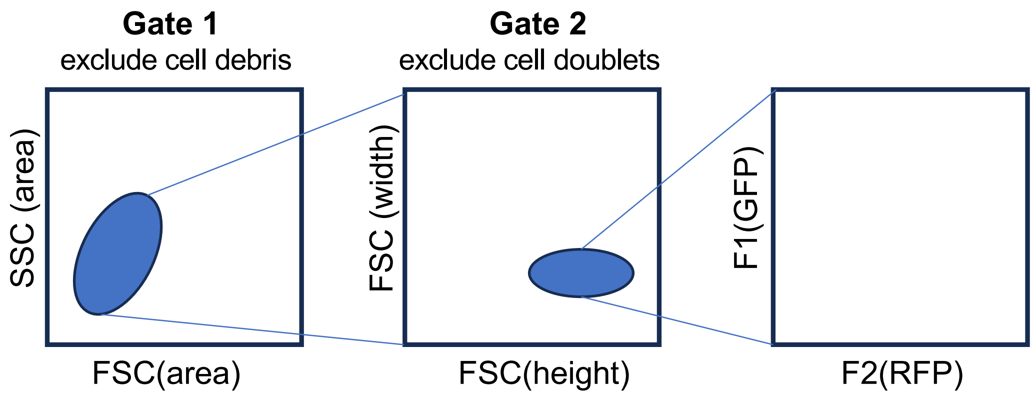
